# Microenvironment Impacts the Molecular Architecture and Interactivity of Resident Cells in Marmoset Brain

**DOI:** 10.1101/2021.01.25.426385

**Authors:** Jing-Ping Lin, Hannah M. Kelly, Yeajin Song, Riki Kawaguchi, Daniel H. Geschwind, Steven Jacobson, Daniel S. Reich

## Abstract

The microenvironments of the brain consist of specialized cell types that together influence physiological functions in health and pathological outcomes in disease. Despite apparent differences in the density of neurons and oligodendrocytes in various milieus, such as gray matter (GM) and white matter (WM), the extent of structural and functional heterogeneity of other resident cells remains unclear. We profiled RNA in ~500,000 nuclei from 19 tissue types across the central nervous system of the healthy adult common marmoset (*Callithrix jacchus*) and mapped 87 identified subclusters (including neurons, glia, and vasculature) spatially onto a 3D MRI atlas. We performed cross-species comparison, explored regulatory pathways, surveyed cellular determinants of neurological disorders, and modeled regional intercellular communication. We found spatially segregated microglia, oligodendrocyte lineage cells, and astrocytes in WM and GM. WM-glia are diverse, are enriched with genes involved in stimulus response and biomolecule modification, and interact with other resident cells more extensively than their GM counterparts. GM-glia preserve the expression of developmental morphogens into adulthood and share 6 differentially enriched transcription factors that restrict the transcriptome complexity. Our work in marmoset, an experimentally tractable animal model with >5 times more WM volume and complexity than mouse, identifies novel WM-glia subtypes and their contributions to different neurological disorders. A companion *Callithrix jacchus* Primate Cell Atlas (CjPCA) is available through an online portal https://cjpca.ninds.nih.gov to facilitate data exploration.

## Main

An understanding of microenvironmental heterogeneity and its wide impact on biological processes is necessary for the interpretation of experimental perturbations. Recent advances in genetic profiling tools with single-cell resolution have revealed that the extent of regional cellular diversity in the brain’s gray matter is far beyond what had traditionally been appreciated (Bakken et al., 2020; Hodge et al., 2020; Zeisel et al., 2018). However, characterization of cellular profiles in white matter (particularly subcortical white matter) is limited due to its modest representation in mouse, the most common mammalian model system in biology. In the common marmoset (*Callithrix jacchus*), an emerging animal model that bridges mouse and higher primates behaviorally, genetically, immunologically, and neuroanatomically, we surveyed the diversity of single cells throughout the brain, including both gray and white matter. Leveraging the massively greater (>5-fold more) subcortical white matter to cortical gray matter volumetric ratio in marmoset compared to mouse (Ventura-Antunes et al., 2013), we found unexpected and previously undescribed glial heterogeneity in WM. We conclude, based on comprehensive analysis of gene-expression patterns and regulatory pathways, that WM glia *accrued* additional features, are further *advanced* in the program of specialization, *forgot* their morphogenic origin, and are more *talkative* than their GM counterparts.

### Glia reflect differential residence in gray and white matter

We performed in vivo magnetic resonance imaging (MRI) of the brains of two marmosets and identified brain structures by cross-referencing to 3D MRI atlases (Figure 1A). For each animal, we made a customized brain holder to facilitate image-guided sampling from a confined region (corresponding to ~10 μL of tissue, Figure 1B, S1), isolated nuclei from 22 locations (Figure 1C), and categorized them in three different ways (Figure 1D). We surveyed cells without preselection, integrating all 42 samples. A total of 534,553 nuclei were recovered after a first round of preprocessing and quality control (Figure S2 – 4 and Table S1). In the Level 1 analysis, we divided nuclei into 6 classes, as determined by the expression of canonical markers (NEU, *CNTN5*^*+*^ neurons; OLI, *MBP*^*+*^ oligodendrocytes; AST, *ALDH1L1*^*+*^ astrocytes; OPC, *PDGFRA*^*+*^ oligodendrocyte progenitor cells; MIC, *PTPRC*^*+*^ microglia/immune cells; VAS, *LEPR*^*+*^ vascular cells / *CEMIP*^*+*^ meningeal cells / *TMEM232*^*+*^ ventricular cells) (Figure S5A). The relative composition of marmoset cell types (47% neurons, 35% oligodendrocyte-lineage cells, 12% astrocytes, and 4% immune cells; Figure 1E) across our selected brain regions corresponds well to a morphological counting of cell types in human neocortex across age (18 – 93 years) and sex (42% neurons, 43% oligodendrocyte-lineage cells, 11% astrocytes, and 3% immune cells) (Bartheld et al., 2016; Pelvig et al., 2008).

**Figure 1.**
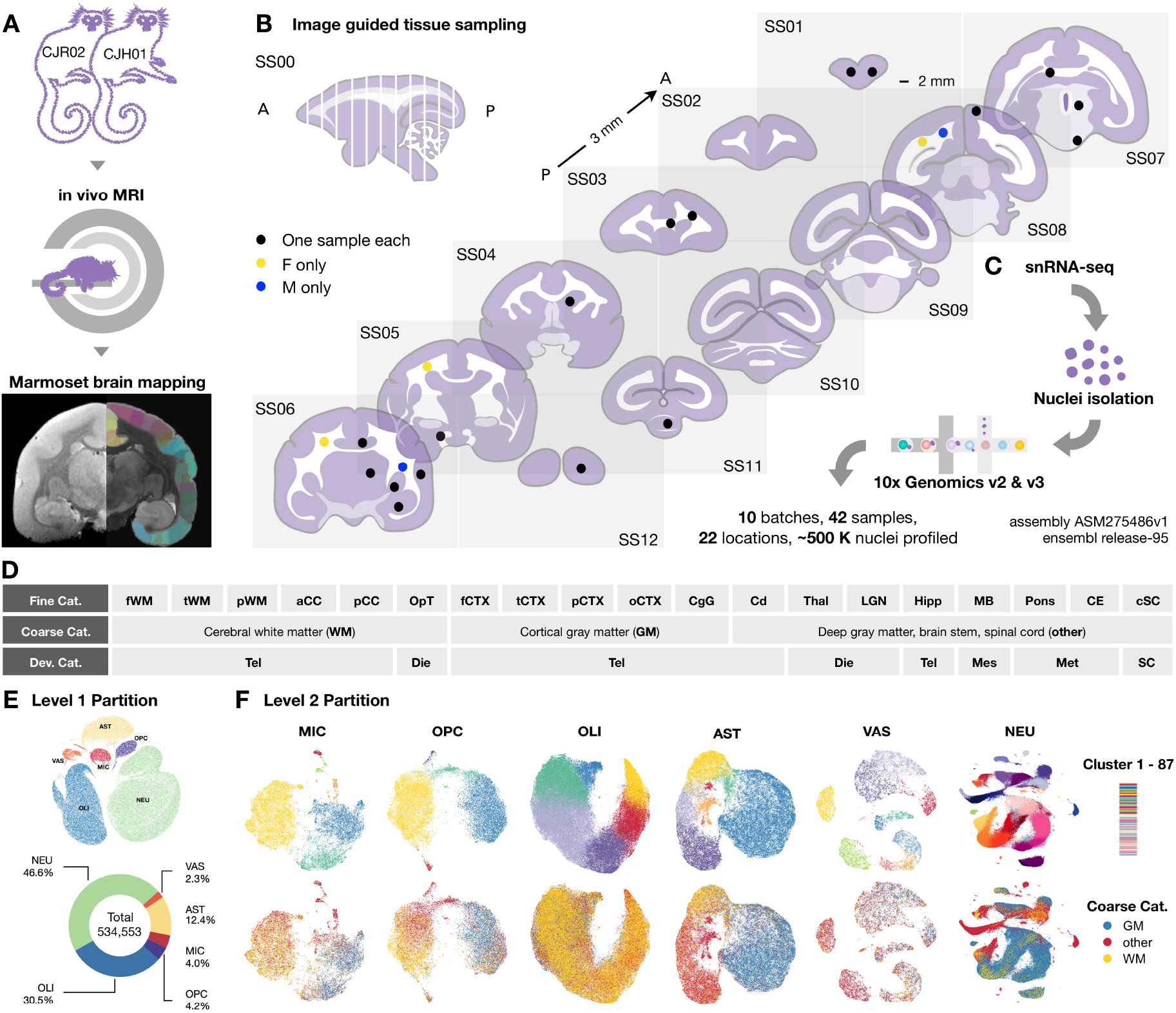
Single-nucleus transcriptional profiling of adult marmoset brain and spinal cord identifies 6 cell classes and 87 subclusters, with key differences in glial phenotypes in gray vs. white matter. (A) Schematic illustration of experimental workflow to scan and map brain to marmoset MRI atlases (Marmoset Brain Mapping V1 & V2). (B) Location of tissue samples on the standard slab (SS) index. Samples from 22 different areas were collected as cylinders of 2 mm diameter by 3 mm height. (C) Nuclei were isolated to prepare cDNA library with 10x Genomics technology and sequenced to profile transcriptomes. Reads were mapped to a marmoset genome assembly (ASM275486v1) with Ensembl release-95 for annotation. In Level 1 clustering, 534,553 nuclei were included. (D) Total sampled areas are labeled by three types of tissue categories (Cat): fine, coarse, and developmental (Dev). **fWM**, frontal white matter; **tWM**, temporal white matter; **pWM**, parietal white matter; **aCC**, anterior corpus callosum; **pCC**, posterior corpus callosum; **OpT**, optic tract; **fCTX**, frontal cortex; **tCTX**, temporal cortex; **pCTX**, parietal cortex; **oCTX**, occipital cortex; **CgG**, cingulate gyrus; **Cd**, caudate; **Thal**, thalamus; **LGN**, lateral geniculate nucleus; **Hipp**, hippocampus; **MB**, midbrain; **Pons**, Pons; **CE**, cerebellum; **cSC**, cervical spinal cord; **Tel**, telencephalon; **Die**, Diencephalon; **Mes**, Mesencephalon; **Met**, Metencephalon; **SC**, Spinal cord. (E) Cells were initially divided into 6 major classes (Level 1 clustering), comprising neurons (NEU), oligodendrocytes (OLI), oligodendrocyte progenitor cells (OPC), microglia/immune cells (MIC), astrocytes (AST), and vascular/meningeal/ventricular cells (VAS). The number and percentage of nuclei that passed the first round of standardized quality control ranged from 46.6% (NEU) to 2.3% (VAS). To avoid overcrowding, the scatter plots presented here represent a random sample of 100K nuclei. (F) Top, each of the Level 1 classes was further subclustered and colored by Level 2 clustering. Bottom, the same UMAP scatter plots analyzed in Level 2 are colored by coarse tissue category. GM-enriched and WM-enriched microglia, OPC, and astrocytes subtypes were discovered. See also Figure S1–S12, Table S1, S2, and S4.

The Level 2 analysis consisted of additional rounds of quality control and manifold learning (Methods), in which the 6 major cell classes were further grouped into 87 subclusters (Figure 1F). We mapped the general landscape of the dataset (Figure S3D – E, S5 – 7) and inspected the major features related to neurons (Figure S8 – 12), finding close agreement with several previous reports (Bakken et al., 2020; Hodge et al., 2020; Krienen et al., 2020). Interestingly, we found apparent spatial segregation of microglia, oligodendrocyte progenitor cells (OPC), and astrocytes between cerebral white matter (hereafter denoted “**WM**”) and cortical gray matter (hereafter denoted “**GM**”). We therefore leveraged the unprecedented level of regional resolution to assess glial heterogeneity across the brain.

### White matter microglia appear to be older than gray matter microglia

In the microglia/immune cell class (MIC), a total of 18,279 nuclei were included in the Level 2 analysis (Figure S13). We found 7 distinct subclusters, of which 4 were circulating peripheral immune populations and 3 were brain-resident immune cells (microglia). In additional to canonical markers (*P2RY13*, *ITGAX*), the expression of *FLT1* (vascular endothelial growth factor receptor 1) diverged substantially across immune cells (Figure 2C), such that circulating peripheral blood mononuclear cells (PBMC) were *FLT1*^−^ and microglia were *FLT1*^+^.

**Figure 2.**
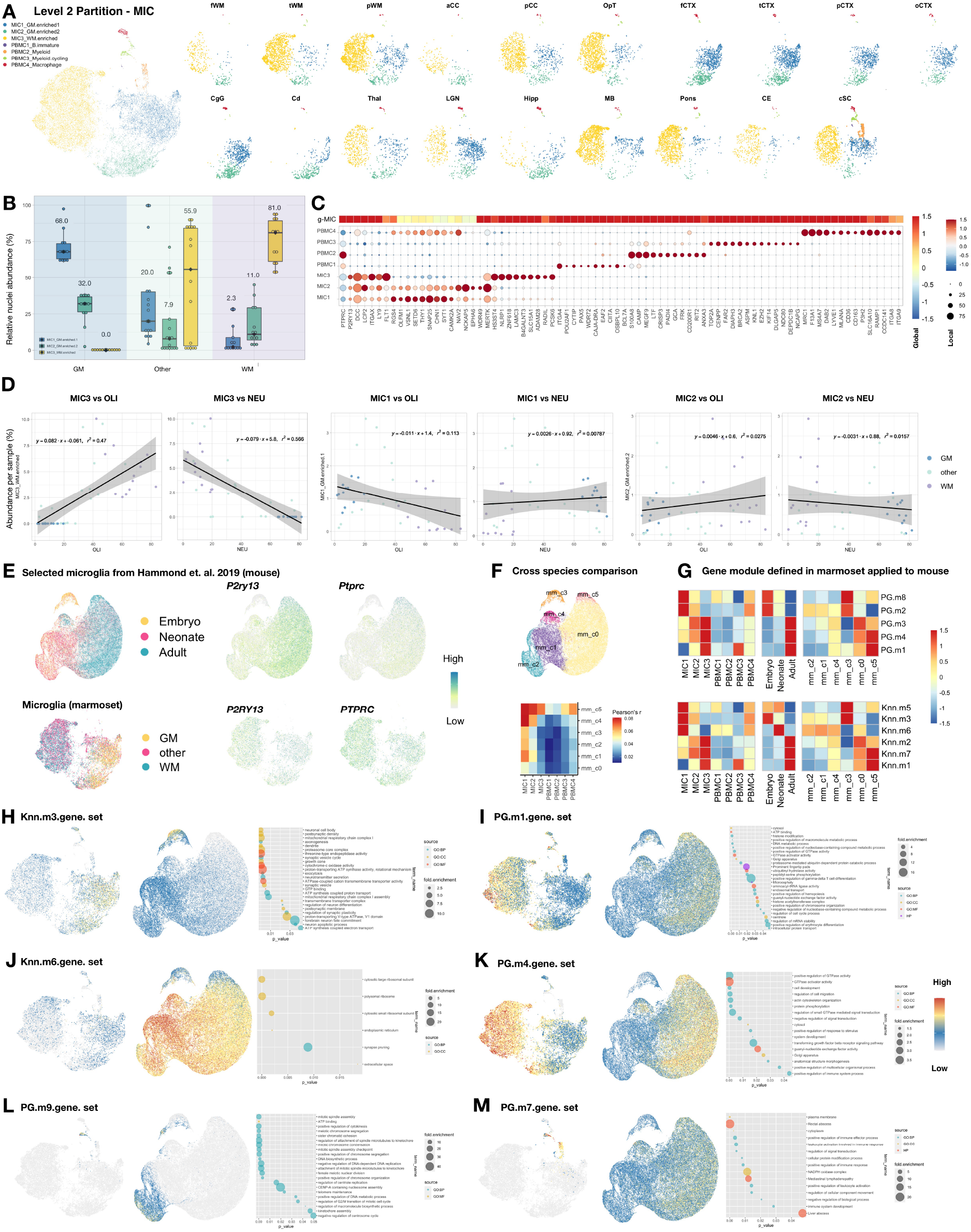
Microglia subclusters are spatially segregated, the transcriptomic specialization of marmoset GM-microglia resembles that in microglia of young mice, and pathways related to stimulus response are elevated in WM-microglia. (A) UMAP scatter plot visualization of microglia/immune cells (MIC) colored by subcluster and split by sampling site. See Figure 1D for tissue label abbreviation list. (B) Box plot showing the relative abundance of MIC subclusters in each tissue type in coarse category (see Figure 1D for list). MIC1 and MIC2 are enriched in GM and MIC3 in WM. Median is annotated (◆) and listed. (C) Heatmap and dot plot showing mean-centered and z-score-scaled marker gene expression in global space (heatmap, MIC average expression among Level 1 classes) and local space (dot plot, among Level 2 subclusters). The portion of the heatmap that represents global microglia/immune cells (g-MIC) is reproduced here from the full heatmap shown in Figure S13E. (D) Linear regression on the normalized abundance of MIC1, MIC2, and MIC3 subclusters with oligodendrocyte (OLI) or neuron (NEU) clusters. The abundance of WM-enriched microglia (MIC3) corelates with the abundance of OLI and NEU. The density of gray matter-enriched microglia (MIC1 and MIC2) is independent of tissue type. (E) Top, UMAP scatter plot visualization of mouse microglia from Hammond et. al. 2019 colored by age and expression of *P2ry13* and *Ptprc* (See Figure 14 for detail). Bottom, UMAP scatter plot visualization of marmoset microglia colored by tissue type in coarse category and expression of *P2RY13* and *PTPRC*. (F) Top, UMAP scatter plot visualization of mouse microglia from Hammond et. al. 2019 colored by subclusters (See Figure 14 for detail). Bottom, heatmap showing Pearson’s correlation coefficient (r) between 7 MIC subclusters from marmoset and 6 microglia subclusters from mouse (mm_c0 – mm_c5). (G) Heatmap showing the expression of gene modules in 7 MIC subclusters from marmoset (left), mouse microglia grouped by age (middle), and 6 mouse microglia subclusters (right). Gene modules that are highly enriched in gray matter microglia (MIC1) of marmoset are enriched in microglia of younger mouse (PG.m8/2, Knn.m5/3/6), and gene modules that are enriched in white matter microglia (MIC3) of marmoset are enriched in microglia of older mouse (PG.m3/4/1, Knn.m2/7/1). (H – M) UMAP scatter plot visualization of marmoset MIC (left) and mouse microglia (middle) colored by averaged expression of MIC gene modules. Right, dot plot showing the enriched GO terms from the list of genes in each module. GO terms relating to pre- and post-synaptic assembly, neurotransmitter secretion, and regulation of neuron development were enriched in GM-microglia (Knn.m3, H). Terms related to biomolecule activity, cell movement, stimulus response, and regulation of nucleic acids were enriched in WM-microglia (PG.m1/4, I and K). **GO**, gene ontology; **BP**, biological process; **CC**, cellular component; **MF**, molecular function; **HP**, human phenotype ontology. See also Figure S13–14, Table S2 and S4.

Within the 3 clusters of microglia, we noticed a spatial separation in subtypes across brain regions. MIC1 and MIC2 were enriched in GM and MIC3 in WM (Figure 2A – B). This relationship was so strong that the abundance of WM-enriched (MIC3) microglia was positively and negatively correlated with the number of oligodendrocytes (OLI) and neurons (NEU), respectively. In contrast, GM-enriched populations (MIC1 and MIC2) had similar densities across brain regions (Figure 2D). We further investigated the enrichment of genes in GM- and WM-enriched microglia using gene module analysis (Cao et al., 2019), grouping genes into lists by their expression similarity (Methods), and performed gene ontology (GO) analysis (Figure 2E – M and Table S2). The expression of a given module is visualized by aggregating the expression of multiple genes in the list (Figure S13C), and cross-referenced with its expression pattern on microglia isolated from the whole brain of mice by translating marmoset gene names to mouse (Hammond et al., 2019). Specifically, we clustered mouse microglia with our pipeline and grouped clusters by age (embryo, neonate, adult; Figure 2E – F, S14A – F). Interestingly, we found that gene modules (PG.m8/2, Knn.m5/3/6) enriched in marmoset GM-microglia were highly expressed in young mouse microglia (Figure 2G, 2H, S14G – H). In contrast, adult mouse microglia showed higher expression of gene modules (PG.m3/4/1, Knn.m2/7/1) also enriched in marmoset WM-microglia (Figure 2G, 2I, 2K, S14G – H). Together, these findings suggest that the transcriptome profile of GM-microglia is more naïve than WM-microglia.

Illustrating the ability of this workflow to highlight relevant biological processes, GO terms like “positive regulation of leukocyte activation,” “positive regulation of immune responses,” and “immune system development” were associated with PBMC (PG.m7 genes, Figure 2M). Cell cycle features like “DNA biosynthesis process” and “regulation of G2/M transition of mitotic cell cycle” were discovered in cycling myeloid cells (PG.m9, Figure 2L). On the other hand, genes related to “synapse pruning” were uniformly detected in all microglia subclusters (Knn.m6, Figure 2J) (Wilton et al., 2019). In GM-microglia, we found enrichment of GO terms relating to pre- and post-synaptic assembly, neurotransmitter secretion, and regulation of neuron development (Knn.m3, Figure 2H). Terms related to bio-molecule activity, cell movement, stimulus response, and regulation of nucleic acids were enriched in WM-microglia (PG.m1/4, Figure 2I, 2K). These findings suggest that, in homeostasis, GM-microglia are relatively stationary and more involved in the modulating neuronal synaptic activity, whereas WM-microglia are primed to a more active, migratory state.

### White matter oligodendrocyte progenitor cells form a unique population

A similar GM-WM segregation was noted for OPC populations, grouped into 5 subclusters from a total of 20,306 nuclei (Figures 3A–B, S15–16). WM-enriched OPC (OPC3) were positively correlated with the abundance of oligodendrocytes and negatively with the abundance of neurons, whereas GM-enriched OPC (OPC1) were similar in density regardless of sampling site (Figure 3D). Interestingly, several top differentially expressed gene (DEG) related to general neuronal functioning (*RGS4*, *OLFM1*, *VSNL1*, *SETD6*, *THY1*, *SNAP25*, *CHN1*, *CAMK2A*; Figure 2B) were shared between GM-OPC and GM-microglia (OPC1 and MIC1), and both OPC1 and MIC1 had fewer detected genes compared to their WM counterparts (OPC3 and MIC3; Figures S13B and S15B). We therefore hypothesize that GM-microglia and GM-OPC are in a more naïve state relative to WM-microglia and WM-OPC, which in turn have acquired gene expression features specific to their microenvironment.

**Figure 3.**
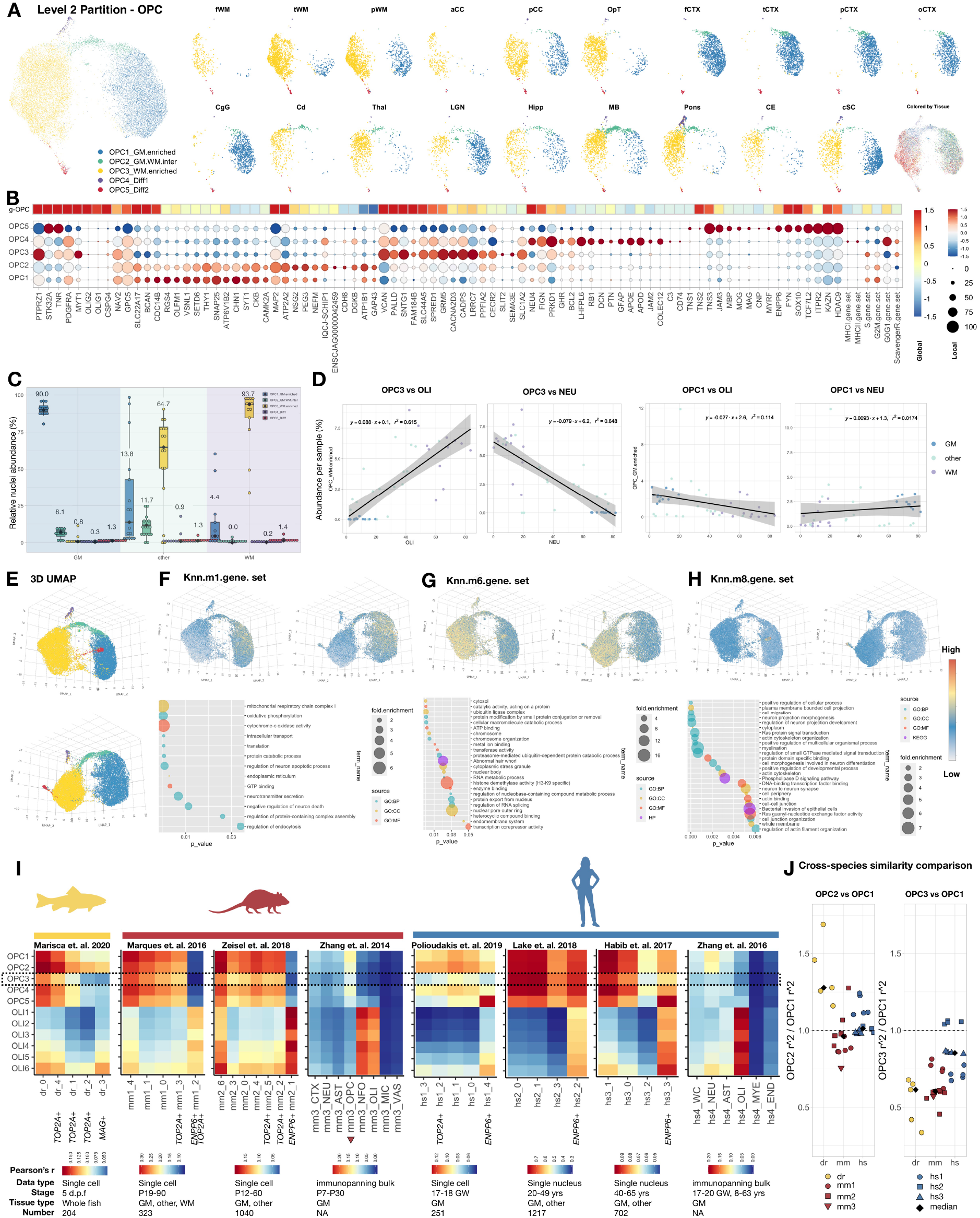
Oligodendrocyte progenitor cell (OPC) subclusters are spatially segregated, GM-OPC are enriched with pathways related to neuronal support, and WM-OPC are a unique population compared to previously reported OPC in other species. (A) UMAP scatter-plot visualization of oligodendrocyte precursor cells (OPC) colored by Level 2 subclustering and split by sampling site. See Figure 1 for tissue label abbreviation. (B) Heatmap and dot plot showing marker gene expression in global space (heatmap, OPC average expression among Level 1 classes) and local space (dot plot, Level 2 subclusters). See Table S4 for gene set list. The portion of the heatmap that represents global OPC (g-OPC) is reproduced here from the full heatmap shown in Figure S15F. (C) Box plot showing the relative abundance of OPC subclusters of each tissue type in coarse category (see Figure 1D for list). Median is annotated (◆) and listed. (D) Linear regression on the normalized abundance of Level 2 subclusters (OPC1 and OPC3) with OLI or NEU partition annotated in Level 1 analysis. Individual samples were colored by coarse category. The abundance of WM-enriched OPC (OPC3) corelates with the abundance of OLI and NEU. The abundance of GM-enriched OPC (OPC1) is proportional across different tissue type. (E – H) Two angles of 3D UMAP scatter-plot visualizations of nuclei colored by OPC subcluster annotation (E) and average expression of selected gene modules (Top, F–H). Dot plot showing the enriched GO terms from the list of genes in each module (Bottom, F–H). (I) Heatmap showing the Pearson’s correlation coefficient (r) between 11 oligodendrocyte lineage cells (OPC1-5 and OLI1-6, see Figure 4 for detail) from marmoset and multiple subclusters of oligodendrocyte-lineage cells in zebrafish (dr), mouse (mm), and human (hs). **d.p.f.**, days post fertilization; **P**, postnatal day; **GW**, gestational weeks. (J) Scatter plot showing the ratio of r^2^ between OPC2 and OPC1 (left) and OPC3 and OPC1 (right), calculated from the correlation analysis across species shown in Figure 3I. OPC1 and OPC2 are similar to one another and to OPC found in other species. On the other hand, WM-OPC (OPC3) is a distinct subcluster, consistently showing lower correlation with previously defined clusters compared to GM-OPC (OPC1). Median is annotated (◆). See also Figure S15–S19, Table S2 and S4

To further explore this notion, we performed gene module analysis (Figure S15C, Table S2) as described above. As a positive control, “myelination” term (Knn.m8; Figure 3H) is enriched in OPC5, which had transcriptomic evidence of differentiation to oligodendrocytes (*ENPP6*, *CNP*, *MBP*) but preserved expression of a canonical OPC marker (*PDGFRA*). We found regulation of synapse, neurotransmitter secretion, and regulation of neuron survival to be enriched in OPC1 (Knn.m1; Figure 3F), GO terms that are very similar to those enriched in MIC1 (PG.m8; Figure 2H). On the other hand, WM-OPC (OPC3) were enriched with terms related to component organization, molecule modification, and cell motility (Knn.m6 and PG.m5; Figure 3G, S15D). Markers enriched in OPC3 are known for regulating OPC dispersal (*SLIT2*) (Liu et al., 2012) and inhibiting CNS angiogenesis (*SEMA3E*) (Mecollari et al., 2014) (Figure 3B, S13D). Together, these observations support our hypothesis that adult WM-OPC, in homeostasis, are a population set a to more active, migratory state compare to their GM counterpart.

To understand how OPC subclusters we identified in marmoset compare to OPC in other species (Habib et al., 2017; Lake et al., 2018; Marisca et al., 2020; Marques et al., 2016; Polioudakis et al., 2019; Zeisel et al., 2018; Zhang et al., 2014; 2016), we performed a Pearson’s correlation analysis (Figure S17–19). Prior to comparison, we humanized gene names of each species with a one-to-one ortholog index based on BioMart information (Methods). As a positive control, we found agreement in oligodendrocyte lineage differentiation features across species (Figure 3I): marmoset differentiating OPC (OPC5) and oligodendrocytes (OLI1–6) correlate with *ENPP6*^+^/*MAG*^+^ oligodendrocyte lineage cells in zebrafish (dr_3), mouse (mm1_2, mm2_1, mm3_NFO, mm3_OLI), and human (hs1_4, hs2_2, hs3_3, hs4_OLI). Overall, we noted greater heterogeneity of OPC subpopulations in marmoset, such that there was higher correlation between OPC in all other species and marmoset GM-OPC (OPC1) than marmoset WM-OPC (OPC3). We further quantified this observation by comparing the fold-change of similarity between OPC subclusters (ratio of r^2^ values) (Figure 3J). For OPC3, the median of the ratio is consistently lower than 1 across datasets, indicating that OPC3 is less similar than OPC1 to previously described OPC in other species. As a negative control, the similarity ratio between OPC2 (transcriptomically more similar to OPC1 than OPC3, Figure 3B) and OPC1 is centered at 1, with the exception of zebrafish OPC. These comparisons are limited by differences in methodology (including full-length SMART-Seq versus droplet methods, sequencing of single cells vs. nuclei, use of adult vs. prenatal tissues). More robust cross-species comparison is therefore needed, but it is likely that the marmoset WM-OPC-like population was missed in other datasets is in part due to insufficient WM sampling and low numbers of sequenced OPC rather than a specific marmoset innovation.

### Cortical gray matter contains relatively few young oligodendrocytes

A total of 128,710 nuclei were included in the marmoset OLI class, from which 6 subclusters were identified (Figure S20). Marmoset oligodendrocytes were arranged into a continuous, graded pattern in 2D UMAP space (Figure 4A), similar to what was observed in mouse brain under two different sampling paradigms (Marques et al., 2016; Zeisel et al., 2018) (Figure S22–23). We denoted as OLI1 the subcluster that appeared transcriptionally youngest based on a mouse study of differentiation-committed oligodendrocyte precursors, which were *Pdgfra*^-^ and *Tns3*^+^ (Marques et al., 2016), and named the other OLI subclusters sequentially. Instead of a clear GM-WM segregation as observed for microglia and OPC (Figure 2A and 3A), we found proportional differences along the intermingled oligodendrocyte subtypes across brain regions. OLI1 was lowest in GM (median abundance ~0.5%), compared to ~10% relative abundance in the WM and “other” sampling sites (Figure 4C). Similarly, OLI2 and OLI3 were relatively sparse in GM.

**Figure 4.**
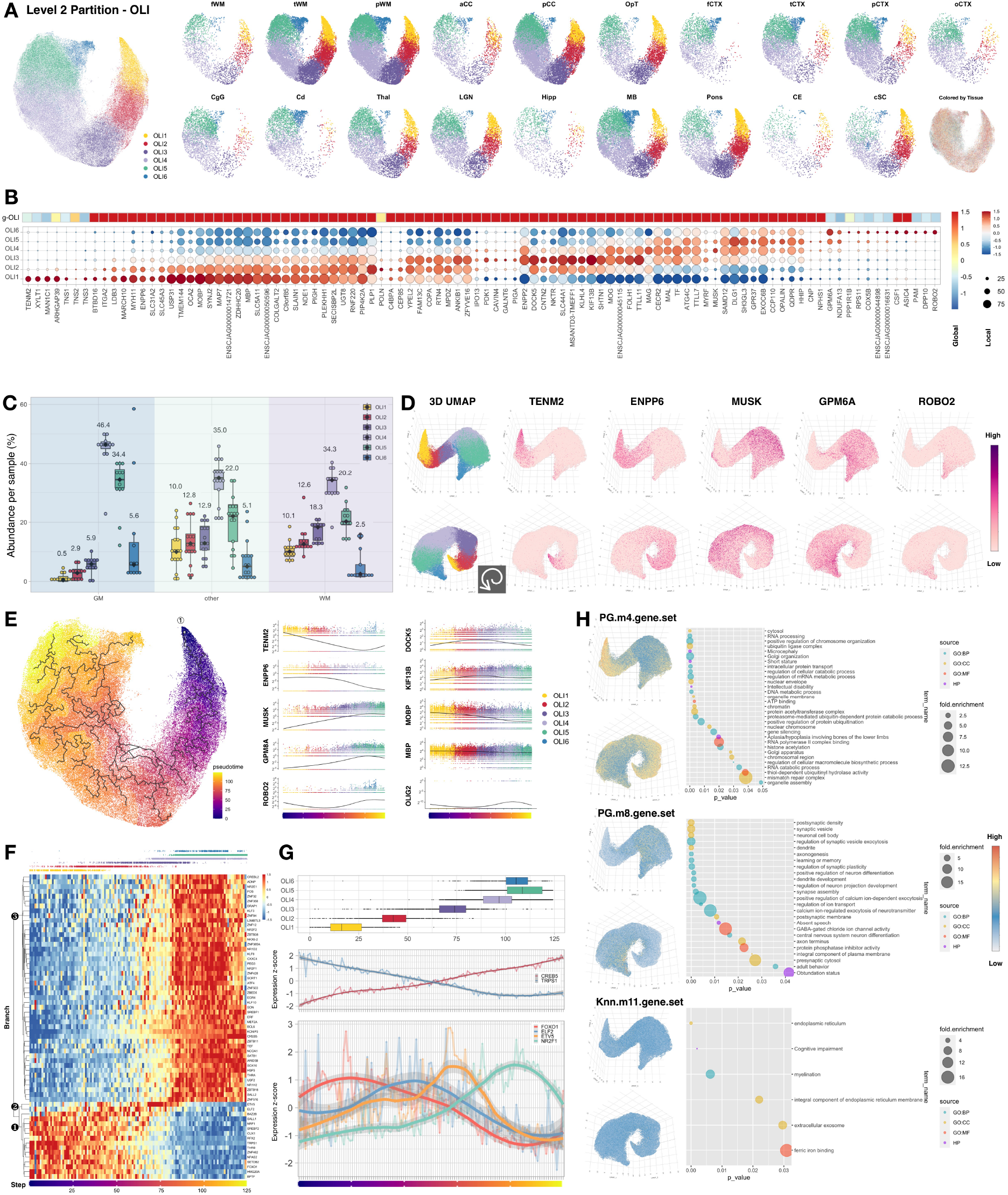
Oligodendrocyte differentiation state varies by region, oligodendrocytes in cortical gray matter are mostly mature, and sets of transcription factors are elevated in an orderly fashion along the developmental trajectory in pseudotime. (A) 2D UMAP scatter-plot visualization of oligodendrocytes (OLI) colored by Level 2 subcluster and split by sampling site. See Figure 1 for tissue label abbreviation list and Figure 5 for color code of each tissue. (B) Heatmap and dot plot showing mean-centered and z-score-scaled marker gene expression in global space (heatmap, OLI average expression among Level 1 classes) and local space (dot plot, Level 2 OLI subclusters). The portion of the heatmap that represents global OLI (g-OLI) is reproduced here from the full heatmap shown in Figure S20F. (C) Box plot showing the relative abundance of OLI subclusters in each tissue type (coarse category; see Figure 1D for list). Putative early (young) populations (OLI1 and OLI2, *ENPP6*^high^) are low in GM compared to WM and other tissue types. OLI4 is the major cluster in all tissue types. Median is annotated (◆) and listed. (D) Two angles of 3D UMAP scatter plot visualization of nuclei colored by OLI subcluster annotation and expression of selected marker genes. A spiral-like 3D UMAP pattern along the subclusters was noted. (E) Left, 2D UMAP scatter plot visualization of nuclei colored by pseudotime. The starting point of pseudotime was defined at the tip of the OLI1 subcluster (① denoted). Right, plot of gene expression as a function of pseudotime. Dots in each subplot are colored by OLI subcluster identity. (F) Heatmap showing the mean-centered and z-score-scaled expression of transcription factors across pseudotime. Nuclei were grouped into 125 bins (columns) by step size of 1 pseudo-time unit. Jitter plot on the top of heatmap is colored by OLI subcluster identity, showing the distribution of nuclei as a function of pseudotime for visual reference. Three major branches of genes were annotated to facilitate indexing (❶, ❷, and ❸). (G) Top, box plot showing the distribution of nuclei in each OLI subcluster across pseudotime bins. Median is annotated (–). Middle, line chart showing the expression of 2 representative transcription factors with linearly decreasing (*TRPS1*, branch ❶ in F) and increasing (*CREB5*, branch ❸ in F) expression pattern over the trajectory of pseudotime. Bottom, line chart showing the expression of 4 representative transcription factors with expression levels that peak at various pseudotime points. A locally estimated scatterplot smoothing (LOESS) curve was fitted for each expression pattern, and the flanking gray bands indicate 95% confidence intervals. (H) Two angles of 3D UMAP scatter plot visualization of nuclei colored by averaged expression of 3 OLI gene modules (PG.m4, PG.m8, and Knn.m11). Dot plot showing the enriched GO terms from the list of genes in each module. See also Figure S20-23, Table S2 and S4.

Interestingly, OLI1 (putatively youngest) and OLI6 (putatively oldest) are close together in the 2D UMAP projection, suggesting transcriptomic similarity. We therefore pursued a 3D UMAP analysis of oligodendrocytes, finding a spiral pattern that we also observed upon reanalysis of previously reported human (Jäkel et al., 2019) (Figure S21) and mouse (Zeisel et al., 2018)(Figure S22) oligodendrocyte transcriptomes. This spiral pattern was also captured at the level of DEG across oligodendrocyte subclusters (Figure 4D). The expression patterns of most oligodendrocyte DEG (*XYLT1*, *TNS1*, *TNS3*, *MAN1C1*, *BTBD16*, *CCP110*, *CSF1*, *DOCK5*, *PAM*, *MUSK*, *GPM6A*, *DPP10*) were aligned across species and were therefore used to label the gross developmental trajectory of oligodendrocytes (Figure 4B, S21–22).

We asked whether graded transcriptomic changes along the spiral oligodendrocyte trajectory can be modeled by waves of influence from within and/or directional stimuli from the environment. To address this, we performed a pseudotime analysis of marmoset oligodendrocytes and mapped the expression of transcription factors along the pseudotime trajectories. We transferred the precalculated UMAP embeddings and used Monocle3 to learn the principal graph (Methods). We set *ENPP6*^high^ oligodendrocytes as the starting point (Figure 4E, left) for this analysis and visualized gene expression dynamics along the pseudotime axis by overlaying the contribution from each OLI subcluster (Figure 4E, right). The pattern of expression dynamics agrees with the visual impression of gene expression along the spiral 3D UMAP path. In 3D, the molecular distances from OLI4 to OLI5 and from OLI4 to OLI6 are similar, indicating that OLI5 and OLI6 might develop in parallel, rather than dependently (Figure 4F and 4G, top).

We next extracted the calculated pseudotime values and arbitrarily binned them into 125 steps to map the expression pattern of a list of transcription factors along the trajectory. Interestingly, we found that expression of these 63 master regulators could be coarsely grouped into 3 sets (Steps 0–60, 60–80, and 80–125), each of which was largely distinct and used by different subsets of OLI along the pseudotime trajectory (Figure 4F). Set 1 was mainly present in OLI1 and OLI2 (nuclei in Steps 0–60), with high expression of transcription factors in Branch 1 (❶) and low expression in Branch 3 (❸). Set 3 was mainly present in OLI4–6 (nuclei in Steps 80–125), with high expression levels of Branch 3 transcription factors and low levels of Branch 1. Interestingly, an intermediate state (nuclei in Steps 60–80) was mainly observed in OLI3, with intermediate levels of both Branch 1 and 3 genes but high *ETV5* expression (Branch 2, ❷). The dynamic expression levels of selected transcription factors that fluctuated along the trajectory were plotted as a function of pseudotime (Figure 4G). Of all genes examined, only *ELF2* and *ETV5* peaked in the middle stages of oligodendrocytes (OLI2 and OLI3, respectively), whereas the other master regulators were clustered either at the early (OLI1) or late (OLI4–6) stages. A positive correlation between *ELF2* and myelin was supported in a human snRNA-seq study, in which *ELF2* was high in control WM, normal appearing WM, and remyelinated multiple sclerosis lesions but lower in WM lesions (active, chronic active, and chronic inactive) (Jäkel et al., 2019). On the other hand, *Etv5* can act as a suppressor of oligodendrocyte differentiation, such that enforced expression of *Etv5* in rat OPC decreased the production of MBP^+^ oligodendrocytes (Wang et al., 2017). That *ETV5* expression peaks in OLI3 (Step 60–80) suggests that OLI3 might be a population that is poised to differentiate (Figure 4G).

To further understand the functional significance of OLI subclusters, we performed gene module and GO analysis as described above (Figure S20, Table S2). Not surprisingly, we found that oligodendrocytes are involved in “myelination” (Knn.m11; Figure 4H) and “positive regulation of cell projection organization” (Knn.m2; Figure S20D). Terms related to chromosome organization, DNA/RNA metabolism, and protein modification (PG.m4 and PG.m11; Figure 4H and S20D) were enriched in younger oligodendrocytes (OLI1–3). In older oligodendrocytes (OLI4–6), terms related to neurotransmitter signaling, synaptic structure and plasticity, and higher functions (such as learning, memory, and speech) were enriched (PG.m8 and Knn.m9; Figure 4H and S20D). It has been shown that oligodendrocytes are highly responsive to neuronal activity (Gibson et al., 2014; Hines et al., 2015; Hrvatin et al., 2018), and their rapid production is important for motor skill learning (McKenzie et al., 2014; Xiao et al., 2016). These results suggest neuronal activity-dependent specification of OLI4–6, consistent with findings from studies of adaptive myelination (Baraban et al., 2016; Bechler et al., 2018; McKenzie et al., 2014; Young et al., 2013).

Interestingly, we found that *MUSK* was differentially enriched in OLI4–5 (Figure 4B). *MUSK* is well known to function at the neuromuscular junction and, but its expression in the brain is thought to mediate cholinergic responses, synaptic plasticity, and memory formation (Garcia-Osta et al., 2006). The expression of *MUSK* in oligodendrocytes has not been reported previously. In marmoset, we found it almost exclusively in oligodendrocytes and a group of *PDGFRA*^+^ vascular and leptomeningeal cells (VLMC1) (Figure S20E). Clinically, autoantibodies against *MuSK* are characteristic in many cases of acetylcholine receptor antibody-negative myasthenia gravis, and coincident central nervous system demyelination in these cases may be underreported (Sylvester et al., 2013). MUSK expression in oligodendrocytes seems to be unique to primate, as it is also detected in human oligodendrocytes (Figure S21) but not in mouse.

### Astrocyte subtypes are segregated by compartments of the neural tube

We observed shared transcriptomic features and intermingled distribution of nuclei from astrocyte (AST) and vascular (VAS) classes, so we pooled these two classes for the second round of quality control to facilitate artifact imputation (Figure S24). A total of 74,204 nuclei were studied, which included the major cell types present at CNS barriers (astrocytes, endothelial cells, ependymal cells, and meningeal cells) (Figure S25 – 28). The localization of marker gene expression for these cell types is demonstrated in reference to *in situ* hybridization (ISH) on P0 marmoset brain from the Marmoset Gene Atlas (Shimogori et al., 2018), which was matched to an adult marmoset MRI (Marmoset Brain Mapping, (Liu et al., 2020; 2018)) (Figure S25).

As for microglia and OPC, we observed spatial separation of AST subclusters across the brain after partitioning a total of 61,147 nuclei into 8 subclusters (Figure 5A, S26). GM-enriched astrocytes (AST1) and WM-enriched astrocytes (AST2) were identified (Bayraktar et al., 2018), but subclusters AST4 and AST5 were also enriched in the posterior corpus callosum (pCC) and optic tract (OpT) relative to other WM areas. Interestingly, the AST subcluster composition in these two WM structures was similar to that found in thalamus (including lateral geniculate nucleus), midbrain, pons, and cervical spinal cord. This result leads to the prediction that the brain’s response to injury may not be consistent across WM areas. With respect to gray matter, the astrocyte composition of the caudate and hippocampus resembled that in GM samples, whereas the cerebellum stood out in a category of its own (Figure 5A, S26).

**Figure 5.**
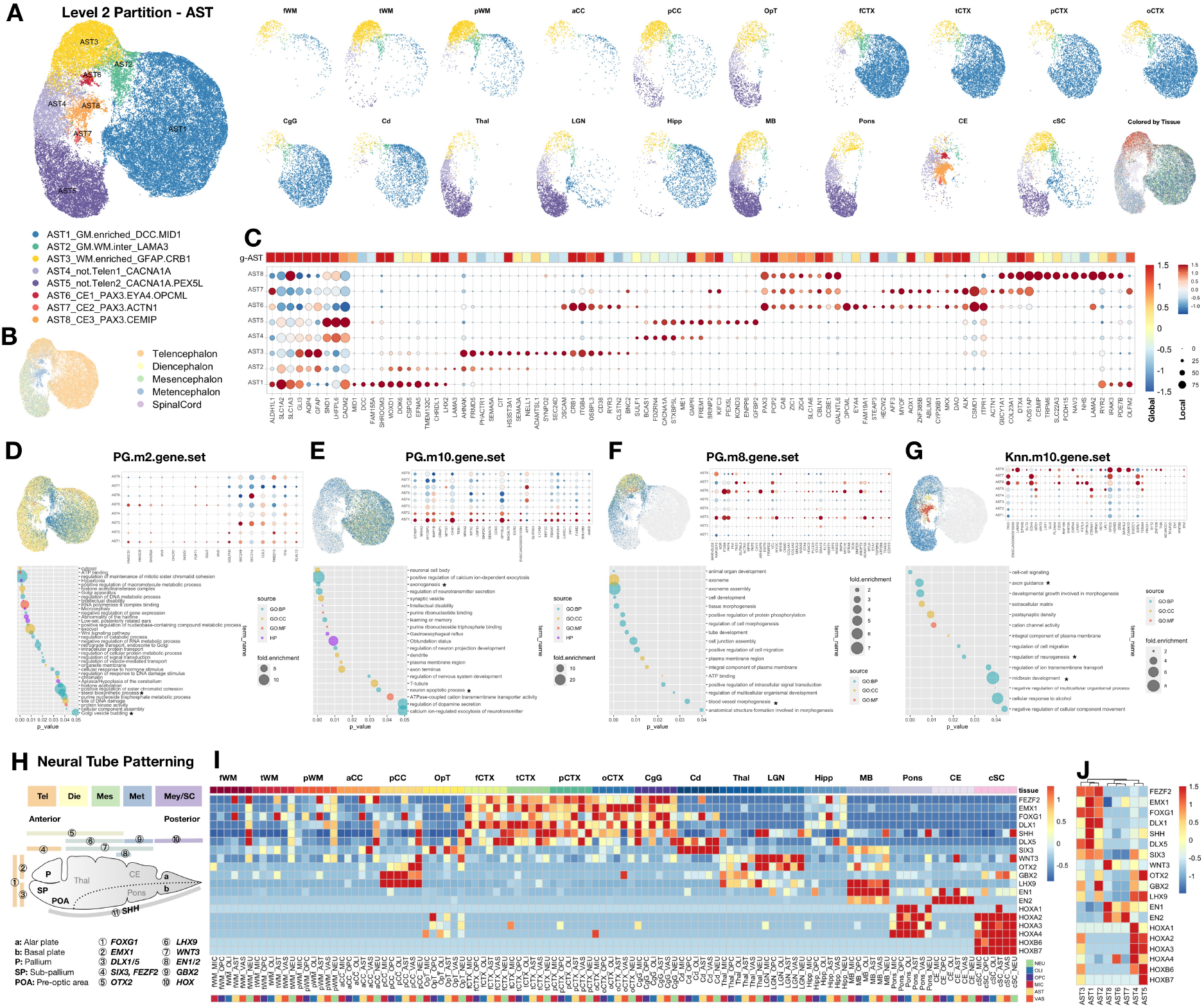
Astrocyte subclusters are spatially segregated and respect the boundaries of secondary vesicles during neural tube development. (A) UMAP scatter-plot visualization of astrocytes (AST) colored by subcluster identity and split by sampling site. (B) UMAP scatter-plot visualization of AST nuclei colored by tissue type in developmental category (See Figure 1D for list). (C) Heatmap and dot plot showing mean-centered and z-score-scaled marker gene expression in global space (heatmap, AST average expression among Level 1 classes, C50) and local space (dot plot, AST subclusters). The portion of the heatmap that represents global AST (g-AST) is reproduced here from the full heatmap shown in Figure S26F. (D – G) Upper left, UMAP scatter-plot visualization of AST nuclei colored by averaged expression of selected modules. Bottom, dot plots showing enriched Gene Ontology (GO) terms from the list of genes in each module; selected terms for the marker expression plot are annotated (*). Upper right, dot plots showing the detected genes from selected terms. Examples include “sterol biosynthetic process” and “Golgi vesicle budding” (D); “axonogenesis” and “neuron apoptotic process” (E); “blood vessel morphogenesis” (F); “axon guidance,” “regulation of neurogenesis,” and “midbrain development” (G). (H) Schematic illustration of morphogen expression along the anterior-posterior axis of the neural tube during development. See Figure 1 for tissue label abbreviation list. (I) Heatmap showing the expression of selected morphogens in Level 1 classes, split by sampling site. The sampling sites are ordered approximately along the anterior-posterior axis of the neural tube during development from left to right (tWM to cSC); the genes that are enriched along the same axis are ordered from top to bottom (*FEZF2* to *HOXB7*). (J) Heatmap showing the expression of selected morphogens across Level 2 subclustering in astrocytes. See also Figure S24-28, Table S2 and S4.

These observations led us to consider ways to group tissue that might better explain this segregation developmentally. Similar to a published mouse brain atlas (Zeisel et al., 2018), we found that grouping tissue by their secondary vesicle origin during neural tube patterning, together with WM-GM disparity, most effectively describes astrocyte segregation, with some exceptions. We defined telencephalon white (fWM, tWM, pWM, aCC, pCC), telencephalon gray (fCTX, tCTX, pCTX, oCTX, CgG, Cd, and Hipp), non-telencephalon (diencephalon: OpT, Thal; mesencephalon: MB; metencephalon: Pons; spinal cord: cSC), and metencephalon (CE) as the 4 categories on which we mapped the landscape of AST populations (Figure 5B). We found common markers, such as *ALDH1L1* and *GLI3*, most effectively label the whole lineage of astrocytes across regions, including Bergmann glia (AST8) in the cerebellum (Figure 5C). *SLC1A2* is enriched in GM astrocytes and *GFAP* and *AQP4* in WM astrocytes.

To further understand the functional implication of this segregation, we performed gene module and GO analysis (Figure S26, Table S2) as described above. As a positive control, we found that astrocytes are generally involved in “sterol biosynthesis process,” as they are the major cholesterol producer in the brain (Figure 5D). As with other cell types, terms related to neuron transmitter secretion and nervous system development were enriched in GM-enriched astrocytes (AST1) (Figure 5E). WM-enriched astrocytes (AST3) were enriched with terms related to cell migration, intracellular signaling transduction, and blood vessel morphogenesis (Figure 5F). Cerebellar astrocytes (AST6–8) were enriched in genes involved in axon guidance and neurogenesis (Figure 5G).

### The transcriptome of white matter cells deviates from the profile of forebrain patterning

To further investigate the effect of local patterning signals in defining AST subclusters and to assess whether these signals also affect other cell types in the same region, we examined the expression of morphogens along the anterior-posterior (AP) axis across cell and tissue types (Figure 5H). Indeed, morphogen patterning across brain regions was grossly preserved across cell types, with some interesting exceptions. In telencephalon, all cell types in cortical gray matter expressed high levels of morphogens expressed most prominently in the forebrain (*FEZF2*, *EMX1*, *FOXG1*, *DLX1*, *SHH*). Whereas caudate was patterned by *DLX5* and *SIX3*, hippocampus, though belonging to telencephalon gray, had an expression pattern that more resembled that of WM. Most WM cells appeared to have lost this specification, except for some *FEZF2* and *FOXG1* expression in astrocytes and neurons. Cells in posterior corpus callosum, however, had a morphogen pattern similar to what was observed in thalamus, LGN, and midbrain (*WNT3*, *OTX2*, *GBX2*, *LHX9*, *EN1*). *EN2* was enriched in cerebellum, but hindbrain morphogens (*HOX* genes) were high in pons and cervical spinal cord and sporadically spotted in optic tract (Figure 5I). This forebrain, midbrain, and hindbrain specification was preserved more prominently in AST than any other cell class, and indeed determined their identity as distinct AST subclusters (Figure 5J).

### Gray matter glia share regulatory pathways

The GM-WM segregation of glia and our observation of transcriptional similarity across glia in each tissue type led us to hypothesize that there might be pathways that are shared across cell classes within the same microenvironment. To explore this possibility, we extracted and compared differentially expressed transcription factors between analogous GM- and WM-specific glia pairs (MIC1 vs. MIC3, OPC1 vs. OPC3, AST1 vs. AST3) (Figure 6A). We found greater overlap in differentially enriched transcription factors in GM-glia (15 overlapping transcription factors) compared to WM-glia (3 overlapping transcription factors). Interestingly, 6 transcription factors (*EGR1*, *HLF*, *PEG3*, *MYT1L*, *HIVEP2*, *BHLHE40*) were shared across MIC1, OPC1, and AST1, whereas no transcription factor was shared across the WM-glia classes MIC3, OPC3, and AST3 (Figure 6A–B). These GM-glia transcription factors are known to restrict RNA biosynthesis (Figure 6C), consistent with the observation that GM-glia are low in RNA complexity compared to their WM counterparts (Figure 6D). We further quantified the similarity of GO terms found in each gene module by calculating the Jaccard index between module pairs across cell classes and visualizing their similarity as networks (Figure 6E). Again, GO terms were more similar among gene modules enriched in GM-glia than other gene modules, and regulatory programs in GM-microglia showed highest similarity with those in GM-OPC (Figure 6F).

**Figure 6.**
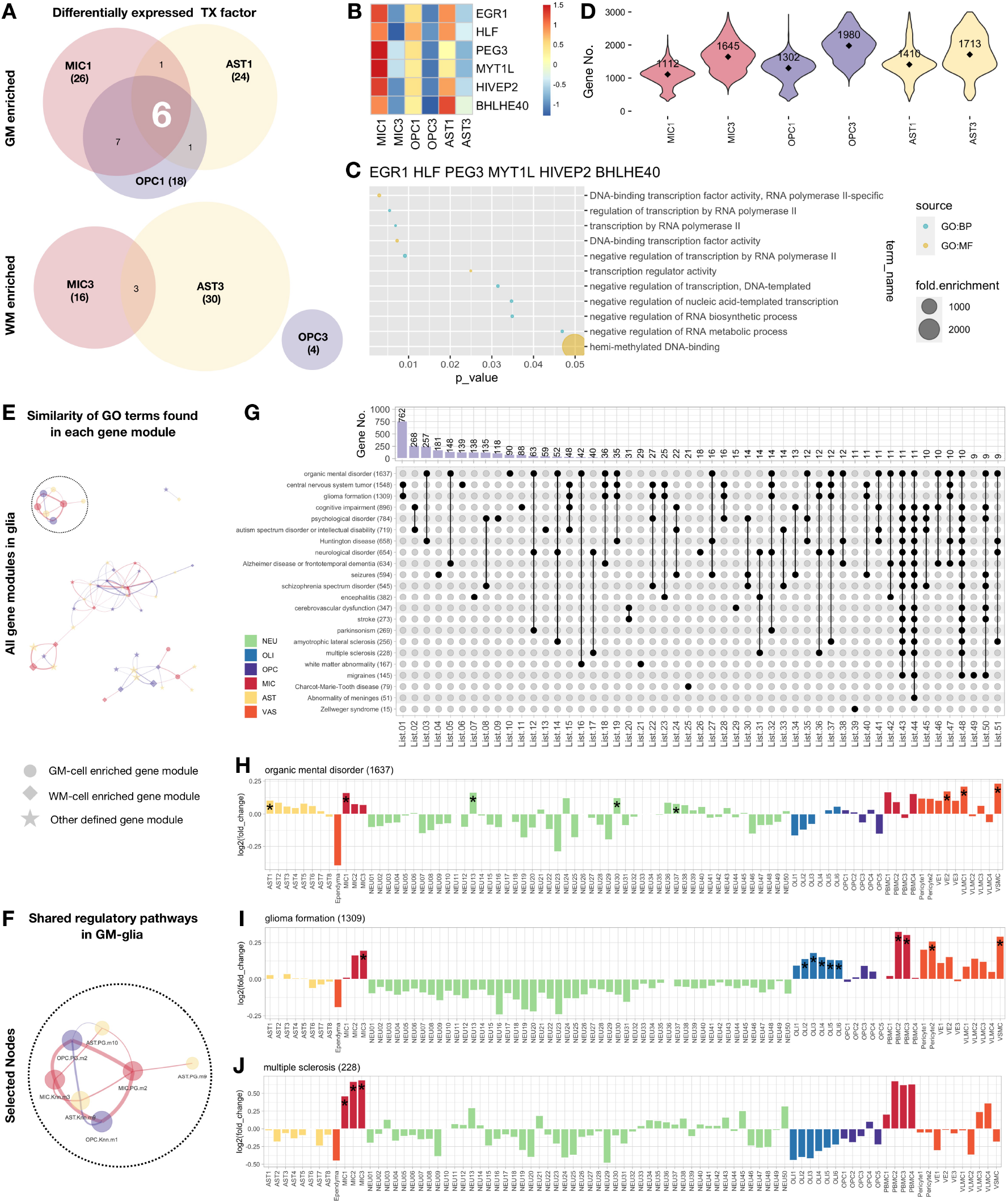
Gray matter and white matter glia are distinguished by their shared regulatory pathways and contribution to neurological disorders. (A) Venn diagram showing the shared transcription factors across gray matter glia (top) and white matter glia (bottom). Differentially enriched transcription factors between gray and white matter subclusters are tabulated (MIC1 against MIC3, OPC1 against OPC3, AST1 against AST3). (B) Heatmap showing the expression of the 6 shared transcription factors enriched in GM glia. (C) Dot plot showing the enriched GO terms from the 6 shared transcription factors that are enriched in GM glia. GM glia are enriched with regulatory pathways that negatively impact RNA biosynthesis, consistent with the lower number of genes detected in GM glia compared to their WM counterpart (See Figure S13B, S15B, S26B). (D) Violin plot showing the number of genes detected in each cluster. Median is annotated (◆). (E) Network plot showing the similarity of GO terms identified in each gene module. The size of nodes in the network represents the number of significant GO terms found in each gene module, and nodes are annotated by shape to reflect differential loading of each module (dot, gene module enriched primarily in GM-glia; diamond, gene module enriched primarily in WM-glia; star, gene module enriched in multiple and/or “other” tissues). Edges between nodes are scaled to reflect similarity between two list of GO terms: thicker edges indicate greater similarity between the gene modules. Only edges with Jaccard index >0.25 are shown to avoid crowding, which results in the separation into 3 distinct networks. (F) Enlargement of the network plot indicated in the dashed circle showed in Figure 6E. (G) Bar and UpSet plots showing the overlap of genes associated with a variety of neurological disorders as defined in the IPA database. The number of genes is listed next to the name of the disorder (bottom panel, y axis label). The number of intersecting genes between indicated disorders (solid black dot and line), but not shared by any other disorders (empty gray dot), is labeled and shown on the bar graph (top panel). (H) Bar graph showing strong enrichment of genes associated with organic mental disorder in GM microglia (MIC1) and GM astrocytes (AST1). (I) Bar graph showing strong enrichment of genes associated with glioma formation in WM microglia (MIC3) as well as oligodendrocytes, but not astrocytes. (J) Bar graph showing strong enrichment of genes associated with multiple sclerosis in all microglia subclusters. See also Figure S29, Table S2 and S4.

### White and gray matter glia differentially contribute to neurological disorders

We explored bioinformatically whether regionally diverse glia are predicted to function differently in pathological conditions. We reasoned that by examining the expression of disease-associated genes in our healthy marmoset transcriptomic atlas, we might identify previously overlooked cellular contributors to human neurological disease. We therefore examined the cellular enrichment of genes associated with a spectrum of disorders using Expression-Weighted Cell-type Enrichment (EWCE) analysis (Skene and Grant, 2016) (Figure S29). Based on manually curated information in the Ingenuity Pathway Analysis (IPA) database (Table S4), we sorted genes into lists, ordered them based on phenotypical similarity between disorders, and displayed the number of candidate genes in each list that were unique or shared across disorders (Figure 6G); lists with <9 candidate genes were dropped for simplicity. As expected, genes annotated as being associated with CNS tumors were shared with those associated with glioma formation (762 genes), cognitive impairment genes were shared with autism spectrum disorder or intellectual disability genes (268), and organic mental disorder genes were shared with Alzheimer disease or frontotemporal dementia genes (148 genes). Consistent with the microenvironment specialization of glia reported here, we found examples in which genes associated with particular disorders were differentially expressed in GM-microglia only (e.g., organic mental disorder) and WM-microglia only (e.g, glioma formation) (Figure 6H, I). By contrast, all microglia subtypes, but not other cell types, appear to contribute to multiple sclerosis pathogenesis (Figure 6J) (International Multiple Sclerosis Genetics Consortium, 2019).

### White matter glia interact with other resident cells more than gray matter glia

As described above, we observed the greatest GM-WM spatial segregation in subclusters of microglia, OPC, and astrocytes. GM cells were generally naïve, protoplasmic, and enriched in GO terms related to neuronal functioning, whereas WM cells were more active, fibrous, and enriched in GO terms related to morphogenesis and signaling dynamics. We therefore reasoned that intercellular communication within the different microenvironments may contribute to subcluster specification. To test this hypothesis, we modeled ligand-receptor interactions between cells found within either GM or WM using NicheNet (Browaeys et al., 2020), which curates known ligand-receptor and receptor-target relationships and ranks them based on level of support in published literature. We performed this analysis taking the major GM and WM subclusters of microglia, OPC, and astrocytes as “receivers” and other cells in the same tissue type as “senders” (Figure 7A – C).

**Figure 7.**
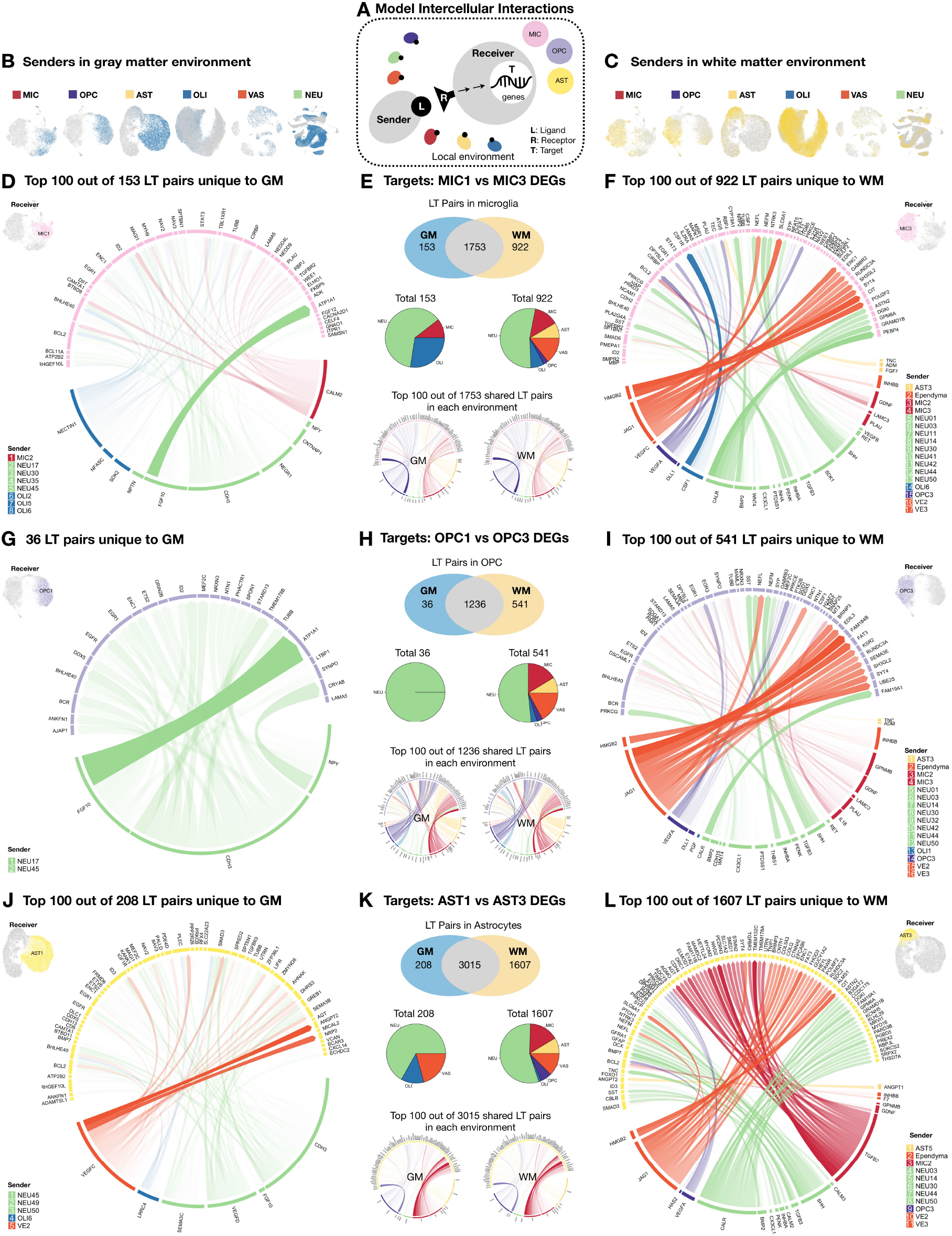
Interactions between resident cells are more extensive in white matter than in gray matter. (A) We modeled intercellular communication in cerebral white matter (WM) and cortical gray matter (GM) microenvironments (see Figure 1D for list) using a pre-established database in NicheNet (Browaeys et al., 2020), which contains information about ligand-receptor and receptor-target pairs. We aimed to identify cell types (“senders”) that express ligands with matching receptors in target cells (“receivers”). To limit false positivity, we kept linked pairs only if the expression of downstream target genes also matched the receptor-target pairing information in the database. All cell classes defined in Level 1 analysis were included as senders in each microenvironment, and their influence on microglia, OPC, and astrocytes were modeled. (B) Senders present in GM are colored blue for each cell class. (C) Senders present in WM are colored yellow for each cell class. (D – F) Genes differentially expressed between MIC1 in GM and MIC3 in WM were set as target genes in the NicheNet analysis. Ligand genes expressed by senders in GM or WM (coarse category, see Figure 1D for list) were extracted and filtered separately, and links to receptors expressed by MIC1 or MIC3 were matched and scored by NicheNet independently. The number of final established ligand-target (LT) pairs in GM or WM tissue are compared in Venn diagrams (E, top). The number of shared ligand-target pairs (1753) and the top 100 features in each environment are plotted in the center (E, bottom; see Figure S30 for full panel). Sender cell types (in Level 1 classes) for ligand-target pairs that are unique to each environment are further depicted in pie charts (E, middle). In the 153 ligand-target pairs that were unique to GM, ligands were found from neurons, oligodendrocytes, and microglia (D). In the 922 ligand-target pairs that were unique to WM tissue, ligands were found from additional cell types (astrocytes, vascular cells, and OPC, F). (G – I) Genes differentially expressed between OPC1 in GM and OPC3 in cWM were set as target genes in the NicheNet analysis. Ligand genes expressed by senders in GM or WM (coarse category, see Figure 1D for list) were extracted and filtered separately, and links to receptors expressed by OPC1 or OPC3 were matched and scored by NicheNet independently. The number of final established ligand-target (LT) pairs in GM or WM tissue are compared in Venn diagrams (H, top). The number of shared ligand-target pairs (1236) and the top 100 features in each environment are plotted in the center (H, bottom; see Figure S30 for full panel). Sender cell types (in Level 1 classes) for ligand-target pairs that are unique to each environment are further depicted in pie charts (H, middle). In the 36 ligand-target pairs that were unique to GM, ligands were found from neurons, oligodendrocytes, and microglia (G). In the 541 ligand-target pairs that were unique to WM tissue, ligands were found from additional cell types (microglia, astrocytes, vascular cells, oligodendrocytes, and OPC, I). (J – L) Genes differentially expressed between AST1 in GM and AST3 in WM were set as target genes in the NicheNet analysis. Ligand genes expressed by senders in GM or WM (coarse category, see Figure 1D for list) were extracted and filtered separately, and links to receptors expressed by AST1 or AST3 were matched and scored by NicheNet independently. The number of final established ligand-target (LT) pairs in GM or WM tissue are compared in Venn diagrams (K, top). The number of shared ligand-target pairs (3015) and the top 100 features in each environment are plotted in the center (K, bottom; see also Figure S30 for full panel). Sender cell types (in Level 1 classes) for ligand-target pairs that are unique to each environment are further depicted in pie charts (K, middle). In the 208 ligand-target pairs that were unique to GM, ligands were found from neurons, oligodendrocytes, and microglia (J). In the 1607 ligand-target pairs that were unique to WM tissue, ligands were found from additional cell types (astrocytes, vascular cells, and OPC, L). See also Figure S30 and Table S3.

We consistently found more, and more unique, ligand-target pairs in WM than in GM, and these were generated by a wider variety of sender types (Table S3 and Figure 7E, H, K). In WM, endothelial cells were the most frequently observed additional sender types (Figure 7F, I, L). Astrocytes (4,830 ligand-target pairs) formed more ligand-target pairs than microglia (2,828 pairs) or OPC (1,813 pairs). Among ligand-target pairs found in both GM and WM (1,753 for microglia, 1,236 for OPC, 3,015 for astrocytes), certain senders (Figure S21, pie charts) were represented disproportionately to their relative abundance (Figure 1E, S3E), with microglia and OPC constituting the most common top-ranked senders (Figure S30, Circos plots). This is consistent with prior reports that microglia and OPC can survey their microenvironments in both physiological and pathological conditions (Kirby et al., 2019; Nimmerjahn et al., 2005).

## Discussion

We characterized >500,000 nuclei across adult marmoset brain and spinal cord and found that the gray and white matter microenvironments dramatically impact the molecular profile of resident cells, suggesting both developmental and functional differences. We observed that the molecular complexity of glia is highest in white matter (Figure 6D, S13B, S15B, S26B), where transcriptomic profiles deviate most from those observed in the developing neural tube (Figure 5I), and where there is more communication between nearby cells (Figure 7E, H, K). We found shared master regulators that negatively impact transcription (Figure 6C), and shared pathways that are involved in neuronal support (Figure 6E, F), in all gray matter glia. Whether these transcriptomically heterogeneous subclusters defined in this study respond differently to stimuli in pathological conditions remain to be addressed; however, we found initial evidence that gray and white matter glia differentially contribute to neurological disorders (Figure 6G).

The GM-WM segregation of the microglial transcriptome was observed as early as P7 (during myelinogensis) in mouse (Staszewski and Hagemeyer, 2019) and persisted along the trajectory of normal aging in both human and mouse (Gefen et al., 2019; Hart et al., 2012; Raj et al., 2017). Significantly more microglia are found in WM than in GM of normal human brain (Mittelbronn et al., 2001), and WM-microglia are primed to be more active and respond to injury faster than their GM counterparts (Bachstetter et al., 2013; Cătălin et al., 2013; Hart et al., 2012; Staszewski and Hagemeyer, 2019; van der Poel et al., 2019). Consistent with previous reports, we found 3 times more microglia in WM than in GM tissue (Figure S3E), and the expression of genes related to stimulus-response (such as CD11b, CD11c, and CD64) were elevated in WM-microglia. It is still not clear if these two populations diverge autonomously (such as from different progenitors) or are shaped environmentally; however, we observed more intercellular activity for WM-microglia than GM-microglia (Figure 7E). Additional transcriptomic comparison at a time point before the onset of myelination will be necessary to address whether the relative absence of neuron cell bodies or the presence of excess myelin explains microglial priming in WM.

The effect of the WM environment on priming glia to be more advanced in response to pathological challenges was also observed for astrocytes (Bachstetter et al., 2013; Shannon et al., 2007). WM-astrocytes have a higher capacity for glutamate clearance to handle excitotoxic insults, and disproportionally higher senescence-induced expression of GFAP, a reactive gliosis indicator, than GM-astrocytes (Lundgaard et al., 2014). Similarly, migratory pathways appear to be active in marmoset WM-astrocytes (Figure 5F), and the differential interactivity of cells in each milieu (Figure 7K) we observed in this study might explain the GM-WM disparity of astrocytes. However, multiple lineage-tracing experiments support that there is clonal exclusion to generate either protoplasmic (GM enriched) or fibrous (WM enriched) astrocytes, which are likely to be produced by independent progenitors (Tabata, 2015). In line with this view, we found that even anterior-posterior patterning genes were grossly preserved across all types of glia (Figure 5I), whereas regional determination of subset identities was only present in astrocytes (Figure 5J). In summary, although environmental cues contribute to diversifying astrocytes, our data suggest that the heterogeneity in their developmental origin plays a bigger role in subtype specialization.

Similar with our finding that marmoset gray matter microglia appear younger than their white matter counterparts (Figure 2), it has been reported that rat white matter OPC are more mature than gray matter OPC (Lentferink et al., 2018) and differentiate into mature oligodendrocytes more efficiently (Viganò et al., 2013). Electrophysiological properties of OPC vary between white and gray matter and with age, and they correlate with differentiation potentiality (Chittajallu et al., 2004; Spitzer et al., 2019). Therefore, we examined the OPC expression of ion channel genes as a surrogate of electrophysiological function, and examined the tissue origin of differentiating OPC (OPC5). We found differential expression of ion channels between gray and white matter OPC (Figure S16) but a similar abundance (<1.5%) of OPC5 across brain regions (Figure 3C). Further study will be necessary to address whether tissue-of-origin influences the relative abundance of OPC5 under conditions, such as development or repair, where OPC differentiation is stimulated.

We also interrogated the similarities and differences of oligodendrocyte lineage cells across species and developmental trajectories in depth by re-analyzing data from prior studies in zebrafish (Marisca et al., 2020), mouse (Marques et al., 2016; Zeisel et al., 2018; Zhang et al., 2014), and human (Habib et al., 2017; Lake et al., 2018; Polioudakis et al., 2019; Zhang et al., 2016). Similar to OPC derived from adult human brain, we did not observe a separate cycling OPC cluster (*TOP2A*^+^) in adult marmoset brain, as has been reported for OPC in derived from zebrafish, adult mouse, and developing human cortex (Figure S17–18). Instead, cells expressing S and G2/M phase genes were dispersed within OPC1–3 subclusters. OPC4, however, was enriched with genes that defined as G0G1 (Figure 3B), i.e., quiescent cells (Feldman et al., 2019). Marmoset white matter OPC (OPC3) had lower transcriptomic agreement with OPC from other datasets than marmoset gray matter OPC (Figure 3I), supporting our speculation that white matter glia might be “older,” i.e., further deviated from their developmental status. Perhaps surprisingly, differentiating OPC (*MAG*^+^ or *ENPP6*^+^) in zebrafish and mouse (dr, mm1, mm2; Figure 3I) were more similar to marmoset oligodendrocytes (OLI1–6) than to marmoset differentiating OPC (OPC5). In contrast, human (hs1, hs3) differentiating OPC had high agreement with marmoset OPC5, and similarity with marmoset oligodendrocytes gradually decreased from OLI6 to OLI1. These observations suggest either that OPC5 is a transitional state unique to primates, or that mouse OPC have a much quicker transcriptomic switch from OPC to oligodendrocytes that is hard to capture in single cell analysis.

The finding that OLI6 are most similar to OPC in other species runs counter to our initial definition of *ENPP6*^high^ cells, most common in OLI1, as the “youngest” oligodendrocyte subcluster (Xiao et al., 2016). To resolve this paradox, we note that across both species (rodents and primates) and techniques (scRNA-seq and snRNA-seq), oligodendrocyte-lineage cells are arranged into a continuous path in dimension-reduced space (tSNE or UMAP); the most parsimonious explanation of this observation is the trajectory of lineage development. There is experimental data in support of this idea from mouse studies, in which clusters of oligodendrocyte lineage have been defined that explain graded transcriptomic changes toward terminal differentiation. Despite some discrepancy in the assignment of subcluster identity, 5 major stages of oligodendrocyte lineage cells are widely accepted: OPC, differentiation-committed oligodendrocyte precursors (COP), newly formed oligodendrocytes (NFOL), myelin-forming oligodendrocytes (MFOL), and mature oligodendrocytes (MOL). Unfortunately, there is no single marker or small group of markers that can cleanly separate these groups. To gain more insight, we compared the transcriptome of a bulk-RNA sequencing data from immunopanned neural cells (Zhang et al., 2014) and two single-cell sequencing datasets in mouse (Marques et al., 2016; Zeisel et al., 2018). We found that the transcriptome of NFOL, as defined by a cell surface marker (GalC^+^), was most closely correlated with that of mature oligodendrocytes, as defined by single cell analysis in both datasets, and the pattern of correlation did not track with the ordering of those oligodendrocyte subclusters in UMAP space (Figure S23). Interestingly, we found the same correlational pattern for immunopanning-defined MO. Together, these observations raise the possibility that the single-path model of the differentiation trajectory of oligodendrocytes requires modification.

Although our study was carefully designed and executed, and rigorous quality control steps were implemented at every stage of the experimental and analysis pipeline, technical variation and artifacts remain intermingled with biological effects. For example, spinal cord samples were outside the region covered by the MRI atlas, and results were derived from 2 libraries prepared with 10x v2 and 1 library with 10x v3 chemistry (fewer genes were recovered using v2 than from v3 on average). Some spinal cord neurons and cerebellar Purkinje cells were likely under-sampled, as they are larger and might not be captured in droplets or pass through microfluidic chambers similarly to smaller cells. Additionally, while snRNA-seq has several advantages, including higher tolerance for tissue processing, it captures only nascent RNAs and cannot interrogate locally enriched species along neural cell processes. Another limitation is that annotations for genes and transcripts, and associated functional terms, are less complete for marmoset than for mouse or human, and many annotations are inferred and might change in the future. We were therefore relatively conservative in clustering and defining cell types, and it is likely that further subclustering would yield more distinct cell types. In our work, we defined cell types and linked their molecular properties to functions by pathway analysis and disease association mapping, but morphological and electrophysiological features remain unlinked. With respect to sampling, although we profiled as many nuclei as possible in each round, often <1% of nuclei were profiled from each region (Figure S1E), meaning that rare subclusters were probably missed. Finally, given finite resources we elected to sample the brain richly rather than to include samples from additional marmosets, limiting, for example, our ability answer sex-specific or left-right asymmetry-related questions. These limitations aside, our analysis provides a framework for understanding the diversity of cell types in the marmoset brain and lays the groundwork for future work.

In summary, we developed a protocol that is easily adapted to other settings and allows nuclei to be mapped onto relatively small regions to increase reproducibility and aid future validation studies. We focused less on using one or a few genes to define cellular subtype and instead considered the whole transcriptome to highlight relative differences in function among similar cells of the same class. We demonstrated ways of using our data to classify unknown cell types, query intercellular communication, and discover associations with disease. We hope that the carefully annotated marmoset brain cell atlas we have built, CjPCA, will benefit and inform future studies in evolutional, developmental, and pathological neurobiology.

## Data availability

Raw and processed datasets will be submitted to Gene Expression Omnibus (GEO). Data can also be visualized at https://cjpca.ninds.nih.gov.

## Acknowledgments

We thank Dr. Pascal Sati, Dr. Maxime Donadieu, and Mr. Roger Depaz (TNS) for help in collecting in vivo MRI data and monitoring animals. We thank Dr. Ariel Levine (NINDS), Ms. Kaya J.E. Matson (NINDS), Dr. Yuesheng Li (NHLBI), Dr. Poching Liu (NHLBI), Dr. Yan Luo (NHLBI), Dr. Qing Wang (UCLA), and the Adelson Medical Research Foundation (AMRF) Functional Genomics Resource (UCLA) for expertise and assistance with sequencing. We thank Dr. Chang-Ting Lin for insights in quantitative data analysis. We thank the AMRF Program in Neurodegenerative Diseases–Multiple Sclerosis (APND-MD) and the Intramural Research Program of NINDS for funding.

## Author contributions

J-PL and DSR designed the study and interpret the results. J-PL, DSR, and SJ developed protocols. J-PL and DSR wrote and prepared manuscript. J-PL acquired, processed, and analyzed snRNAseq data. J-PL and HMK acquired, processed, and analyzed histology data. J-PL and YS cleaned and processed published datasets. DHG and RK analyzed data. DSR supervised the study.

## References

Bachstetter, A.D., Webster, S.J., Van Eldik, L.J., and Cambi, F. (2013). Clinically relevant intronic splicing enhancer mutation in myelin proteolipid protein leads to progressive microglia and astrocyte activation in white and gray matter regions of the brain. J Neuroinflammation 10, 1–14.

Bakken, T.E., Jorstad, N.L., Hu, Q., Lake, B.B., Tian, W., Kalmbach, B.E., Crow, M., Hodge, R.D., Krienen, F.M., Sorensen, S.A., et al. (2020). Evolution of cellular diversity in primary motor cortex of human, marmoset monkey, and mouse. bioRxiv 573, 2020.03.31.016972.

Baraban, M., Mensch, S., and Lyons, D.A. (2016). Adaptive myelination from fish to man. Brain Res 1641, 149–161.

Bartheld, von, C.S., Bahney, J., and Houzel, S.H. (2016). The search for true numbers of neurons and glial cells in the human brain: A review of 150 years of cell counting. J Comp Neurol 524, 3865–3895.

Bayraktar, O.A., Bartels, T., Polioudakis, D., Holmqvist, S., Ben Haim, L., Young, A.M.H., Prakash, K., Brown, A., Paredes, M.F., Kawaguchi, R., et al. (2018). Single-cell in situ transcriptomic map of astrocyte cortical layer diversity. bioRxiv 432104.

Bechler, M.E., Swire, M., and ffrench-Constant, C. (2018). Intrinsic and adaptive myelination-A sequential mechanism for smart wiring in the brain. Dev Neurobiol 78, 68–79.

Browaeys, R., Saelens, W., and Saeys, Y. (2020). NicheNet: modeling intercellular communication by linking ligands to target genes. Nature Methods 2017 14:3 17, 159–162.

Cao, J., Spielmann, M., Qiu, X., Huang, X., Ibrahim, D.M., Hill, A.J., Zhang, F., Mundlos, S., Christiansen, L., Steemers, F.J., et al. (2019). The single-cell transcriptional landscape of mammalian organogenesis. Nature 566, 496–502.

Cătălin, B., Mitran, S., Albu, C., and Iancău, M. (2013). Comparative aspects of microglia reaction in white and gray matter. Current Health Sciences Journal 39, 151–154.

Chittajallu, R., Aguirre, A., and Gallo, V. (2004). NG2-positive cells in the mouse white and grey matter display distinct physiological properties. J. Physiol. (Lond.) 561, 109–122.

Feldman, H.M., Toledo, C.M., Arora, S., Hoellerbauer, P., Corrin, P., Carter, L., Kufeld, M., Bolouri, H., Basom, R., Delrow, J., et al. (2019). Neural G0: a quiescent-like state found in neuroepithelial-derived cells and glioma. bioRxiv 8, 446344.

Garcia-Osta, A., Tsokas, P., Pollonini, G., Landau, E.M., Blitzer, R., and Alberini, C.M. (2006). MuSK Expressed in the Brain Mediates Cholinergic Responses, Synaptic Plasticity, and Memory Formation. Journal of Neuroscience 26, 7919–7932.

Gefen, T., Kim, G., Bolbolan, K., Geoly, A., Ohm, D., Oboudiyat, C., Shahidehpour, R., Rademaker, A., Weintraub, S., Bigio, E.H., et al. (2019). Activated Microglia in Cortical White Matter Across Cognitive Aging Trajectories. Front Aging Neurosci 11, 1090.

Gibson, E.M., Purger, D., Mount, C.W., Goldstein, A.K., Lin, G.L., Wood, L.S., Inema, I., Miller, S.E., Bieri, G., Zuchero, J.B., et al. (2014). Neuronal activity promotes oligodendrogenesis and adaptive myelination in the mammalian brain. Science 344, 1252304–1252304.

Habib, N., Avraham-Davidi, I., Basu, A., Burks, T., Shekhar, K., Hofree, M., Choudhury, S.R., Aguet, F., Gelfand, E., Ardlie, K., et al. (2017). Massively parallel single-nucleus RNA-seq with DroNc-seq. Nat. Methods 14, 955–958.

Hammond, T.R., Dufort, C., Dissing-Olesen, L., Giera, S., Young, A., Wysoker, A., Walker, A.J., Gergits, F., Segel, M., Nemesh, J., et al. (2019). Single-Cell RNA Sequencing of Microglia throughout the Mouse Lifespan and in the Injured Brain Reveals Complex Cell-State Changes. Immunity 50, 253–271.e256.

Hart, A.D., Wyttenbach, A., Hugh Perry, V., and Teeling, J.L. (2012). Age related changes in microglial phenotype vary between CNS regions: Grey versus white matter differences. Brain Behav Immun 26, 754–765.

Hines, J.H., Ravanelli, A.M., Schwindt, R., Scott, E.K., and Appel, B. (2015). Neuronal activity biases axon selection for myelination in vivo. Nat. Neurosci. 18, 683–689.

Hodge, R.D., Miller, J.A., Novotny, M., Kalmbach, B.E., Ting, J.T., Bakken, T.E., Aevermann, B.D., Barkan, E.R., Berkowitz-Cerasano, M.L., Cobbs, C., et al. (2020). Transcriptomic evidence that von Economo neurons are regionally specialized extratelencephalic-projecting excitatory neurons. Nature Communications 11, 1172–14.

Hrvatin, S., Hochbaum, D.R., Nagy, M.A., Cicconet, M., Robertson, K., Cheadle, L., Zilionis, R., Ratner, A., Borges-Monroy, R., Klein, A.M., et al. (2018). Single-cell analysis of experience-dependent transcriptomic states in the mouse visual cortex. Nat. Neurosci. 21, 120–129.

International Multiple Sclerosis Genetics Consortium (2019). Multiple sclerosis genomic map implicates peripheral immune cells and microglia in susceptibility. Science 365, eaav7188.

Jäkel, S., Agirre, E., Mendanha Falcão, A., van Bruggen, D., Lee, K.W., Knuesel, I., Malhotra, D., ffrench-Constant, C., Williams, A., and Castelo-Branco, G. (2019). Altered human oligodendrocyte heterogeneity in multiple sclerosis. Nature 566, 543–547.

Kirby, L., Jin, J., Cardona, J.G., Smith, M.D., Martin, K.A., Wang, J., Strasburger, H., Herbst, L., Alexis, M., Karnell, J., et al. (2019). Oligodendrocyte precursor cells present antigen and are cytotoxic targets in inflammatory demyelination. Nature Communications 10, 1–20.

Krienen, F.M., Goldman, M., Zhang, Q., C H Del Rosario, R., Florio, M., Machold, R., Saunders, A., Levandowski, K., Zaniewski, H., Schuman, B., et al. (2020). Innovations present in the primate inter-neuron repertoire. Nature 586, 262–269.

Lake, B.B., Chen, S., Sos, B.C., Fan, J., Kaeser, G.E., Yung, Y.C., Duong, T.E., Gao, D., Chun, J., Kharchenko, P.V., et al. (2018). Integrative single-cell analysis of transcriptional and epigenetic states in the human adult brain. Nat. Biotechnol. 36, 70–80.

Lentferink, D.H., Jongsma, J.M., Werkman, I., and Baron, W. (2018). Grey matter OPCs are less mature and less sensitive to IFNγ than white matter OPCs: consequences for remyelination. Sci Rep 8, 1–15.

Liu, C., Ye, F.Q., Newman, J.D., Szczupak, D., Tian, X., Yen, C.C.-C., Majka, P., Glen, D., Rosa, M.G.P., Leopold, D.A., et al. (2020). A resource for the detailed 3D mapping of white matter pathways in the marmoset brain. Nat. Neurosci. 23, 271–280.

Liu, C., Ye, F.Q., Yen, C.C.-C., Newman, J.D., Glen, D., Leopold, D.A., and Silva, A.C. (2018). A digital 3D atlas of the marmoset brain based on multi-modal MRI. Neuroimage 169, 106–116.

Liu, X., Lu, Y., Zhang, Y., Li, Y., Zhou, J., Yuan, Y., Gao, X., Su, Z., and He, C. (2012). Slit2 regulates the dispersal of oligodendrocyte precursor cells via Fyn/RhoA signaling. Journal of Biological Chemistry 287, 17503–17516.

Lundgaard, I., Osório, M.J., Kress, B.T., Sanggaard, S., and Nedergaard, M. (2014). White matter astrocytes in health and disease. Neuroscience 276, 161–173.

Marisca, R., Hoche, T., Agirre, E., Hoodless, L.J., Barkey, W., Auer, F., Castelo-Branco, G., and Czopka, T. (2020). Functionally distinct subgroups of oligodendrocyte precursor cells integrate neural activity and execute myelin formation. Nat. Neurosci. 23, 363–374.

Marques, S., Zeisel, A., Codeluppi, S., van Bruggen, D., Mendanha Falcão, A., Xiao, L., Li, H., Haring, M., Hochgerner, H., Romanov, R.A., et al. (2016). Oligodendrocyte heterogeneity in the mouse juvenile and adult central nervous system. Science 352, 1326–1329.

McKenzie, I.A., Ohayon, D., Li, H., de Faria, J.P., Emery, B., Tohyama, K., and Richardson, W.D. (2014). Motor skill learning requires active central myelination. Science 346, 318–322.

Mecollari, V., Nieuwenhuis, B., and Verhaagen, J. (2014). A perspective on the role of class III semaphorin signaling in central nervous system trauma. Front. Cell. Neurosci. 8, 2674.

Mittelbronn, M., Dietz, K., Schluesener, H.J., and Meyermann, R. (2001). Local distribution of microglia in the normal adult human central nervous system differs by up to one order of magnitude. Acta Neuropathol 101, 249–255.

Nimmerjahn, A., Kirchhoff, F., and Helmchen, F. (2005). Resting microglial cells are highly dynamic surveillants of brain parenchyma in vivo. Science 308, 1314–1318.

Pelvig, D.P., Pakkenberg, H., Stark, A.K., and Pakkenberg, B. (2008). Neocortical glial cell numbers in human brains. Neurobiol. Aging 29, 1754–1762.

Polioudakis, D., la Torre-Ubieta, de, L., Langerman, J., Elkins, A.G., Shi, X., Stein, J.L., Vuong, C.K., Nichterwitz, S., Gevorgian, M., Opland, C.K., et al. (2019). A Single-Cell Transcriptomic Atlas of Human Neocortical Development during Mid-gestation. Neuron 103, 785–801.e788.

Raj, D., Yin, Z., Breur, M., Doorduin, J., Holtman, I.R., Olah, M., Mantingh-Otter, I.J., Van Dam, D., De Deyn, P.P., Dunnen, den, W., et al. (2017). Increased White Matter Inflammation in Aging- and Alzheimer’s Disease Brain. Front Mol Neurosci 10, 853.

Shannon, C., Salter, M., and Fern, R. (2007). GFP imaging of live astrocytes: regional differences in the effects of ischaemia upon astrocytes. J. Anat. 210, 684–692.

Shimogori, T., Abe, A., Go, Y., Hashikawa, T., Kishi, N., Kikuchi, S.S., Kita, Y., Niimi, K., Nishibe, H., Okuno, M., et al. (2018). Digital gene atlas of neonate common marmoset brain. Neurosci. Res. 128, 1–13.

Skene, N.G., and Grant, S.G.N. (2016). Identification of Vulnerable Cell Types in Major Brain Disorders Using Single Cell Transcriptomes and Expression Weighted Cell Type Enrichment. Front Neurosci 10, 16.

Spitzer, S.O., Sitnikov, S., Kamen, Y., Evans, K.A., Kronenberg-Versteeg, D., Dietmann, S., de Faria, O., Jr., Agathou, S., and Káradóttir, R.T. (2019). Oligodendrocyte Progenitor Cells Become Regionally Diverse and Heterogeneous with Age. Neuron 101, 459–471.e5.

Staszewski, O., and Hagemeyer, N. (2019). Unique microglia expression profile in developing white matter. BMC Res Notes 12, 1–6.

Sylvester, J., Purdie, G., Slee, M., Gray, J.X., Burnet, S., and Koblar, S. (2013). Muscle-specific kinase antibody positive myaesthenia gravis and multiple sclerosis co-presentation: A case report and literature review. J Neuroimmunol 264, 130–133.

Tabata, H. (2015). Diverse subtypes of astrocytes and their development during corticogenesis. Front Neurosci 9, 629.

van der Poel, M., Ulas, T., Mizee, M.R., Hsiao, C.-C., Miedema, S.S.M., Adelia, Schuurman, K.G., Helder, B., Tas, S.W., Schultze, J.L., et al. (2019). Transcriptional profiling of human microglia reveals grey– white matter heterogeneity and multiple sclerosis-associated changes. Nature Communications 10, 1139.

Ventura-Antunes, L., Mota, B., and Herculano-Houzel, S. (2013). Different scaling of white matter volume, cortical connectivity, and gyrification across rodent and primate brains. Front Neuroanat 7, 3.

Viganò, F., Mobius, W., Götz, M., and Dimou, L. (2013). Transplantation reveals regional differences in oligodendrocyte differentiation in the adult brain. Nat. Neurosci. 16, 1370–1372.

Wang, H., Moyano, A.L., Ma, Z., Deng, Y., Lin, Y., Zhao, C., Zhang, L., Jiang, M., He, X., Ma, Z., et al. (2017). miR-219 Cooperates with miR-338 in Myelination and Promotes Myelin Repair in the CNS. Devcel 40, 566–582.e5.

Wilton, D.K., Dissing-Olesen, L., and Stevens, B. (2019). Neuron-Glia Signaling in Synapse Elimination. Annu. Rev. Neurosci. 42, 107–127.

Xiao, L., Ohayon, D., McKenzie, I.A., Sinclair-Wilson, A., Wright, J.L., Fudge, A.D., Emery, B., Li, H., and Richardson, W.D. (2016). Rapid production of new oligodendrocytes is required in the earliest stages of motor-skill learning. Nat. Neurosci. 19, 1210–1217.

Young, K.M., Psachoulia, K., Tripathi, R.B., Dunn, S.-J., Cossell, L., Attwell, D., Tohyama, K., and Richardson, W.D. (2013). Oligodendrocyte dynamics in the healthy adult CNS: evidence for myelin remodeling. Neuron 77, 873–885.

Zeisel, A., Hochgerner, H., Lonnerberg, P., Johnsson, A., Memic, F., van der Zwan, J., Haring, M., Braun, E., Borm, L.E., La Manno, G., et al. (2018). Molecular Architecture of the Mouse Nervous System. Cell 174, 999–.

Zhang, Y., Chen, K., Sloan, S.A., Bennett, M.L., Scholze, A.R., O’Keeffe, S., Phatnani, H.P., Guarnieri, P., Caneda, C., Ruderisch, N., et al. (2014). An RNA-sequencing transcriptome and splicing database of glia, neurons, and vascular cells of the cerebral cortex. J. Neurosci. 34, 11929–11947.

Zhang, Y., Sloan, S.A., Clarke, L.E., Caneda, C., Plaza, C.A., Blumenthal, P.D., Vogel, H., Steinberg, G.K., Edwards, M.S.B., Li, G., et al. (2016). Purification and Characterization of Progenitor and Mature Human Astrocytes Reveals Transcriptional and Functional Differences with Mouse. Neuron 89, 37–53.

