## Supplemental Document for "Microenvironment Impacts the Molecular Architecture and Interactivity of Resident Cells in Marmoset Brain"

**This PDF file includes:**

Materials and Methods: page 2 – 14
Supplementary Text: page 15 – 21
References: page 22 – 27
Figures S1 to S30: page 28 – 82
Data S1 to S4 title: page 83

#### Materials and Methods

##### Key Resources table

| REAGENT or RESOURCE | SOURCE | IDENTIFIER |
|---|---|---|
| **Biological Sample** | | |
| Marmoset | NINDS/NIH | Callithrix jacchus |
| **Critical Commercial Assays** | | |
| Chromium Single Cell 3' Library & Gel Bead Kit v2 | 10x Genomics | PN-120237 |
| Chromium Single Cell A Chip Kit | 10x Genomics | PN-120236 |
| Chromium Single Cell 3' GEM, Library & Gel Bead Kit v3 | | PN-1000075 |
| Chromium Single Cell B Chip Kit | 10x Genomics | PN-1000153 |
| Chromium i7 Multiplex Kit | 10x Genomics | PN-120262 |
| **Deposited Data** | | |
| Hammond et. al. 2019 | (Hammond et al., 2019) | GEO: GSE121654 |
| Marisca et. al. 2020 | (Marisca et al., 2020) | GEO: GSE132166 |
| Marques et. al. 2016 | (Marques et al., 2016) | GEO: GSE75330 |
| Zeisel et. al. 2018 | (Zeisel et al., 2018) | http://mousebrain.org/ |
| Zhang et. al. 2014 | (Zhang et al., 2014) | GEO: GSE52564 |
| Polioudakis et. al. 2019 | (Polioudakis et al., 2019) | dbGaP: phs001836 |
| Lake et. al. 2018 | (Lake et al., 2018) | GEO: GSE97930 |
| Habib et. al. 2017 | (Habib et al., 2017) | GEO: GSE104525 |
| Zhang et. al. 2016 | (Zhang et al., 2016) | GEO: GSE73721 |
| Jäkel et. al. 2019 | (Jäkel et al., 2019) | GEO: GSE118257 |
| Marmoset Gene Atlas | (Shimogori et al., 2018) | https://gene-atlas.brainminds.riken.jp/ |
| Marmoset Brain Mapping | (Liu et al., 2020; 2018a) | https://marmosetbrainmapping.org/ |
| **Software and Algorithms** | | |
| R (v3.6.1 2019-07-05) | | https://cran.r-project.org/bin/macosx/ |
| Cellranger (v3.0.2) | 10x Genomics | https://www.10xgenomics.com/ |
| seurat (v3.1.5) | | https://github.com/satijalab/seurat |
| DoubletFinder (v2.0.2) | | https://github.com/chris-mcginnis-ucsf/DoubletFinder |
| SoupX (v1.4.5) | | https://github.com/constantAmateur/SoupX |
| harmony (v1.0) | | https://github.com/immunogenomics/harmony |
| monocle3 (v 0.2.0) | | https://github.com/cole-trapnell-lab/monocle3 |
| gprofiler2 (v0.1.9) | | https://cran.r-project.org/web/packages/gprofiler2/index.html |
| nichenetr (v 0.1.0) | | https://github.com/saeyslab/nichenetr |
| EWCE (v0.99.2) | | https://github.com/NathanSkene/EWCE |
| Ingenuity Pathway Analysis (IPA) (v01-16) | Qiagen | https://digitalinsights.qiagen.com/product-login/ |
| Fiji (v2.1.0/1.53c) | | https://imagej.net/Fiji/Downloads |
| **Other** | | |
| CjPCA website | This manuscript | https://cjpca.ninds.nih.gov |

##### Lead contact and materials availability

Direct requests for reagents and information to Daniel S. Reich and Jing-Ping Lin.

#### Method details

#### Animals

Our central nervous system marmoset cell atlas was generated from 2 healthy, 5.5-year-old common marmosets (*Callithrix jacchus*), one female (CJH01) and one male (CJR02). Staining was done with 4 healthy 4–6-year-old marmosets, two females (CJT11, CJV13) and two males (CJB10, CJD12). All marmosets were housed and handled with the approval of the Institutional Animal Care and Use Committee. On the day of imaging, marmosets were anesthetized by intramuscular injection of 10 mg/kg ketamine, intubated, and ventilated with a mixture of isoflurane and oxygen during *in vivo* MRI scans. MRI was performed on a 7T Bruker to generate a series of proton density-weighted images with a resolution of 0.15 x 0.15 x 1 $mm^3$ per voxel and a matrix of 213 x 160 x 36 per session (Sati et al., 2012). The images were used for volume reconstruction and anatomy identification. The volume was then used to create a custom-made brain holder by 3D printing (Ultimaker $2^+$) for each marmoset to guide ex vivo tissue sampling (Luciano et al., 2016). After each in vivo scan section, marmosets were weaned from 2% isoflurane, recovered with a lactate ringer injection subcutaneously, and returned to their original housing.

#### Single nucleus RNA sequencing (snRNA-seq)

##### Tissue dissection for nuclei isolation

On the day of tissue harvest, marmosets were deeply anesthetized with 5% isoflurane until all visible signs of breathing were no longer detected. Animals were transcardially perfused with ice-cold artificial cerebrospinal fluid (aCSF) for 5 min with a pump. Brains were removed from the skull and submerged into ice-cold aCSF, and after removal of meninges within the solution, were positioned in a custom-designed brain holder within 10 min post-perfusion. The brain was sectioned at 3 mm into 12–13 slabs in one step with a homemade blade-separator set in the solution. Each brain slab was transferred into a 6-well plate, submerged into RNAlater (RNAlater™ Stabilization Solution, AM7021, Invitrogen) with a homemade brain trap, and stored at 4°C overnight. The following day, brain slabs were positioned in 25 x 20 x 5 mm molds (Tissue-Tek® Cryomold®, 4557, Sakura Finetek) on ice to facilitate target sampling. Slabs were matched to MRI for each animal, and tissue annotation for gray (Liu et al., 2018a) and white matter (Liu et al., 2020) was informed by marmoset 3D MRI atlases V1 and V2. A cylinder of tissue 2 mm in diameter and 3 mm in height for each region (Figure 1B, S1) was collected with a tissue punch (EMS-core sampling tool, 69039-20, EMS). There were 5 white matter samples from temporal and parietal lobes that did not exactly match in the two animals, however they were paired in lobes of the brain and showed no significant differences in later analysis (Figure 1B, SS05, SS06, and SS08). The cylinders were ejected into PCR tubes filled with 100 μL of RNAlater and stored at −80°C.

#### Single-nucleus dissociation

Nuclei preparation was carried out as described (Matson et al., 2018), with minor modifications. Briefly, on the day of dissociation, tissue samples were thawed on ice, removed from solution, dabbed with Kimwipes to remove residual RNAlater, and placed in a 1 mL douncer tube (Dounce Tissue Grinder, 357538, Wheaton). Each tissue was homogenized in 500 μL of lysis buffer containing 0.1% Triton-X100 in low sucrose buffer (0.32 M sucrose, 10 mM HEPES, 5 mM $CaCl_2$, 3 mM MgAc, 0.1 mM EDTA, and 1 mM DTT in $ddH_2O$, pH8) with loose pestle 25 times and tight pestle 10 times. The homogenate was filtered through a 40-μm mesh (Falcon® 40 μm Cell Strainer, 352340, Corning) to a 50-mL Falcon tube on ice. An additional 5 mL of low sucrose buffer was used to rinse the douncer tube and cell strainer. The filtered homogenate was further mixed with a handheld homogenizer (VWR® 200 Homogenizer) at a speed of ~1000 rpm to brake nuclei clumps for 5 sec. After homogenization, a serological pipet filled with 12 mL of high sucrose buffer (1 M sucrose, 10 mM HEPES, 3 mM MgAc, and 1 mM DTT in $ddH_2O$, pH8) was placed underneath the lysate and disconnected from the pipettor, and the buffer was released from the serological pipette by gravity and set on ice. When most of high sucrose buffer was released to form a density layer underneath the homogenate, the serological pipet was retrieved along the wall of the Falcon tube gently, without disturbing the low-high sucrose interface. The Falcon tube was capped and placed in a swing bucket to be centrifuged at 3,200 rcf for 30 min at 4°C. At the end of spin, the supernatant was decanted quickly without tabbing, and 1 mL of resuspension buffer (0.02% BSA in 1X PBS, pH7.4) was added to the Falcon tube. Slow pipetting was employed to resuspend nuclei along the Falcon tube wall below the 5-mL mark to preserve nuclei integrity. Specifically, nuclei were rinsed off the wall in courses of 2 sec per trituration for 20 times total per tube. The Falcon tube was then capped and spun at 3,200 rcf for 10 min at 4°C. At the end of spin, the supernatant was removed by gently tabbing the tube until no visible liquid drop was left behind, and 200 μL of resuspension buffer was added to each sample to collect the nuclei. The nuclei suspension was filtered through a 35-μm mesh (Cell Strainer Snap Cap, 352235, Corning) twice and counted on a hemocytometer by trypan blue staining. During counting, the size and quantity of myelin and other debris were visually inspected under the scope, and the suspension was filtered 1–3 more times through the 35-μm mesh if necessary. Only round and dark-blue stained nuclei were considered of good quality and included in the final count.

**cDNA Library & Sequencing**

Most single nucleus libraries (38) were prepared using 10x Genomics Chromium Single Cell 3' Library & Gel Bead Kit v3, though 4 libraries were done in v2 chemistry following the manufacture's protocol. Briefly, nuclei suspensions were prepared as described above and diluted with resuspension buffer to achieve the desired concentration, and then loaded into Chromium Controller to generate Gel-beads in Emulsion (GEMs). For both cDNA amplification and library sample index PCR, 12 cycles were used. Most libraries were sequenced on Illumina Novaseq S2, but some used Illumina Miseq, Hiseq 2500, or Hiseq 4000, according to manufacturer's protocol; see Figure S4 and Table S1 for details.

**Alignment**

The raw sequencing reads were aligned to a marmoset genome assembly, ASM275486v1 (GCA_002754865.1). To build a reference package suitable for analyzing both unspliced pre-mRNA and mature mRNA in the nuclei, as well as to include sequences of mitochondrial genome, marmoset DNA sequence (FASTA) and annotation (GTF) files were acquired from the Ensembl release-95 and modified as follows. The complete mitochondrial sequence (NC_025586.1, GenBank) and its annotation (Wang et al., 2016) were manually added to the FASTA and GTF files. Next, a pre-mRNA GTF was made by replacing "transcript" with "exon" as the feature-type entry in the original GTF before making a reference package with CellRanger software (v3.0.2, 10x Genomics). This custom-built reference package was then used in CellRanger (version 3.1.0, 10x Genomics) to align sequencing reads for all samples. The option to estimate cell number automatically was used for most of the samples, unless otherwise specified (see Table S1 for details). A filtered cell barcode-to-gene feature matrix was generated from the software and used for downstream analysis (Figure S2A).

**Preprocessing and quality control**

The matrix was loaded to create an object in Seurat v3 (Butler et al., 2018; Stuart et al., 2019). Cells with <200 genes, >5000 genes, or >5% of counts mapped to mitochondrial genome were excluded. Genes observed in <5 cells were excluded. The filtered raw count matrix was then log normalized (ln(counts*100,000+1)) within each cell and scaled to account for differences in sequencing depth with Seurat. Next, DoubletFinder (McGinnis et al., 2019) was used to estimate and remove putative doublets to mitigate technical confounding artifacts in droplet-based sequencing data analysis. The top 3000 variable genes calculated by Seurat were used in linear dimension reduction (principal components analysis, PCA), and the top 30 PCA were used for clustering at low resolution (parameter = 0.4) to define crude cell types. These unsupervised clusters were used to provide a quick cluster annotation for homotypic doublet probability modeling in DoubletFinder. Doublet rate was estimated by fitting a linear equation over a multiplet rate table provided by 10x Genomics. The rate = (0.0008*cell.number + 0.0527)/100) was used to calculate a Poisson distribution with and without homotypic doublet proportion to generate low confidence (DF.found.1) and high confidence (DF.found.2) doublet annotation. Unless otherwise specified, pN = 0.25, pK = 0.005 and automatic doublet removal based on DF.found.2 annotation were used, as the first line of screening. In parallel, SoupX (Young and Behjati, 2020) was used for ambient RNA background correction. Taken from the output of the 10x pipeline (raw_feature_bc_matrix), the ambient RNA from empty droplets that contained <10 unique molecular identifiers (UMI) were profiled, and the "soup" contamination fraction was calculated for each cluster. Given that the nuclear transcriptome was profiled, genes that mapped to the mitochondrial genome could be considered as ambient input. Therefore, the top mitochondrial genes (species with >1000 accumulated counts across profiled empty droplets) were used to estimate the global contamination fraction and adjust the raw count matrix. Next, the cell barcode that passed the DoubletFinder was used as an index to subset the SoupX-corrected matrix to generate a new matrix as our downstream input (Figure S2A). For individual samples, a Seurat object was created, and the index labels (IL01_uniqueID, IL02_species, IL03_source, IL04_sex, IL05_ageDays, IL06_tissue.1 (coarse category), IL06_tissue.2 (developmental category), IL06_tissue.3 (fine category), IL07_location, IL08_condition, IL09_illumina, IL10_chemistry, IL11_batch, IL12_LMinDays, IL13_LMaxDays, IL14_dataset, IL15_annotation) were added to the metadata as cell attributes. For each sample, 90% of nuclei were randomly selected, then all 42 samples were merged for downstream comparison. The remaining 10% of the nuclei were set aside for classification assessment and validation.

#### Clustering and visualization

Clustering was performed using Seurat, iteratively with different parameter sets, to understand the data structure. A merged Seurat object was created from the 42 samples, and the aggregated raw count matrix was log-normalized and scaled again as stated above.

##### *Preliminary exploratory data analysis*

The top 3000 variable genes calculated by Seurat were used in PCA. In the first round of clustering and visualization, 100 principal components (PC) were computed and used for Harmony (v1)

(Korsunsky et al., 2019) to integrate different samples, specifically variability over the IL01_uniqueID attribute, with default setting (theta=2, lambda=1, sigma=0.1). The UMAP space and nearest neighbor analyses were calculated on the top 50 Harmony embeddings with resolutions from 0.4–1.2. Cell barcodes of a cluster of nuclei annotated as "low quality," which resided at the center of the 2D UMAP (H50), were recorded; these nuclei had a high percentage of reads mapped to the mitochondrial genome, low RNA counts and features, and/or expressed genes that mapped to multiple canonical markers of different cell types. No single set of parameters can adequately separate ~500K nuclei to identify subclusters from all major cell types simultaneously, as either over-splitting for low complexity cells (e.g., glia) or under-splitting for high complexity cells (e.g., neurons) would result. Therefore, a stepwise clustering approach was used, whereby major cell classes (neurons, oligodendrocytes, etc.) were first identified and then and divided into subclusters for each class.

*Level 1 quality control and analysis*

To divide nuclei into classes and facilitate artifact identification, nuclei were first classified using a set of parameters that does not highlight granular detail. In this round of clustering, only 50 PC for Harmony were computed to perform linear correction over IL01_uniqueID, as the elbow plot from the preliminary analysis showing the standard deviation stopped visually decreasing after top 50 PC. The top 5 Harmony-corrected embeddings (H5) were used for Seurat to learn the UMAP and find cell classes at a low resolution (0.2). Canonical cell-type markers (*PTPRC* for immune cells, *PDGFRA* for OPC, *MAG* for oligodendrocytes, *GFAP and SLC1A2* for astrocytes, *LEPR* and *CEMIP* for vasculature and meningeal cells, and *CNTN5* and *NRG1* for neurons) annotated 6 of the classes unambiguously. One cluster in the middle of the H5 UMAP had mixed expression of canonical markers, which suggested artifact. The "low quality" cell barcodes that were found from the H50 condition (defined above) were overlaid on the H5 UMAP, which exclusively highlighted the putative artifact cluster. These nuclei were removed from further analysis, although the original UMAP embeddings were maintained for plotting purposes (Figure 1 and S3).

*Level 2 quality control and analysis*

Nuclei that passed Level 1 QC were divided into 5 classes (MIC, OPC, OLI, VAS/AST, NEU) based on the H5 UMAP result. The astrocytes and vasculature/meningeal cells were pooled into a single class prior to subclustering to facilitate artifact identification. Shared features in this class of cells were potentially explainable by their close association at CNS barriers (blood–CSF interface, blood–brain interface, and the CSF–brain interface). For each class, log-normalization and scaling were repeated from the divided raw count matrix, the top 3000 variable genes were used for 50 PC computation, Harmony correction over IL01_uniqueID, UMAP learning, and clustering, as described above. The clustering resolution was iteratively increased from low to high (0–1.2), and clustering stability was tracked with clustree (v0.4.3) (Zappia and Oshlack, 2018). NEU Level 2 clustering stability was also tracked by calculating Jaccard index at resolution 2 (res.2, Figure S12B) (Polioudakis et al., 2019). Aided by the branch visualization provided by the clustree, a tentative resolution that was relatively stable was selected, then differentially expressed gene (DEG) analysis on the clusters found with this parameter set was performed. The expression pattern of the top-expressed genes for each cluster within and across classes were checked, and artifact clusters were manually imputed. Doublets tended to form small distinct clusters in the UMAP plots that branched early in the clustering tree analysis with low splitting resolution, had mixed canonical marker-gene expression, and had similar expression patterns to cells in other partitions; thus, these doublets could be easily spotted and removed. For putative doublets within each class, additional rounds of DEG analysis were performed as necessary. Each time nuclei were removed, basic normalization, scaling, Harmony, and UMAP learning were repeated. To control for over-splitting, for clusters that appeared to be a single pile in the 2D UMAP space but were annotated into >1 cluster, additional rounds of DEG analysis were performed to see if binary markers could be found to label them. In addition, clustering was projected onto a 3D UMAP space to ensure effects were not masked due to overcrowding in 2D. This strategy also helped to further elucidate cluster associations, aid decision-making with respect to groups of clusters that should be tested further, and spot potential gradient changes among clusters. If unique and or binary patterns could not be found in the current splitting resolution after these steps were performed, a step lower in resolution on the clustering tree was examined, and the analysis process was repeated. The following compound naming convention to label the 87 subclusters was used: general category in numeric order, major tissue or location contributor for each cell type, and binary marker combination where applicable.

*Preparation of objects for cross-cluster analysis*

Once the subclustering and UMAP embedding were finalized for each cell class, several annotated objects were created to facilitate downstream analysis and comparison. To enable cluster overview, compare global and local gene expression, and classify the 10% set-aside data, an object containing all 87 subclusters and 50 nuclei per cluster was prepared by random sampling (C50 object). For white and gray matter comparison, 4000 nuclei were randomly sampled from each tissue type and pooled into two objects, WM (24,000 nuclei, including fWM, tWM, pWM, aCC, pCC, OpT) and GM (20,000 nuclei, including fCTX, tCTX, pCTX, oCTX, and CgG).

*Data visualization*

Unless otherwise specified, gene expression values in the dot plots and heatmaps were averaged, mean-centered, and z-score-scaled (from -1.5 to +1.5, to which values below or above these levels were assigned). Dot size indicates the percentage of nuclei in the subcluster in which the gene was detected. Among the nuclei in which a given gene was detected, the expression level was mean-centered and scaled. For aggregated gene lists or gene module expression, a relative color scheme was used to indicate the level of expression, from low to high. For dendrogram creation, the top 50 enriched genes calculated in Level 1 analysis were used to calculate Euclidean distances, using "hclust(dist())" functions in basic R. To aid cluster tracking, branches of the dendrogram were reordered and colored to show the origin of cell classes while retaining the tree structure.

**Pseudotime analysis**

Monocle3 (v0.2.0) was used to construct nuclei trajectories based on transcriptomic distance (Cao et al., 2019). The OLI Seurat object with finalized UMAP from Level 2 analysis was converted to a Monocle object. All index labels and cell attributes, cluster assignment, and UMAP embeddings were transferred. A partition was then assigned for each nuclei by the cluster_cells() function, and a principal graph was fit within each partition by the learn_graph() function. From the principal graph, Monocle3 defined a unitless transcriptome progression along the learned trajectory as "pseudotime." The distance between two given points along the trajectory path indicates the amount of expression change required to connect the ends. The starting point of pseudotime is self-defined by the order_cells() function. Based on prior knowledge (Xiao et al., 2016), the node at the side of *ENPP6*$^{high}$ oligodendrocyte cluster was selected as the starting point. To visualize gene expression dynamics along pseudotime, the plot_genes_in_pseudotime() function was used to fit a spline using the following trend formula: "~ sm.ns(Pseudotime, df=3)". The calculated pseudotime value was extracted for further analysis as indicated in the figure legend (cds@principal_graph_aux@listData[["UMAP"]][["pseudotime"]]).

**Gene module analysis**

Monocle3 was used to find and group genes by similarity along the learned principal graph. Genes that passed Moran's *I* statistic spatial test (< 5% FDR) over the *k*-nearest neighbor graph (Knn, k=25), or trajectory learned principal graph (PG) by Monocle3 graph_test() function, were used for module assignment. Genes were grouped into modules identified in each type of graph test by the find_gene_modules() function with resolution of $10^{-3}$. The list of genes of each module was then aggregated and added back to the Seurat object through the AddModuleScore() function and visualized in Seurat v3. Genes that mapped to the mitochondrial genome were dropped before performing gene ontology and pathway analysis. See Table S2 for the full list.

**Gene ontology (GO) and pathway analysis**

The list of genes from the selected modules and/or DEG, discovered as stated above, were used for various pathway analysis. The GO analysis for marmoset was performed by gprofiler2 (v0.1.9) (Raudvere et al., 2019) with the gost() function. The database for "cjacchus" was used, electronic GO annotations (IEA) was included, g:SCS threshold was used for multiple testing correction as suggested by gprofiler2. Three major subontologies — Molecular Functions (MF), Biological Process (BP), and Cellular Component (CC) — were included in analysis. Additional annotations from KEGG and HP database were included when available. Terms that passed a significance cutoff of $p$=0.05 after correction were filtered at the following criteria in case of overcrowding. The parent terms were removed if child terms from the same branch were present in the same list, and if the term had at least one parent term in the database prioritized, as terms lower in each branch are usually more specific and informative. For terms that passed filtering, the corrected p-value and fold enrichment were plotted. The fold enrichment was calculated as follows: (intersection_size/query_size)/(term_size/effective_domain_size).

**NicheNet ligand-receptor-target analysis**

Potential intercellular communication in WM and GM was modeled using nichenetr (v0.1.0) (Browaeys et al., 2020). The cross-partition objects for WM and GM generated as described above were used for this analysis. Bioinformatic resources and protocols were modified from https://github.com/saeyslab/nichenetr. Briefly, NicheNet studies intercellular communication computationally by leveraging known ligand-to-receptor and receptor-to-target relationships in its database. It allows prediction of interactions between ligands expressed by "sender" cells and receptors expressed by "receiver" cells, and models how these interactions might drive gene expression changes in cells of interests (target DEG in the receivers).

We hypothesized that differences between the white and gray matter milieus might partially explain tissue-type-specific subpopulations in microglia, OPC, and astrocyte classes. Therefore,

NicheNet was used to test if transcriptome changes between subclusters could be explained by environmental signals from nearby cells. White and gray matter differentially enriched microglia, OPC, and astrocyte subpopulations were defined as receivers in each test, and DEG between MIC1 and MIC3, OPC1 and OPC3, and AST1 and AST3 were derived. DEG were filtered at adjusted p-value (<0.05) and absolute log (ln) fold-change (>0.25). Potential senders were defined from clusters with >50 nuclei in the same tissue type as each receiver, and genes detected in >10% of the nuclei in a cluster were kept for further analysis. Preconstructed databases were downloaded for ligand target martix, ligand receptor database, and weighted networks. Gene names for these databases were built with human data, therefore human genes with one-to-one orthologs were translated to marmoset gene names with BioMart. The weighted ligand-receptor-target (LRT) matrix was thereby constructed, with weighting factors implemented so that informative data sources maximized prediction accuracy in the final model. The list of sources used to build this database and the method to calculate the weighted scores have been specified (Browaeys et al., 2020). After "expressed ligands" were defined for senders and "expressed receptors" for receivers (>10% detection rate), the existence of ligand-target pairs was established. The ligand-target pairs were ranked based on the presence of the target genes (defined by the receptor-target database) in the calculated DEG using the predict_ligand_activities() function. The filtered DEG present in the top 200 predicted target genes per ligand were kept for further ligand-target analysis.

To aid visualization of this complicated intercellular interaction, Circos plots were generated. First, the lists of ligand-target pairs found in WM and GM were compared and divided into 3 categories (GM, WM, and shared), and a Venn diagram was generated for each type of receiver (microglia, OPC, and astrocytes; Figure 7E, H, K). The unique ligand-target pairs for each environment were plotted in enlarged Circos plots, and the shared ligand-target pairs in smaller Circos plots, for categorical visual reference (Figure 7E, H, K, bottom panels). The shared Circos plots were also enlarged to aid visibility of individual genes (Figure S30A, C, D, F, G, I). Since any given ligand might be expressed by >1 cell type, each ligand was assigned to the cell cluster that ranks highest in the product of detection rate (%) and expression level (z-score scaled). Since the probability is low for any ligand to be assigned to a particular cell type with this strategy, it is sufficient to map intercellular interaction qualitatively and categorically (see Table S4 for full report). The senders by Level 1 classes (MIC, OPC, OLI, AST, VAS, NEU) were colored, and pie charts tabulating the proportion of unique (Figure 7E, H, K) and shared (Figure S30A, C, D, F, G, I) ligand-target pairs were generated. For each Circos plot, up to top 100 weighted ligand-target interactions were presented to limit overcrowding. For shared ligand-target pairs, inter-categorical agreement was calculated and presented in Sankey diagrams using the networkD3 (v0.4) package.

#### Gene set enrichment analysis

Themes were collected from the following sources, after which the aggregated score was calculated by Seurat v3 AddModuleScore() function (Tirosh et al., 2016). Gene groups included ion channels, scavenger receptors (SCAR), and histocompatibility complex (HLA) from the HUGO Gene Nomenclature Committee (HGNC). Cell-cycle genes were pulled from the built-in gene list in Seurat (cc.genes.updated.2019) for S and G2M phases. Genes enriched in G0G1 phase (Feldman et al., 2019) and the list of human transcription factors (Lambert et al., 2018) were informed from the literature. Neurological disorder-associated genes were acquired from the database curated in the Ingenuity Pathway Analysis (IPA) software. See Table S4 for the full gene lists.

#### Expression-Weighted Cell-type Enrichment (EWCE) analysis

Cellular phenotypes of neurological disorders were calculated by EWCE analysis (Skene and Grant, 2016). Briefly, the expression of a list of *n* genes associated with a disease or disease category was compared with those in 100,000 randomly selected lists of *n* genes from the background. The proportional expression of genes associated with each cell type was calculated to compute the probability of enrichment. Tested disease/disease categories were: cognitive impairment, psychological disorder, seizures, schizophrenia spectrum disorder, autism spectrum disorder or intellectual disability, migraine, Alzheimer disease or frontotemporal dementia, Zellweger syndrome, abnormality of meninges, neurological disorder, Huntington disease, organic mental disorder, encephalitis, multiple sclerosis, cerebrovascular dysfunction, stroke, white matter abnormality, Charcot-Marie-Tooth disease, glioma formation, parkinsonism, and central nervous system tumor. The significance of cell-type enrichment was defined by $p<0.05$ after Benjamini-Hochberg correction.

#### Cross-cluster comparison and validation

##### *Comparison between marmoset subclusters*

The C50 object was used to assess transcriptomic similarity across all 87 subcluster pairs. The expression levels of all genes within each subcluster were normalized and averaged before calculating the linear correlation. The lm() function was used in R, and the adjusted $r^2$ values were extracted for heatmap plotting. Similarly, the transcriptomic distances between all subcluster pairs were assessed by counting the number of DEG, both increased and decreased, between them. DEG were filtered by their log (ln) fold change (>0.25) and detection frequency (detected in ≥10% of nuclei).

*Comparison between clusters from different species*

Deposited data from zebrafish, mouse, and human were reanalyzed. The top 3000 variable genes were used to calculate 50 PCA and harmonized over sample ID if available in the deposited data. Gene names for each species were translated to human gene names using the one-to-one orthologs index with BioMart. The expression levels of all humanized genes within each compared cluster were normalized and averaged before calculating the Pearson's correlation coefficients, which were used for heatmap plotting.

*Comparison between cleaned classifiers and semi-cleaned 10% set-aside data*

To assess reproducibility of our derived subclusters, the C50 object was used as an unbiased classifier to annotate the 10% of nuclei that had been set aside a priori, as described above. A total of 61,852 nuclei were compared. The nuclei were intentionally over-split using the top 5000 variable genes (maximum gene number detected per nuclei) to calculate 100 PC, harmonized over IL01_uniqueID labels. All 100 Harmony embeddings were used to compute UMAP and nearest-neighbor distances with extremely high resolution (12; normal suggested resolution range is 0.4–1.2). A total of 140 clusters were found, and the expression levels of genes within each cluster were normalized and averaged before calculating the Pearson's correlation coefficients across each pair of subclusters in the two datasets, which were used for heatmap plotting.

#### Staining

Sections used for histology were archival formalin-fixed, paraffin-embedded (FFPE) contained in an in-house marmoset tissue library. Serial sections were cut at 5 μm from brain and spinal cord tissue blocks from each animal using a Leica RM2235 Manual Rotary Microtome. Sections were mounted onto Superfrost+/Colorfrost+ microslides (Daigger, 75 x 25 mm, #EF15978Z) and stored at room temperature. Before staining, sectioned slides were deparaffinized with xylene and rehydrated with ethanol in decreasing concentrations. Hematoxylin & eosin staining was subsequently performed. Hematoxylin (basophilic) stains nucleic acids and nuclei purple, whereas eosin (acidophilic) stains cytoplasmic components of the cell pink. For hematoxylin staining: slides were dipped one-by-one in hematoxylin (Leica, 100% Surgipath SelecTech Hematoxylin 560MX, 3801575) for 1 min and immediately placed in running tap water to stop the reaction and rinse off excess stain. Slides were then dipped one-by-one for 30 sec in Define solution (Leica, Surgipath SelecTech Define MX-aq, 3803598) to reduce the intensity of hematoxylin and immediately placed in running tap water to stop the reaction. Sections were dipped one-by-one in Blue Buffer solution (Leica, Surgipath SelecTech Blue Buffer 8, 3802918) for 1 min to change tissue color to blue. This reaction was stopped by placing slides in 80% ethanol for 1 min. For eosin staining: slides were dipped together for 30 sec in eosin (Leica, Surgipath SelecTech Alcoholic Eosin Y 515, 3801615) and put in 100% ethanol for 1 min, 3 times. Coverslips were mounted on slides right away using VectaMount Permanent Mounting Medium (Vector Laboratories, #H-000-60).

**Microscopy and cell quantification**

On hematoxylin & eosin-stained slides from each animal, boxes were drawn around each 2-mm area of interest in brain and spinal cord. Each region was imaged at 10X magnification with a Nikon Eclipse Ci microscope. The number of cells in each area of interest was counted using Fiji ImageJ. A color image threshold of 0–165 was chosen to highlight an optimal number of cells and limit the number of falsely identified cells. Using the Analyze Particles function, the number of cells at the chosen threshold was counted automatically, with the pixel size range set to 20–200 and the circularity range to 0.25–1.00 to exclude as much background noise and linear particles as possible. After the automatic counting, any falsely identified cells were manually deleted, and a new count was saved. Additional cells not detected by the automatic counter were manually added using the Cell-Counter plugin. A final image with automatic and manual cell count markers was saved, and the total number of cells (including manual deletions and additions to the automatic count) was recorded. Cell counts were normalized to the imaged area to get density of the nuclei per tissue type in each animal. The averaged nuclei density per $mm^2$ in each tissue type was then quantified. To estimate the initial number of nuclei for single-nucleus sequencing per cylinder of 2 mm diameter and 3 mm height ($V=3\pi$ μL), the averaged 2D density measured from hematoxylin & eosin-stained sections was used after multiplication by section thickness (5 μm). The percentage of nuclei recovered after Level 1 quality control for each sample was then plotted, each circle representing the percentage of one sample (Figure S1E).

#### Supplementary Text

- **Related to Method (pipeline, quality control)**

The abundance of oligodendrocytes and neurons was correlated across tissue type (Figure S3D, left), such that more oligodendrocytes were found in cerebral white matter (hereafter denoted "**WM**") and more neurons in cortical gray matter (hereafter denoted "**GM**"). We found that only a modest number (~10.5% median abundance of total cells in WM, Figure S3E) of neurons were present in WM samples without the need for pre-selection, indicating the precision of image-guided tissue sampling. By contrast, deep gray matter (caudate nucleus and thalamus), midbrain, and hindbrain regions (hereafter denoted "**other**"), had cellular composition intermediate between WM and GM (Figure S3D–E). Interestingly, there was a positive correlation between the abundance of microglia and oligodendrocyte progenitor cells (OPC) across tissue types (Figure S3D, right), with about 3-fold higher microglia and 2-fold higher OPC density in WM than GM (Figure S3E). Additional rounds of quality control (QC) and manifold learning constituted Level 2 analysis (Methods), where the 6 major cell classes were further grouped into 87 subclusters (Figure S5B). The pairwise linear correlation of all gene expression levels (Figure S6A) and the number of differentially expressed genes (DEG) between subcluster pairs (Figure S6B) were calculated for all 87 subclusters and presented as a heatmap. We found OPC subclusters generally expressed more "neuronal genes" (*CADPS*, *RIMS2*, *DLGAP1*, *NRXN3*, *STBP5L*) compared to other glia and vasculature related cells (Figure S5–6).

To assess bias in our manual, prior knowledge-derived annotation of nuclei classes and subclusters, we performed direct classification of a randomly selected 10% of nuclei that passed our standard preprocessing quality control pipeline and were then set aside a priori. We intentionally over-clustered the set-aside data without parameter optimization (Figure S7A). We inspected basic features of the 142 resulting subclusters (cj10_1–141, Figure S7B) and performed a Pearson's correlation analysis with expression levels in 50 nuclei randomly selected from each of our 87 knowledge-derived subclusters. The coefficients (Pearson's r) were rendered as a heatmap, highlighting correspondence between knowledge-derived and directly determined clusters (Figure S7C). Through this analysis, we were able to assign cluster identities, discover artifact clusters (high similarity with >1 major cell class, low number of detected genes, or high content of mitochondrial genes), and index new clusters in the set-aside dataset (Figure S7D). We conclude that direct classification of nuclei recapitulates the knowledge-derived atlas.

- **Neurons**

A total of 248,091 nuclei yielded 50 NEU subclusters (Figure S9A–B). We intentionally subclustered neurons at relatively low resolution to facilitate tracking of spatial origin. It is very likely that further splitting some of the NEU subclusters would yield more subtypes, and deeper analysis of this dataset should allow correspondence with ongoing efforts to classify and name neocortical cell types (Yuste et al., 2020). Conserved and evolutionarily diversified neuronal cell types between human, marmoset, and mouse have been discussed (Bakken et al., 2020; Hodge et al., 2020; Krienen et al., 2020), and detailed neuronal cell-type annotation of the adult and developing mouse nervous systems are accumulating (Gokce et al., 2016; Habib et al., 2017; 2016; Kozareva, 2020; La Manno, 2020; Rosenberg et al., 2018; Sathyamurthy et al., 2018; Tasic et al., 2016; Zeisel et al., 2018; 2015). For the marmoset atlas presented here, we used only the top 10 Harmony embeddings for subclustering and UMAP analysis, which allowed sufficient granularity to show transcriptomic similarity across clusters while facilitating subcluster tracking. We highlight well-studied markers, cluster annotations, and sampling site contribution to facilitate cross-database comparison (Figure S8A, S8D, S9C).

Nuclei in the UMAP were first colored by sampling site, which demonstrated, as expected, that gray matter was the major contributor (Figure S8). There was relatively high consistency in neuronal composition across GM, with some interesting exceptions, particularly occipital cortex (oCTX) and cingulate gyrus (CgG). We next identified the major tissue contributors for each subcluster by inspecting a heatmap showing the relative contribution of each sampling site and then colored the UMAP accordingly (Figure S8). This revealed distinct spatial segregation for neurons, whereby distinct NEU subclusters from some structures, particularly hippocampus (NEU10, 18, 31, 46–48), posterior corpus callosum (NEU08, 09, 19), caudate (NEU20, 21), and cerebellum (NEU23–29), remained close to one another in UMAP space. On the other hand, NEU subclusters (NEU01–07, 12) from midbrain and hindbrain were often mixed. For GM, 12 clusters were uniformly shared (NEU11, 14, 15, 17, 33, 38–42, 44, 45), 2 clusters were unique to oCTX (NEU36, 37), and 6 subclusters either lacked representation from oCTX (NEU34, 35, 50), CgG (NEU43), or temporal/parietal/occipital cortex (NEU49), or else included a contribution from hippocampus (NEU32).

We also colored nuclei on the UMAP plot by vesicular glutamate transporters (VGLUT; *SLC17A7*, *SLC17A6*, *SLC17A8*) expression, including subclusters with minimum 10% detection rate of VGLUT transcripts (Figure S8B, S9E). In general, *SLC17A7*$^{+}$ glutamatergic neurons were distinct from neurons expressing GABAergic genes (*GAD1*, *GAD2*, *SLC6A1*), however, 2 subclusters of *SLC17A6*$^{+}$ neurons (NEU12, 22) had low GABAergic gene expression (Figure S9E). We observed that subclusters with mixed excitatory and inhibitory neurotransmitter expression often had higher expression of other neurotransmitter genes as well. Neurotransmitter expression scores were calculated by aggregating the expression of a list of genes, as follows: dopaminergic (*TH*, *DDC*, *SLC6A3)*, noradrenergic (*TH*, *DCC*, *DBH*, *SLC6A2*), adrenergic (*TH*, *DDC*, *DBH*, *PNMT*), serotonergic (*TPH2*, *DDC*, *SLC6A4*), cholinergic (*CHAT*, *SLC5A7*), and glycinergic (*SLC6A9*, *SLC6A5*). Only 1 *SLC17A7*$^{+}$/*SLC17A8*$^{+}$ double-positive subcluster was found (NEU50), but 5 clusters from the midbrain and cerebellum were *SLC17A6*$^{+}$/*SLC17A7*$^{+}$ double positive (NEU01, 26–29) (Figure S9E).

Cortical excitatory neurons (primarily VGLUT1$^{+}$) were arranged onto a continuous path in the UMAP plot, corresponding to lamination (L2–L6, NEU33–45), as previously reported in mouse and human (Hodge et al., 2020). This indicates similarity in the transcriptomes of neurons that reside in adjacent laminae. As in mouse and human studies, expression of *STAB2* (L2–6), *LAMP5* (L2/3), *RORB* (L4), and *THEMIS* (L5/6) anchor the ends and follow the progression of this path. The expression of genes enriched in each or a subset of laminae has been examined by ISH on P0 marmoset brain in the Marmoset Gene Atlas (Shimogori et al., 2018). We therefore matched the morphology of a P0 Nissl-stained marmoset brain in coronal section to a 3D T2*-weighted MRI from an atlas (Marmoset Brain Mapping) and cross-referenced the coronal slice to a sagittal reformation of the same MRI data. Sections adjacent to the Nissl-stained slide were used to visualize expression of selected markers. Genes enriched in each subcluster progressed along the UMAP path of excitatory neurons from NEU33 to NEU45, corresponding to an outer-to-inner lamination trajectory in the ISH images (Figure S10A). *RASGFR2* (L2, NEU33), *RORB* (L4, NEU34–41), *FEZF2* (L4–L6, NEU40, 41, 44, 45), and NR4A2 (L6, NEU43) are enriched in subclusters found in all cortical samples. In contrast, *FOXP2* (L4–6, NEU34, 35, 38, 40, 44, 45) and *PDE1A* (L2–6, NEU34, 35, 28–45) were absent from the L4 clusters of occipital cortex. In addition, *ADAMTS17* was exclusively detected in NEU37 and uniquely labeled L4 of primary visual cortex (Figure S10B).

Interestingly, 2 clusters (NEU30, 31) of neurons from the hippocampus and caudate were found to be closely associated with L2/3 cortical excitatory neurons (NEU32, 33) on the UMAP, suggesting a degree of similarity. Though these neurons differed in the neurotransmitter secretion (*GAD2* for NEU30 and *SLC17A7* for NEU 31–33, Figure S8A–B) or reception (dopamine receptor for NEU30, PG.m19; Figure S11B), the expression of *LAMP5* and *KIAA2012* was shared across all 4 clusters (NEU30–33, Figure S9C). There was evidence of neurogenesis (*TOP2A*$^{+}$) in the hippocampus (NEU47), and 2 postmitotic neuron progenitor-like subclusters (*NKX2-1*$^{+}$) were found near the caudate nucleus (NEU20, 21). Neurons from the LGN (NEU02, 03) were enriched with genes that regulate pigmentation (PG.m16; Figure S11B), and *GATA3*$^{+}$ neurons (NEU05–08) were enriched with genes regulating mast cell differentiation (PG.m23; Figure S11B). As with astrocytes (see below), cerebellar neurons (NEU23–29) were dramatically separated from other neurons in the UMAP. Informed by a cerebellum dataset from mouse (Kozareva, 2020), we annotated Golgi cells (NEU23), Purkinje cells (NEU24), molecular layer interneurons (MLI, NEU25), granule cells (NEU26–28), and unipolar brush cells (UBC, NEU29). Genes enriched in cerebellar neurons are involved in inner ear receptor cell function (PG.m10; Figure S11B), which is important for sensing balance (Ango and Reis, 2019).

- **Microglia**

We detected twice as many genes in microglia subclusters (MIC1–3) as in PBMC1 (immature B cells) and PBMC2 (myeloid lineage). Dividing myeloid (PBMC3) and differentiated myeloid cells (macrophage, PBMC4) had similar numbers of genes detected as microglia (Figure S13B). For PBMC, only PBMC4 and, to a lesser extent, PBMC3, were consistently found across brain regions (Figure 2A). PBMC1 and PBMC2 were exclusively found in cervical spinal cord, probably due to contribution from blood in the spinal meninges, which (unlike brain meninges) were not stripped prior to tissue processing. The expression levels of genes shared and differentially enriched across immune nuclei are indicated by color in the dot plot (Figure 2C) and UMAP scatter plots (Figure S13D). The relative expression of the same genes in the MIC class (g-MIC), relative to other cell classes, is referenced as a heatmap bar above the dot plot (see Figure S13E for the full heatmap). The cluster-defining markers of WM-microglia (MIC3) are known to be important in inflammasome signaling (*NLRP1*) (Yap et al., 2019), microglia-astrocyte interactions (*LAMC3*) (Biswas et al., 2017), suppressed neuroblastoma malignant phenotype (*B4GALNT3*), and induced immune response in experimental colitis (*SLC15A1*, *PEPT1*) (Figure 2C).

- **Oligodendrocyte progenitor cells**

The OPC2 subcluster was almost absent in WM tissue and was more prevalent in the deep gray matter, midbrain, and brainstem regions (median abundance of ~11.7%, Figure 3C). OPC2 reside between OPC1 and OPC3 in UMAP space and have a gene-expression profile that is more like OPC1 than OPC3 (Figure 3B). Interestingly, *CDH8*, which was found to be enriched in OPC2, exhibits a strictly spatiotemporally regulated expression pattern during development. Specifically, in mouse E13.5, *Cdh8* is selectively expressed in progenitors at the intermediate zone and proliferative zone of ganglionic eminences. Along the migratory path of the cortical interneuron progenitors, *Cdh8*$^{+}$ cells are detected at the subventricular zone and subplate for the pallium, and striatum for the sub-pallium later at E18.5. Given that cortical neurons and OPC are derived from the same radial glial progenitor lineage, these results lead us to speculate that OPC2 is a subpopulation that retains the transcription program reflective of its developmental origin.

A relatively high proportion of OPC4 was found in the midbrain, pons, and spinal cord. This subcluster has higher gene-expression similarity with WM than GM OPC, and both OPC4 and WM OPC are enriched in genes that encode for scavenger receptors (Figure 3B). Interestingly, OPC4 exhibit several features that were commonly enriched in immune cells, such as phagocytosis (*COLEC12*), innate immunity (*APOE*, *C3*), and antigen presentation (MHCI and MHCII gene sets, *CD74*). These results support the idea that some OPC have an immune function (Falcão et al., 2018; Kirby et al., 2019).

- **Oligodendrocytes**

As a specific example of a gene differentially expressed across OLI subclusters, we examined *TENM2* (labels OLI1), a member of the teneurin family (4 paralogs) known to be important for axon guidance and target selection through homophilic interactions with neighboring cells (Beckmann et al., 2013). To identify potential targets for OLI1, we examined the expression of *TENM2* in other cell classes and found expression in most astrocytes and neurons, with notable absence (low expression in <5% of nuclei) from a subset of GABAergic (NEU05–08, *GATA3*$^{+}$) and cerebellar (NEU24–25, *MEGF10*+) neurons (Figure S20E). This suggests a potential difference in the extent to which different neuron populations are targeted for myelination by the teneurin pathway.

Another gene, *ENPP6* (labels OLI1–3), is a marker of newly differentiating oligodendrocytes (Xiao et al., 2016; Zhang et al., 2014), the protein product of which can be immunohistochemically detected in the mouse brain starting from postnatal day 2, preceding MAG and MBP expression. In marmoset, the detection of newly synthesized *ENPP6* transcript was high in OLI1–3 nuclei but dropped sharply in OLI4 (Figure 4B, 4D), a contrast from mouse oligodendrocytes, in which two peaks of *ENPP6* expression were observed — one at the younger end of newly formed oligodendrocytes and the other in myelin-forming, mature oligodendrocytes (Figure S22B–C). This two-peak pattern is unique to mouse and was seen in our reanalysis of several single-cell datasets from brain and spinal cord (Russ et al., 2020; Sathyamurthy et al., 2018). In human oligodendrocytes (Jäkel et al., 2019), whether there are two *ENPP6* peaks was not clear, as nuclei were arranged into a near-complete circle (Figure S21C).

- **Astrocytes**

We found that common markers, such as *ALDH1L1* and *GLI3*, most effectively label the whole lineage of astrocytes across regions, including Bergmann glia (AST8) in the cerebellum (Figure S13C and S15E). *SLC1A2* is enriched in GM astrocytes and *GFAP* and *AQP4* in WM astrocytes. Interestingly, *DCC* (AST1 marker), a tumor suppressor, is involved in Netrin-1 (*NTN1*) signaling-mediated astroglial development during formation of the corpus callosum and hippocampal commissure (Morcom et al., 2020). Downregulation of *DCC* associates with malignancy, especially astrocytic tumors (Nakatani et al., 1998; Reyes-Mugica et al., 1997). *HS3ST3A1* (AST3 marker) is a GWAS hit associated with multiple diseases, including cancers (Nakano et al., 2012), infectious disease (Atkinson et al., 2012; Nguyen et al., 2018), HIV transmission (Joubert et al., 2010), and the level of WM hyperintensity among non-demented elders (Guo et al., 2020). Given that *HS3ST3A1* was almost exclusively detected in WM-enriched astrocytes but not other neural cells (Figure S15G), it is worth looking into the astrocytic contributions to the phenotype of these diseases in more detail. AST2 presented a mixed feature of AST1 and AST3; we concluded that it is most likely a transitioning subcluster instead of artifact, as *LAMA3* is exclusively detected in this population. *LAMA3* is an anti-Epithelial-to-Mesenchymal Transition (EMT) gene, and its expression is greatly decreased in invasive tumorigenic cells (Huang et al., 2007); however, its role in astrocytes is not clear. *GMPR* (AST4 and AST5 marker), a therapeutic target for Alzheimer's disease (Liu et al., 2018b), was elevated in the posterior cingulate astrocytes of Alzheimer's disease (Sekar et al., 2015) and leukocortical demyelinated GM lesion of multiple sclerosis (van Wageningen et al., 2020). *PAX3* is a marker that exclusively labels astrocytes in the cerebellum (AST6–8, Figure S15G), is highly expressed during embryonic development (Ogawa et al., 2005), and is overexpressed in glioblastomas (Xia et al., 2013). Moreover, multiple subcluster-defining genes, such as *EYA4* (AST6), *ACTN1* (AST7), and *CEMIP* (AST8), have been reported to be elevated in brain metastases and glioma (Li et al., 2020; 2018; Rodrigues et al., 2019).

- **Vascular, meningeal, and ventricular cells**

In addition to astrocytes, cells closely associated with CNS barriers (blood–CSF, blood–brain, and brain–CSF) were identified. A total of 13,057 such nuclei comprised 11 VAS subclusters (Figure S28A–B). Pericytes (Pericyte1 and Pericyte2), vascular endothelial cells (VE1–3), and vascular smooth muscle cells (VSMC) were largely consistent across brain regions (Figure S28C). A higher percentage of ependymal cells, which form a permissive interface between CSF and brain along the ventricular lining, was, as expected, identified in tissue samples that line ventricles (tWM, pCC, Cd, and cSC). The distribution of vascular and leptomeningeal cells (VLMC1–4, brain fibroblast-like cells) was sporadic and most highly detected in hindbrain (pons and cerebellum) (Figure S27A).

Subcluster markers were rather unique for ependymal cells and VSMC (Figure S27B, S28D, S28G), but some subclusters had overlapping features (He et al., 2018; Vanlandewijck et al., 2018; Zeisel et al., 2018). For example, pan-endothelial markers (*KIRREL*, *ATP10A*, *FLT1*) were expressed in Pericyte2, *PDGFRB*, and *NOTCH3* in both pericytes and VSMC (Figure S27B), and *PDGFRA* in VLMC1. Nuclei in VLMC1–3 were arranged into a continuous single island in UMAP space. VLMC4 expressed *SLC47A1*, suggesting that it represents arachnoid barrier cells (Zeisel et al., 2018). A less divergent transcriptome was noted for VLMC2 compared to the other VLMC subclusters, suggesting a transitional state (Figure S27B).

Gene module analysis (Figure S27C–F, S28E–F, Table S2) showed that VLMC2 was enriched in genes involved in "blood vessel morphogenesis" and "T cell proliferation" (Figure S27C). Pericyte2 and vascular endothelial cells (VE1–3) enriched for “angiogenesis” and “innate immune response” (Figure S27D), and ependyma for “cilium movement involved in cell motility” and “smoothened signaling pathway involved in dorsal/ventral neural tube patterning” (Figure S27E). As a positive control, terms like “regulation of cardiac muscle contraction by calcium ion signaling” and “negative regulation of vascular associated smooth muscle cell migration” were enriched in VSMC (Figure S27F).

- **Expression-Weighted Cell-type Enrichment (EWCE) analysis**

We reasoned that our CjPCA, derived from healthy marmosets, can be used to identify previously overlooked cellular contributors to human neurological disease. Toward this end, we examined the cellular enrichment of genes associated with a spectrum of disorders using Expression-Weighted Cell-type Enrichment (EWCE) analysis (Skene and Grant, 2016). As additional proof of concept, we used the original lists of genes categorized in the database without preselection and employed EWCE analysis to calculate cellular enrichment. We calculated fold-change, enrichment probability, and significance by comparison to gene expression in 100,000 randomly selected genes from the background (Figure S29). Genes associated with multiple sclerosis were enriched in microglia, and with “white matter abnormality” in microglia and oligodendrocytes. As described previously in a human study (Polioudakis et al., 2019), genes associated with autism spectrum disorder or intellectual disability were enriched in both excitatory and inhibitory neurons, and there was a remarkably similar profile for seizures and schizophrenia. Interestingly, genes related to migraine were also overrepresented in certain neuronal subclusters. Astrocyte contribution was highlighted in addition to the involvement of pyramidal neurons in Huntington’s disease, independently supporting reports of glial involvement in its pathogenesis (Gleichman and Carmichael, 2020). We observed the potential contribution of a subset of oligodendrocytes (OLI5 and OLI6) to parkinsonism, consistent with recent reports from postmortem brain transcriptomic data (Bryois et al., 2020). Although it affects the peripheral nervous system rather than the CNS, Charcot-Marie-Tooth disease mapped to oligodendrocytes, possibly due to shared gene expression between central and peripheral myelin. Finally, genes related to glioma were enriched in oligodendrocytes, immune cells, and vascular cells, but surprisingly not in astrocytes or OPC.

#### References


Ango, F., and Reis, Dos, R. (2019). Sensing how to balance. Elife *8*, 1029.

Atkinson, A., Garnier, S., Afridi, S., Fumoux, F., and Rihet, P. (2012). Genetic variations in genes involved in heparan sulphate biosynthesis are associated with Plasmodium falciparum parasitaemia: a familial study in Burkina Faso. Malar. J. *11*, 108–109.

Bakken, T.E., Jorstad, N.L., Hu, Q., Lake, B.B., Tian, W., Kalmbach, B.E., Crow, M., Hodge, R.D., Krienen, F.M., Sorensen, S.A., et al. (2020). Evolution of cellular diversity in primary motor cortex of human, marmoset monkey, and mouse. bioRxiv *573*, 2020.03.31.016972.

Beckmann, J., Schubert, R., Chiquet-Ehrismann, R., and Müller, D.J. (2013). Deciphering teneurin domains that facilitate cellular recognition, cell-cell adhesion, and neurite outgrowth using atomic force microscopy-based single-cell force spectroscopy. Nano Lett. *13*, 2937–2946.

Biswas, S., Bachay, G., Chu, J., Hunter, D.D., and Brunken, W.J. (2017). Laminin-Dependent Interaction between Astrocytes and Microglia: A Role in Retinal Angiogenesis. Am. J. Pathol. *187*, 2112–2127.

Browaeys, R., Saelens, W., and Saeys, Y. (2020). NicheNet: modeling intercellular communication by linking ligands to target genes. Nature Methods 2017 14:3 *17*, 159–162.

Bryois, J., Skene, N.G., Hansen, T.F., Kogelman, L.J.A., Watson, H.J., Liu, Z., Eating Disorders Working Group of the Psychiatric Genomics Consortium, International Headache Genetics Consortium, 23andMe Research Team, Brueggeman, L., et al. (2020). Genetic identification of cell types underlying brain complex traits yields insights into the etiology of Parkinson's disease. Nature Publishing Group *52*, 482–493.

Butler, A., Hoffman, P., Smibert, P., Papalexi, E., and Satija, R. (2018). Integrating single-cell transcriptomic data across different conditions, technologies, and species. Nat. Biotechnol. *36*, 411–420.

Cao, J., Spielmann, M., Qiu, X., Huang, X., Ibrahim, D.M., Hill, A.J., Zhang, F., Mundlos, S., Christiansen, L., Steemers, F.J., et al. (2019). The single-cell transcriptional landscape of mammalian organogenesis. Nature *566*, 496–502.

Falcão, A.M., van Bruggen, D., Marques, S., Meijer, M., Jäkel, S., Agirre, E., Samudyata, Floriddia, E.M., Vanichkina, D.P., ffrench-Constant, C., et al. (2018). Disease-specific oligodendrocyte lineage cells arise in multiple sclerosis. Nat Med *24*, 1837–1844.

Feldman, H.M., Toledo, C.M., Arora, S., Hoellerbauer, P., Corrin, P., Carter, L., Kufeld, M., Bolouri, H., Basom, R., Delrow, J., et al. (2019). Neural G0: a quiescent-like state found in neuroepithelial-derived cells and glioma. bioRxiv *8*, 446344.

Gleichman, A.J., and Carmichael, S.T. (2020). Glia in neurodegeneration: Drivers of disease or along for the ride? Neurobiology of Disease *142*, 104957.

Gokce, O., Stanley, G.M., Treutlein, B., Neff, N.F., Camp, J.G., Malenka, R.C., Rothwell, P.E., Fuccillo, M.V., Südhof, T.C., and Quake, S.R. (2016). Cellular Taxonomy of the Mouse Striatum as Revealed by Single-Cell RNA-Seq. Cell Rep *16*, 1126–1137.

Guo, Y., Shen, X.-N., Hou, X.-H., Ou, Y.-N., Huang, Y.-Y., Dong, Q., Tan, L., Yu, J.-T., Alzheimer's Disease Neuroimaging Initiative (2020). Genome-wide association study of white matter hyperintensity volume in elderly persons without dementia. NeuroImage: Clinical *26*, 102209.

Habib, N., Avraham-Davidi, I., Basu, A., Burks, T., Shekhar, K., Hofree, M., Choudhury, S.R., Aguet, F., Gelfand, E., Ardlie, K., et al. (2017). Massively parallel single-nucleus RNA-seq with DroNc-seq. Nat. Methods *14*, 955–958.

Habib, N., Li, Y., Heidenreich, M., Swiech, L., Avraham-Davidi, I., Trombetta, J.J., Hession, C., Zhang, F., and Regev, A. (2016). Div-Seq: Single-nucleus RNA-Seq reveals dynamics of rare adult newborn neurons. Science *353*, 925–928.

Hammond, T.R., Dufort, C., Dissing-Olesen, L., Giera, S., Young, A., Wysoker, A., Walker, A.J., Gergits, F., Segel, M., Nemesh, J., et al. (2019). Single-Cell RNA Sequencing of Microglia throughout the Mouse Lifespan and in the Injured Brain Reveals Complex Cell-State Changes. Immunity *50*, 253–271.e256.

He, L., Vanlandewijck, M., Mäe, M.A., Andrae, J., Ando, K., Del Gaudio, F., Nahar, K., Lebouvier, T., Laviña, B., Gouveia, L., et al. (2018). Single-cell RNA sequencing of mouse brain and lung vascular and vessel-associated cell types. Sci Data *5*, 180160–11.

Hodge, R.D., Miller, J.A., Novotny, M., Kalmbach, B.E., Ting, J.T., Bakken, T.E., Aevermann, B.D., Barkan, E.R., Berkowitz-Cerasano, M.L., Cobbs, C., et al. (2020). Transcriptomic evidence that von Economo neurons are regionally specialized extratelencephalic-projecting excitatory neurons. Nature Communications *11*, 1172–14.

Huang, Y., Fernandez, S.V., Goodwin, S., Russo, P.A., Russo, I.H., Sutter, T.R., and Russo, J. (2007). Epithelial to mesenchymal transition in human breast epithelial cells transformed by 17beta-estradiol. Cancer Research *67*, 11147–11157.

Jäkel, S., Agirre, E., Mendanha Falcão, A., van Bruggen, D., Lee, K.W., Knuesel, I., Malhotra, D., ffrench-Constant, C., Williams, A., and Castelo-Branco, G. (2019). Altered human oligodendrocyte heterogeneity in multiple sclerosis. Nature *566*, 543–547.

Joubert, B.R., Lange, E.M., Franceschini, N., Mwapasa, V., North, K.E., Meshnick, S.R., NIAID Center for HIV/AIDS Vaccine Immunology (2010). A whole genome association study of mother-to-child transmission of HIV in Malawi. Genome Med *2*, 17–11.

Kirby, L., Jin, J., Cardona, J.G., Smith, M.D., Martin, K.A., Wang, J., Strasburger, H., Herbst, L., Alexis, M., Karnell, J., et al. (2019). Oligodendrocyte precursor cells present antigen and are cytotoxic targets in inflammatory demyelination. Nature Communications *10*, 1–20.

Korsunsky, I., Millard, N., Fan, J., Slowikowski, K., Zhang, F., Wei, K., Baglaenko, Y., Brenner, M., Loh, P.-R., and Raychaudhuri, S. (2019). Fast, sensitive and accurate integration of single-cell data with Harmony. Nature Methods 2017 14:3 *16*, 1289–1296.

Kozareva, V. (2020). A transcriptomic atlas of the mouse cerebellum reveals regional specializations and novel cell types. bioRxiv *11*, 2020.03.04.976407.

Krienen, F.M., Goldman, M., Zhang, Q., C H Del Rosario, R., Florio, M., Machold, R., Saunders, A., Levandowski, K., Zaniewski, H., Schuman, B., et al. (2020). Innovations present in the primate interneuron repertoire. Nature *586*, 262–269.

La Manno, G. (2020). Molecular architecture of the developing mouse brain. bioRxiv *135C*, 2020.07.02.184051.

Lake, B.B., Chen, S., Sos, B.C., Fan, J., Kaeser, G.E., Yung, Y.C., Duong, T.E., Gao, D., Chun, J., Kharchenko, P.V., et al. (2018). Integrative single-cell analysis of transcriptional and epigenetic states in the human adult brain. Nat. Biotechnol. *36*, 70–80.

Lambert, S.A., Jolma, A., Campitelli, L.F., Das, P.K., Yin, Y., Albu, M., Chen, X., Taipale, J., Hughes, T.R., and Weirauch, M.T. (2018). The Human Transcription Factors. Cell *172*, 650–665.

Li, Y., Deng, G., Qi, Y., Zhang, H., Gao, L., Jiang, H., Ye, Z., Liu, B., and Chen, Q. (2020). Bioinformatic Profiling of Prognosis-Related Genes in Malignant Glioma Microenvironment. Med Sci Monit *26*, e924054.

Li, Z., Qiu, R., Qiu, X., and Tian, T. (2018). EYA4 Promotes Cell Proliferation Through Downregulation of p27Kip1 in Glioma. Cell. Physiol. Biochem. *49*, 1856–1869.

Liu, C., Ye, F.Q., Newman, J.D., Szczupak, D., Tian, X., Yen, C.C.-C., Majka, P., Glen, D., Rosa, M.G.P., Leopold, D.A., et al. (2020). A resource for the detailed 3D mapping of white matter pathways in the marmoset brain. Nat. Neurosci. *23*, 271–280.

Liu, C., Ye, F.Q., Yen, C.C.-C., Newman, J.D., Glen, D., Leopold, D.A., and Silva, A.C. (2018a). A digital 3D atlas of the marmoset brain based on multi-modal MRI. Neuroimage *169*, 106–116.

Liu, H., Luo, K., and Luo, D. (2018b). Guanosine monophosphate reductase 1 is a potential therapeutic target for Alzheimer's disease. Sci Rep *8*, 2759–10.

Luciano, N.J., Sati, P., Nair, G., Guy, J.R., Ha, S.-K., Absinta, M., Chiang, W.-Y., Leibovitch, E.C., Jacobson, S., Silva, A.C., et al. (2016). Utilizing 3D Printing Technology to Merge MRI with Histology: A Protocol for Brain Sectioning. J Vis Exp e54780–e54780.

Marisca, R., Hoche, T., Agirre, E., Hoodless, L.J., Barkey, W., Auer, F., Castelo-Branco, G., and Czopka, T. (2020). Functionally distinct subgroups of oligodendrocyte precursor cells integrate neural activity and execute myelin formation. Nat. Neurosci. *23*, 363–374.

Marques, S., Zeisel, A., Codeluppi, S., van Bruggen, D., Mendanha Falcão, A., Xiao, L., Li, H., Haring, M., Hochgerner, H., Romanov, R.A., et al. (2016). Oligodendrocyte heterogeneity in the mouse juvenile and adult central nervous system. Science *352*, 1326–1329.

Matson, K.J.E., Sathyamurthy, A., Johnson, K.R., Kelly, M.C., Kelley, M.W., and Levine, A.J. (2018). Isolation of Adult Spinal Cord Nuclei for Massively Parallel Single-nucleus RNA Sequencing. J Vis Exp.

McGinnis, C.S., Murrow, L.M., and Gartner, Z.J. (2019). DoubletFinder: Doublet Detection in Single-Cell RNA Sequencing Data Using Artificial Nearest Neighbors. Cell Syst *8*, 329–337.e4.

Morcom, L., Gobius, I., Marsh, A.P.L., Suárez, R., Bridges, C., Ye, Y., Fenlon, L.R., Zagar, Y., Douglass, A.M., Donahoo, A.-L.S., et al. (2020). The DCC receptor regulates astroglial development essential for telencephalic morphogenesis and corpus callosum formation. bioRxiv *1*, 2020.08.03.233593.

Nakano, T., Shimizu, K., Kawashima, O., Kamiyoshihara, M., Kakegawa, S., Sugano, M., Ibe, T., Nagashima, T., Kaira, K., Sunaga, N., et al. (2012). Establishment of a human lung cancer cell line with high metastatic potential to multiple organs: gene expression associated with metastatic potential in human lung cancer. Oncol. Rep. *28*, 1727–1735.

Nakatani, K., Yoshimi, N., Mori, H., Sakai, H., Shinoda, J., Andoh, T., and Sakai, N. (1998). The Significance of the Expression of Tumor Suppressor Gene DCC in Human Gliomas. J Neurooncol *40*, 237–242.

Nguyen, N.T., Vivès, R.R., Torres, M., Delauzun, V., Saesen, E., Roig-Zamboni, V., Lortat-Jacob, H., Rihet, P., and Bourne, Y. (2018). Genetic and enzymatic characterization of 3-O-sulfotransferase SNPs associated with Plasmodium falciparum parasitaemia. Glycobiology *28*, 534–541.

Ogawa, Y., Takebayashi, H., Takahashi, M., Osumi, N., Iwasaki, Y., and Ikenaka, K. (2005). Gliogenic radial glial cells show heterogeneity in the developing mouse spinal cord. Dev. Neurosci. *27*, 364–377.

Polioudakis, D., la Torre-Ubieta, de, L., Langerman, J., Elkins, A.G., Shi, X., Stein, J.L., Vuong, C.K., Nichterwitz, S., Gevorgian, M., Opland, C.K., et al. (2019). A Single-Cell Transcriptomic Atlas of Human Neocortical Development during Mid-gestation. Neuron *103*, 785–801.e788.

Raudvere, U., Kolberg, L., Kuzmin, I., Arak, T., Adler, P., Peterson, H., and Vilo, J. (2019). g:Profiler: a web server for functional enrichment analysis and conversions of gene lists (2019 update). Nucleic Acids Research *47*, W191–W198.

Reyes-Mugica, M., Rieger-Christ, K., Ohgaki, H., Ekstrand, B.C., Helie, M., Kleinman, G., Yahanda, A., Fearon, E.R., Kleihues, P., and Reale, M.A. (1997). Loss of DCC expression and glioma progression. Cancer Research *57*, 382–386.

Rodrigues, G., Hoshino, A., Kenific, C.M., Matei, I.R., Steiner, L., Freitas, D., Kim, H.S., Oxley, P.R., Scandariato, I., Casanova-Salas, I., et al. (2019). Tumour exosomal CEMIP protein promotes cancer cell colonization in brain metastasis. Nat. Cell Biol. *21*, 1403–1412.

Rosenberg, A.B., Roco, C.M., Muscat, R.A., Kuchina, A., Sample, P., Yao, Z., Graybuck, L.T., Peeler, D.J., Mukherjee, S., Chen, W., et al. (2018). Single-cell profiling of the developing mouse brain and spinal cord with split-pool barcoding. Science *360*, 176–182.

Russ, D.E., Cross, R.B.P., Li, L., Koch, S.C., Matson, K.J.E., and Levine, A.J. (2020). A Harmonized Atlas of Spinal Cord Cell Types and Their Computational Classification. bioRxiv *6*, 2020.09.03.241760.

Sathyamurthy, A., Johnson, K.R., Matson, K.J.E., Dobrott, C.I., Li, L., Ryba, A.R., Bergman, T.B., Kelly, M.C., Kelley, M.W., and Levine, A.J. (2018). Massively Parallel Single Nucleus Transcriptional Profiling Defines Spinal Cord Neurons and Their Activity during Behavior. Cell Rep *22*, 2216–2225.

Sati, P., Silva, A.C., van Gelderen, P., Gaitán, M.I., Wohler, J.E., Jacobson, S., Duyn, J.H., and Reich, D.S. (2012). In vivo quantification of $T_2$ anisotropy in white matter fibers in marmoset monkeys. Neuroimage *59*, 979–985.

Sekar, S., McDonald, J., Cuyugan, L., Aldrich, J., Kurdoglu, A., Adkins, J., Serrano, G., Beach, T.G., Craig, D.W., Valla, J., et al. (2015). Alzheimer's disease is associated with altered expression of genes involved in immune response and mitochondrial processes in astrocytes. Neurobiol. Aging *36*, 583–591.

Shimogori, T., Abe, A., Go, Y., Hashikawa, T., Kishi, N., Kikuchi, S.S., Kita, Y., Niimi, K., Nishibe, H., Okuno, M., et al. (2018). Digital gene atlas of neonate common marmoset brain. Neurosci. Res. *128*, 1–13.

Skene, N.G., and Grant, S.G.N. (2016). Identification of Vulnerable Cell Types in Major Brain Disorders Using Single Cell Transcriptomes and Expression Weighted Cell Type Enrichment. Front Neurosci *10*, 16.

Stuart, T., Butler, A., Hoffman, P., Hafemeister, C., Papalexi, E., Mauck, W.M., Hao, Y., Stoeckius, M., Smibert, P., and Satija, R. (2019). Comprehensive Integration of Single-Cell Data. Cell *177*, 1888–1902.e21.

Tasic, B., Menon, V., Nguyen, T.N., Kim, T.K., Jarsky, T., Yao, Z., Levi, B., Gray, L.T., Sorensen, S.A., Dolbeare, T., et al. (2016). Adult mouse cortical cell taxonomy revealed by single cell transcriptomics. Nat. Neurosci. *19*, 335–346.

Tirosh, I., Izar, B., Prakadan, S.M., Wadsworth, M.H., Treacy, D., Trombetta, J.J., Rotem, A., Rodman, C., Lian, C., Murphy, G., et al. (2016). Dissecting the multicellular ecosystem of metastatic melanoma by single-cell RNA-seq. Science *352*, 189–196.

van Wageningen, T.A., Gerrits, E., Geleijnse, A., Brouwer, N., Geurts, J.J.G., Eggen, B.J.L., Boddeke, H.W.G.M., and van Dam, A.-M. (2020). Distinct gene expression profiles in leukocortical demyelinated white and grey matter areas of Multiple Sclerosis patients. bioRxiv *9*, 2020.06.03.131300.

Vanlandewijck, M., He, L., Mäe, M.A., Andrae, J., Ando, K., Del Gaudio, F., Nahar, K., Lebouvier, T., Laviña, B., Gouveia, L., et al. (2018). A molecular atlas of cell types and zonation in the brain vasculature. Nature *554*, 475–480.

Wang, W., Liu, J.-Y., Wang, H.-F., Yang, M.-Y., Liu, Q.-Y., and Ding, M.-X. (2016). The complete mitochondrial genome of white-tufted-ear marmoset, Callithrix jacchus (Primates: Callitrichinae). Mitochondrial DNA a DNA Mapp Seq Anal *27*, 1920–1921.

Xia, L., Huang, Q., Nie, D., Shi, J., Gong, M., Wu, B., Gong, P., Zhao, L., Zuo, H., Ju, S., et al. (2013). PAX3 is overexpressed in human glioblastomas and critically regulates the tumorigenicity of glioma cells. Brain Res *1521*, 68–78.

Xiao, L., Ohayon, D., McKenzie, I.A., Sinclair-Wilson, A., Wright, J.L., Fudge, A.D., Emery, B., Li, H., and Richardson, W.D. (2016). Rapid production of new oligodendrocytes is required in the earliest stages of motor-skill learning. Nat. Neurosci. *19*, 1210–1217.

Yap, J.K.Y., Pickard, B.S., Chan, E.W.L., and Gan, S.Y. (2019). The Role of Neuronal NLRP1 Inflammasome in Alzheimer's Disease: Bringing Neurons into the Neuroinflammation Game. Mol Neurobiol *56*, 7741–7753.

Young, M.D., and Behjati, S. (2020). SoupX removes ambient RNA contamination from droplet based single-cell RNA sequencing data. bioRxiv *12*, 303727.

Yuste, R., Hawrylycz, M., Aalling, N., Aguilar-Valles, A., Arendt, D., Arnedillo, R.A., Ascoli, G.A., Bielza, C., Bokharaie, V., Bergmann, T.B., et al. (2020). A community-based transcriptomics classification and nomenclature of neocortical cell types. Nat. Neurosci. *18*, 361–13.

Zappia, L., and Oshlack, A. (2018). Clustering trees: a visualization for evaluating clusterings at multiple resolutions. Gigascience *7*.

Zeisel, A., Hochgerner, H., Lonnerberg, P., Johnsson, A., Memic, F., van der Zwan, J., Haring, M., Braun, E., Borm, L.E., La Manno, G., et al. (2018). Molecular Architecture of the Mouse Nervous System. Cell *174*, 999–.

Zeisel, A., Muñoz-Manchado, A.B., Codeluppi, S., Lonnerberg, P., La Manno, G., Juréus, A., Marques, S., Munguba, H., He, L., Betsholtz, C., et al. (2015). Brain structure. Cell types in the mouse cortex and hippocampus revealed by single-cell RNA-seq. Science *347*, 1138–1142.

Zhang, Y., Chen, K., Sloan, S.A., Bennett, M.L., Scholze, A.R., O'Keeffe, S., Phatnani, H.P., Guarnieri, P., Caneda, C., Ruderisch, N., et al. (2014). An RNA-sequencing transcriptome and splicing database of glia, neurons, and vascular cells of the cerebral cortex. J. Neurosci. *34*, 11929–11947.

Zhang, Y., Sloan, S.A., Clarke, L.E., Caneda, C., Plaza, C.A., Blumenthal, P.D., Vogel, H., Steinberg, G.K., Edwards, M.S.B., Li, G., et al. (2016). Purification and Characterization of Progenitor and Mature Human Astrocytes Reveals Transcriptional and Functional Differences with Mouse. Neuron *89*, 37–53.

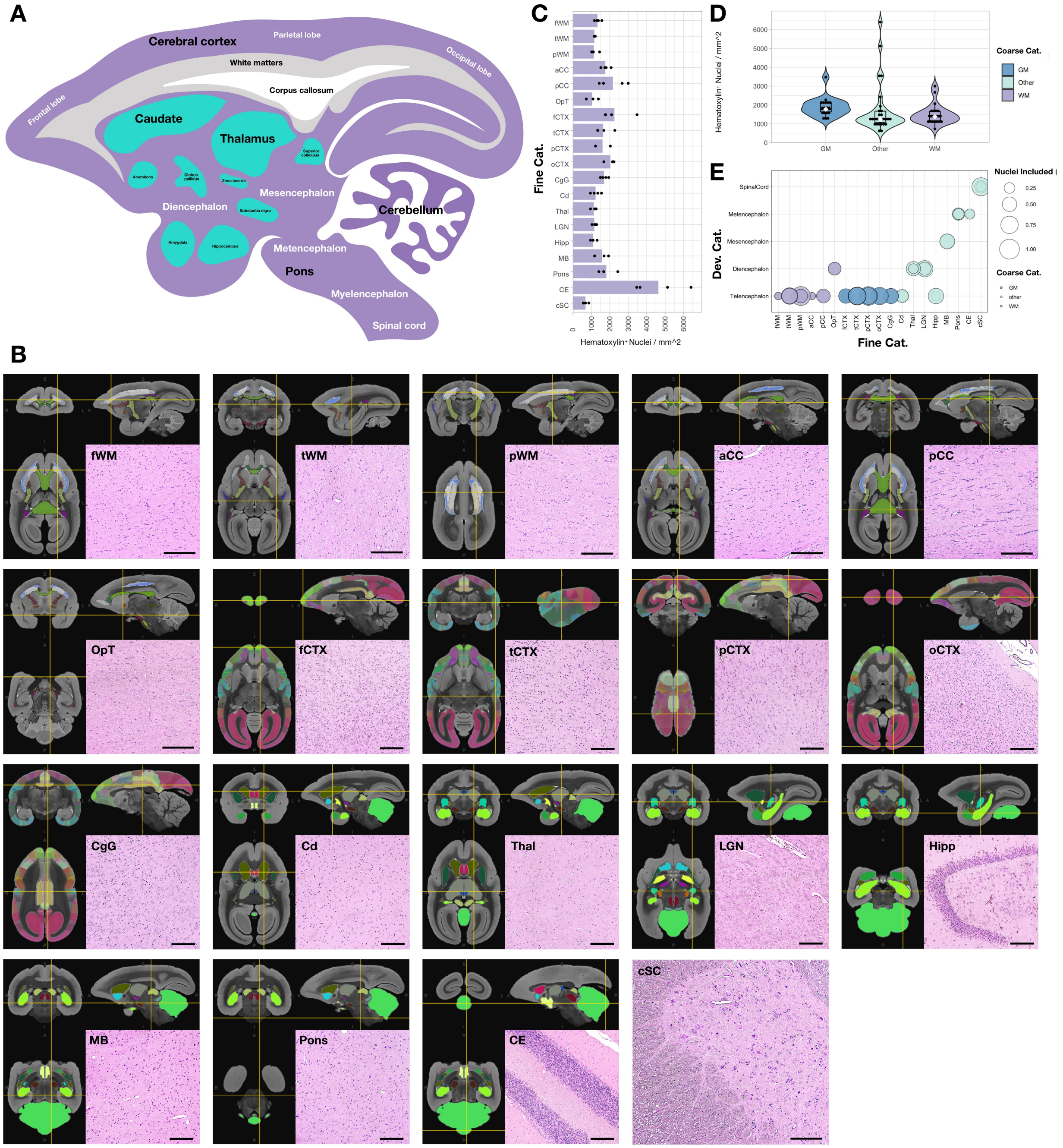


**Figure S1. Related to Figure 1. Location reference for image guided tissue sampling and quantification of nuclei density per area.**

(A) Sagittal schematic illustration of an adult marmoset brain.

(B) Hematoxylin and eosin staining of 5-μm thick tissue sections from areas as indicated on the marmoset brain mapping MRI atlas. Sections were selected by matching with regions that were sampled for snRNA-seq. **fWM**, frontal white matter; **tWM**, temporal white matter; **pWM**, parietal white matter; **aCC**, anterior corpus callosum; **pCC**, posterior corpus callosum; **OpT**, optic tract; **fCTX**, frontal cortex; **tCTX**, temporal cortex; **pCTX**, parietal cortex; **oCTX**, occipital cortex; **CgG**, cingulate gyrus; **Cd**, caudate; **Thal**, thalamus; **LGN**, lateral geniculate nucleus; **Hipp**, hippocampus; **MB**, midbrain; **Pons**, pons; **CE**, cerebellum; **cSC**, cervical spinal cord. Scale bar, 200 μm.

(C) Bar plot of hematoxylin$^+$ nuclei density from each region (category IL06_tissue.3, fine annotation) that were quantified from 2–4 animals (CJT11, CJV13, CJB10, and CJD12).

(D) Violin plot of hematoxylin$^+$ nuclei density in each tissue type (coarse category) "WM" includes fWM, tWM, pWM, aCC, pCC, and OpT; "GM" includes fCTX, tCTX, pCTX, oCTX, and CgG; "other" includes Cd, Thal, LGN, Hipp, MB, Pons, CE, and cSC.

(E) Dot plot of estimated nuclei recovery as a percentage of each snRNA-seq sample from each tissue type (developmental category). Each circle represents the percentage from one sample. "Telencephalon" includes fWM, tWM, pWM, aCC, pCC, fCTX, tCTX, pCTX, oCTX, CgG, Cd, Hipp; "Diencephalon" includes OpT, Thal, LGN; "Mesencephalon" includes MB; "Metencephalon" includes Pons and CE; "Spinal cord" includes cSC.

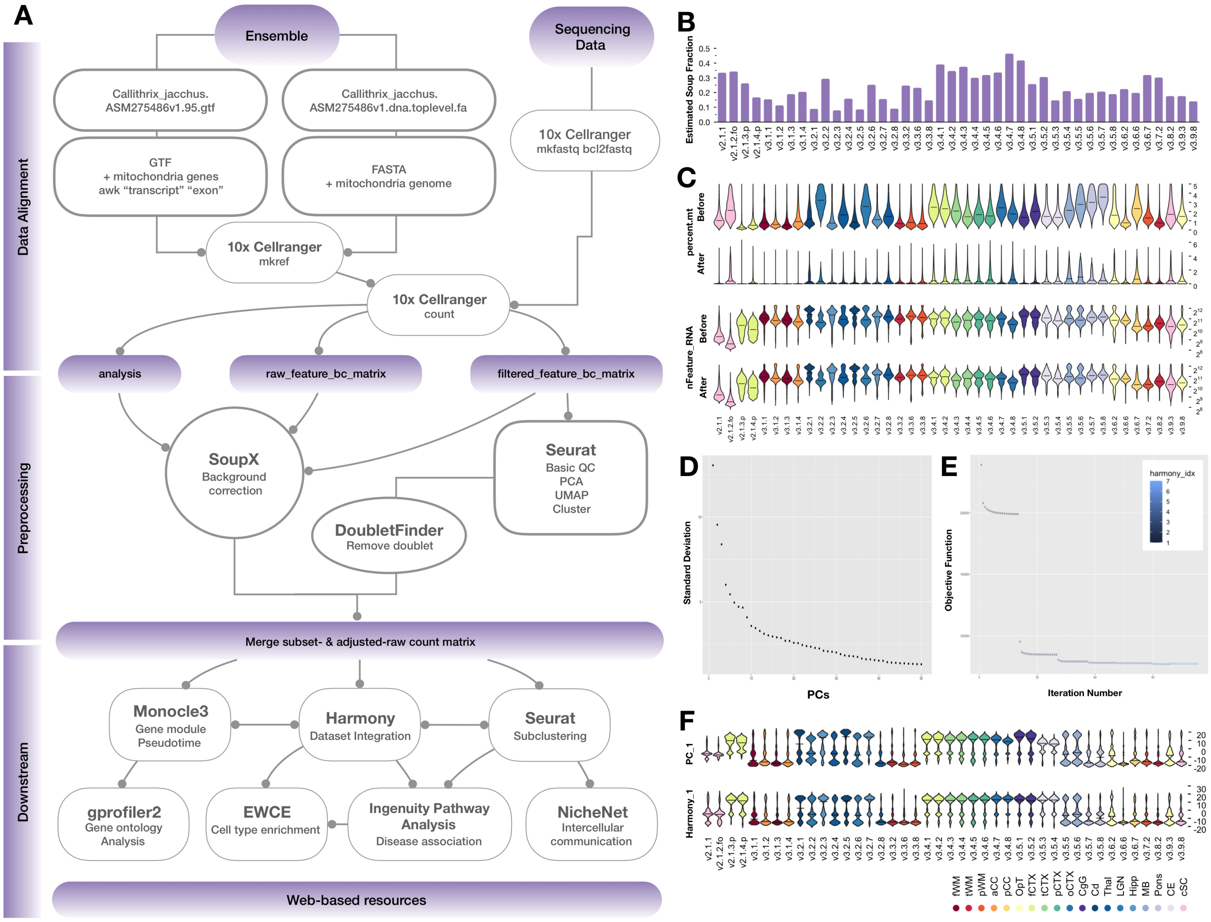


**Figure S2. Related to Figure 1. Workflow of data processing and basic feature assessment of individual samples before data integration and Level 1 partition.**

(A) A genome assembly (ASM275486v1) and annotation were acquired from the Ensembl release-95, and the marmoset mitochondrial sequences (NC_025586.1, GenBank) were manually added to the file. An alignment reference package that was tailored for snRNA-seq was generated by 10x Cellranger mkref function. Raw count matrices were generated by 10x Cellranger count function. For individual samples, the output of 10x Cellranger was then processed in parallel with SoupX to remove ambient RNA, and with Seurat v3 followed by DoubletFinder to remove low quality nuclei and putative doublets. The adjusted matrices of each sample from SoupX output were subset by nuclei names that passed the Seurat v3 and DoubletFinder quality control step to become matrices used for the Level 1 partition. Adjusted matrices from individual samples were combined and integrated using Seurat v3 and Harmony. Monocle, gprofiler2, Expression Weighted Celltype Enrichment (EWCE), Ingenuity Pathway Analysis (IPA) software, and NicheNet algorithms were used for downstream analysis.

(B) Bar plot showing the fraction of ambient RNA estimated by SoupX for each sample.

(C) Violin plots showing the percentage of mitochondrial derived RNA reads (percent.mt), and the number of RNA species (nFeature_RNA), before and after SoupX correction. Median is annotated (–).

(D) Elbow plot showing individual standard deviation of the top 50 principal components (PC) in the Level 1 analysis.

(E) Convergence plot showing that Harmony converged after 7 iterations.

(F) Violin plots showing the nuclei density along the PC_1 and Harmony_1 embedding axes. The distributions of nuclei are aligned between samples acquired from the similar regions after Harmony. The color of each violin denotes the tissue of origin, as shown in the index at the bottom. Median is annotated (–).

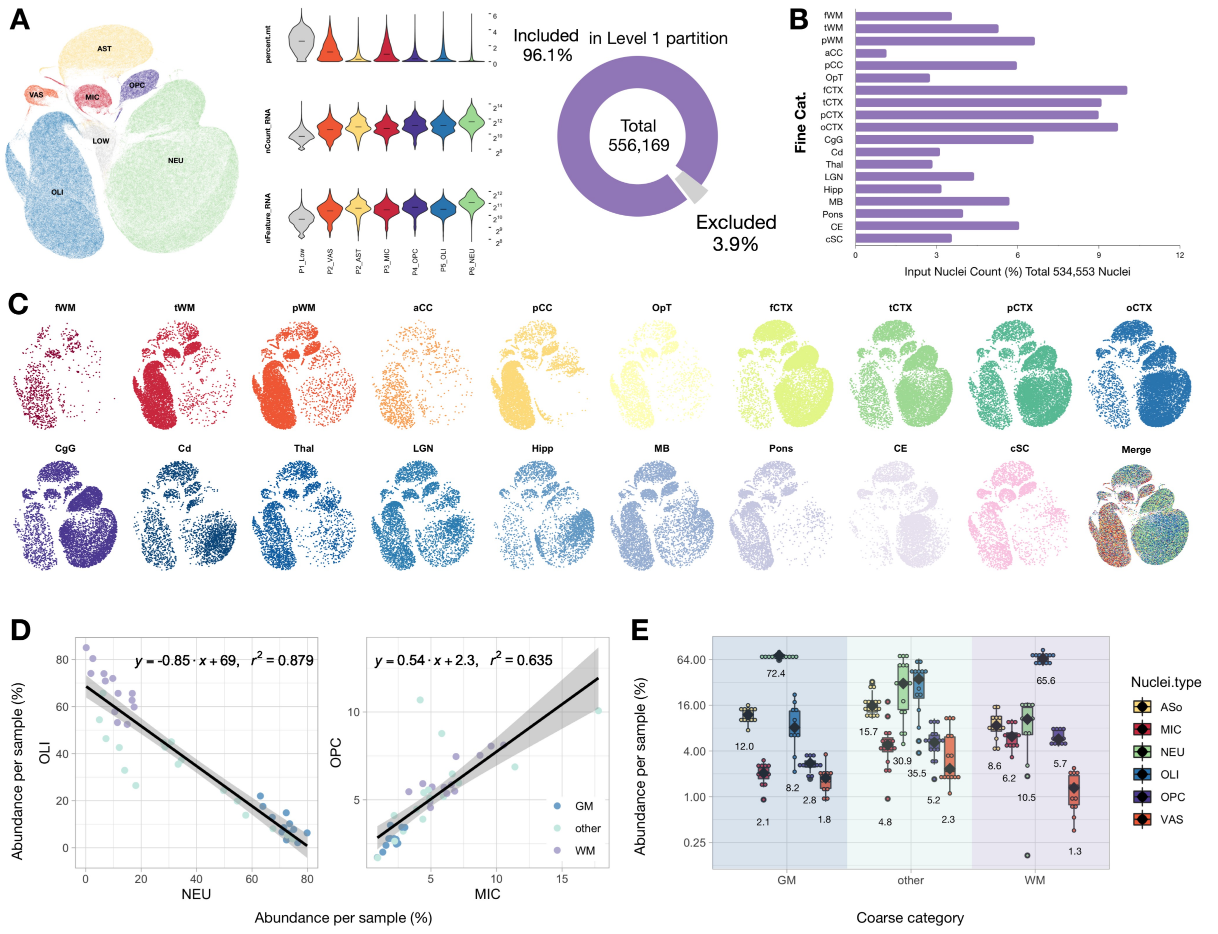


**Figure S3. Related to Figure 1. Assessment of nuclei features in Level 1 partition by tissue location.**

(A) Left, global UMAP scatter plot visualization of 556,169 nuclei after SoupX and DoubletFinder preprocessing steps. The UMAP space and nearest neighbor analyses were calculated from the top 5 Harmony embeddings. A "low quality" cluster was annotated and removed from further Level 1 analysis. The "low quality" cluster was informed by clustering multiple times with different parameters that met exclusion criteria, including high percentage of reads mapped to mitochondria genome, low RNA counts and features, and expressed genes that mapped to multiple canonical markers of different cell types. Middle, violin plots showing the percentage of mitochondrial derived RNA reads (percent.mt), the number of RNA detected (nCount_RNA), and the number of RNA species (nFeature_RNA) in each partition. Median is annotated (–). Right, donut plot showing the percentage of nuclei annotated as "low quality" and excluded from further analysis.

(B) Bar plot showing the nuclei input percentage of each tissue type in level 1 partition, where 2–3 samples were pooled from 2 animals per tissue type.

(C) Global UMAP scatter plot visualization of 100K nuclei split by tissue types. The same object was used to make plots showed in Figure 1E. As expected, white matter contains large numbers of oligodendrocytes (OLI) and gray matter large numbers of neurons (NEU).

(D) Linear regression on the relative abundance of Level 1 annotated nuclei, showing strong to moderate correlations between OLI and NEU (left) and oligodendrocyte progenitor cells (OPC) and microglia (MIC) (right). Percentage of nuclei type per sample was calculated by normalizing to their initial nuclei input. Individual samples were colored by tissue type in coarse category (See Figure 1D legend for the full list).

(E) Box plot showing the abundance of Level 1 clusters as a function of tissue type. Median is annotated (◆) and listed.

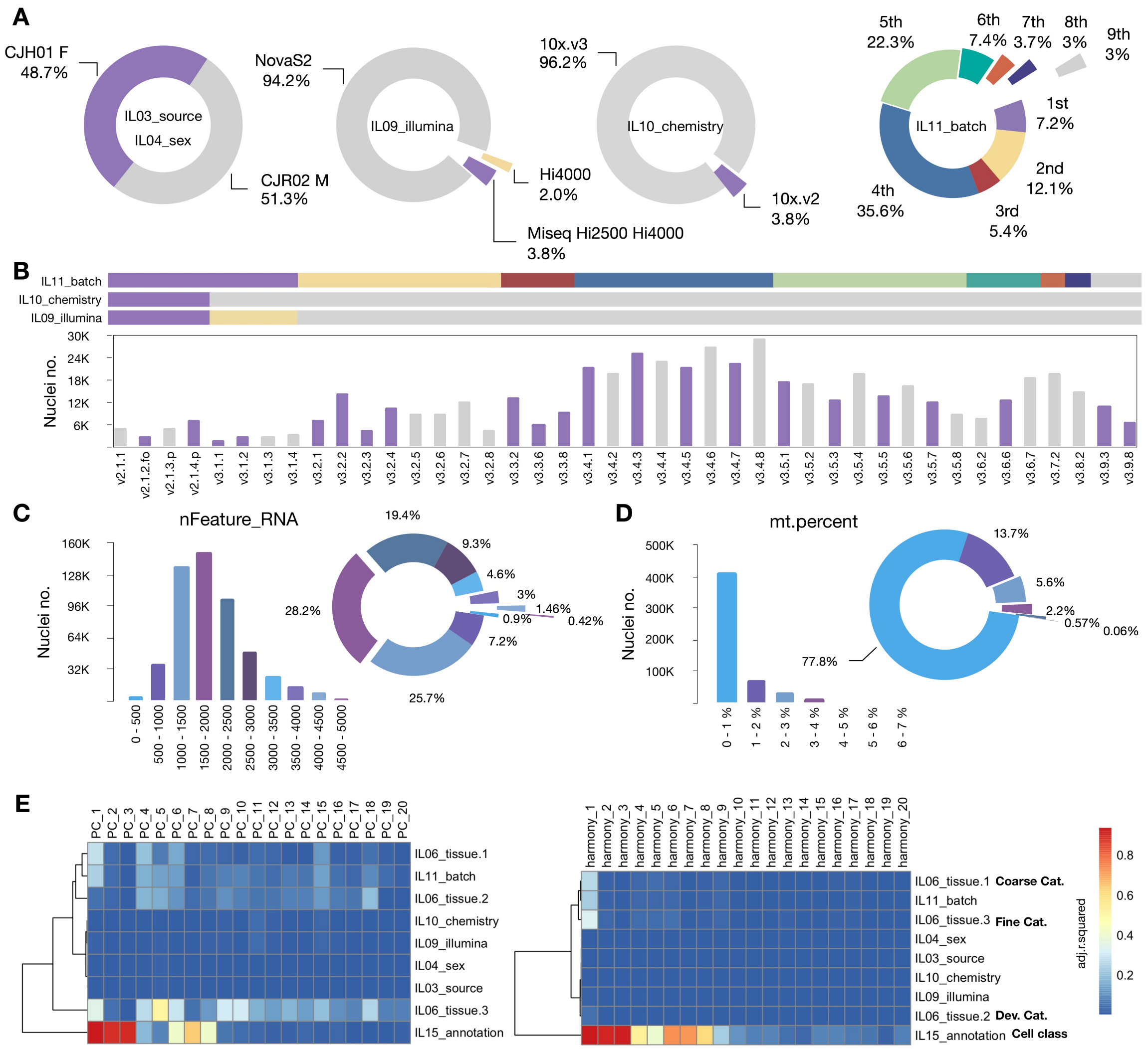


**Figure S4. Related to Figure 1. Quantification of nuclei number over each covariate and correlation of the covariate with the top 20 PC and Harmony embeddings.**

(A) Donut plots showing the proportional nuclei input from each animal and sex, sequencer, 10x chemistry version, and library preparation batch.

(B) Bar plot showing the input of nuclei number from each sample included in the Level 1 analysis. The color bars correspond to the labels showing in Figure S4A.

(C) Bar and donut plots showing the distribution of nuclei over the number of RNA features detected per nucleus. Nuclei were grouped into 10 bins with step size of 500 features (left). Proportional nuclei from each bin in percentage (right).

(D) Bar and donut plots showing the distribution of nuclei over the percentage of reads that were mapped to mitochondria genome detected per nucleus. Nuclei were grouped into 7 bins with step size of 1% (left). Percentage of nuclei in each bin (right).

(E) Heatmap showing the linear correlations (adjusted $r^2$) between the top 20 principal component (PC, left) and Harmony (right) embeddings and each sample category. Level 1 cell partition results (MIC, OPC, OLI, AST, VAS, NEU) were labeled as IL15_annotation category.

See also Table S1.

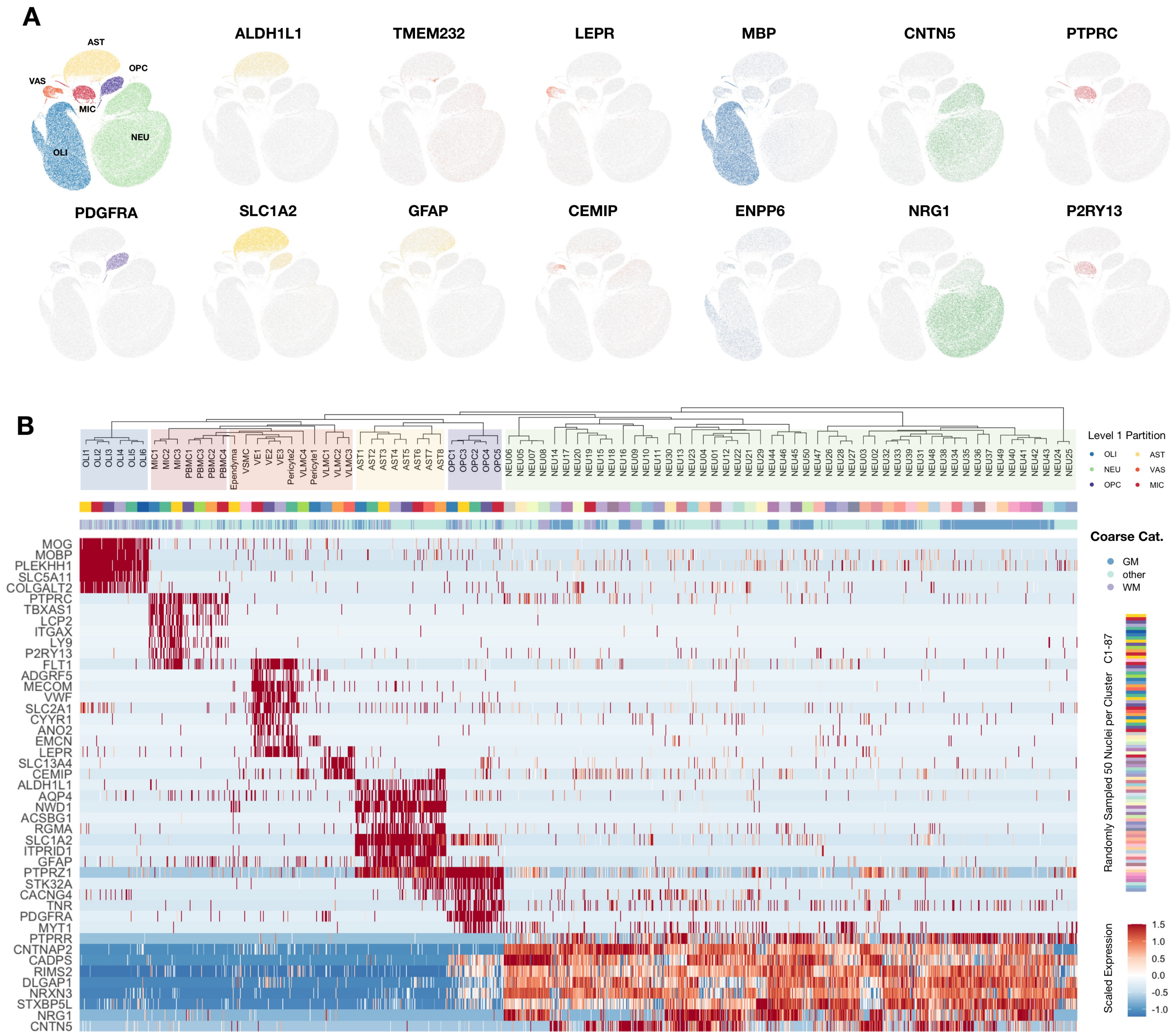


**Figure S5. Related to Figure 1. Single-nucleus transcriptional profiling of adult marmoset brain and spinal cord identifies 6 cell classes and 87 subclusters.**

(A) Cells were initially divided into 6 major classes (Level 1 clustering), comprising neurons (NEU), oligodendrocytes (OLI), oligodendrocyte progenitor cells (OPC), microglia/macrophages (MIC), astrocytes (AST), and vascular/meningeal cells (VAS). UMAP scatter plots show distinct clusters for these 6 groups, exemplified by expression of canonical marker genes. To avoid overcrowding, the scatter plots presented here represent a random sample of 100K nuclei.

(B) Each of the Level 1 classes was further subclustered (Level 2). Shown is a heatmap and dendrogram of cell types annotated from the Level 2 subclusters, comprising 50 NEU, 6 OLI, 5 OPC, 7 MIC, 8 AST, and 11 VAS subclusters. The heatmap presented here is derived from a pool of 4350 nuclei (50 per cluster, randomly selected; “C50 object”). Each column represents the expression pattern of one nucleus. The first color bar is a color code corresponding to the cluster labels at the endpoint of the dendrogram. The second color bar represents the tissue origin of the nuclei; the 22 sampling sites were grouped into 19 tissue types and further grouped in coarse category: "**WM**" (cerebral white matter), "**GM**" (cortical gray matter), and "**other**." See Figure S1D caption for the full list. The top 50 differentially enriched genes in each Level 1 cell class were used to calculate Euclidean distances and generate the dendrogram. To aid cluster tracking, the branches of the dendrogram were reordered and colored to show cluster relatedness while retaining the structure of the tree.

**A** **Cross-cluster comparison: averaged gene expression**

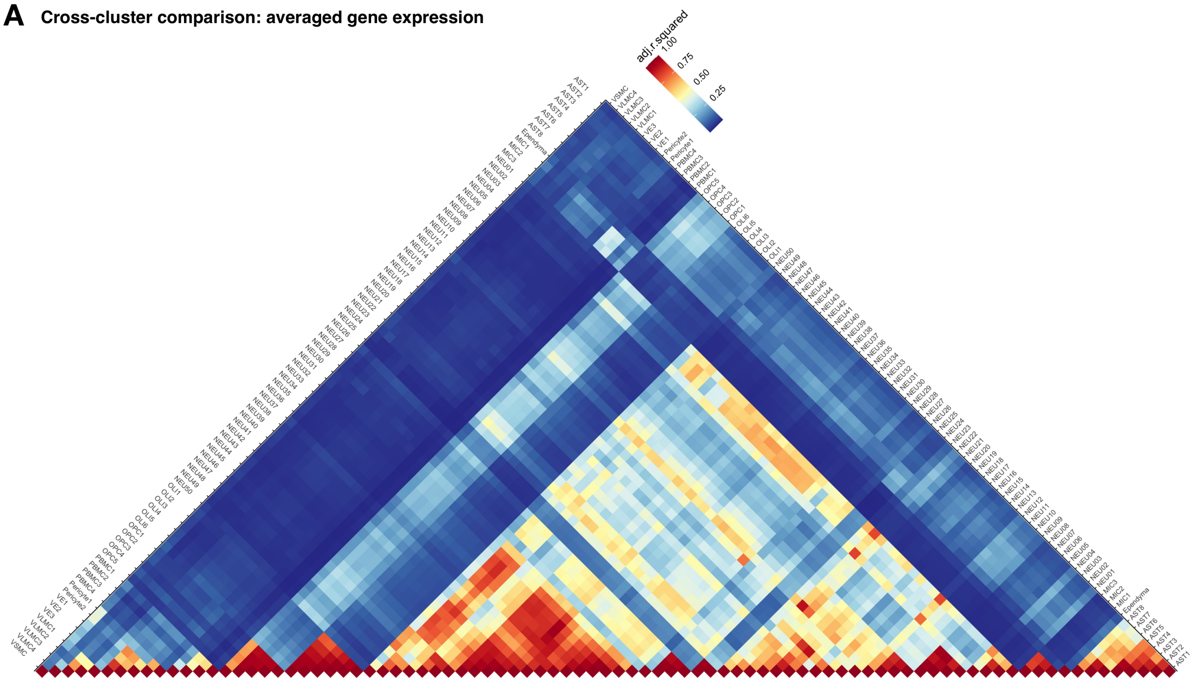

**B** **Cross-cluster comparison: differentially expressed gene number**

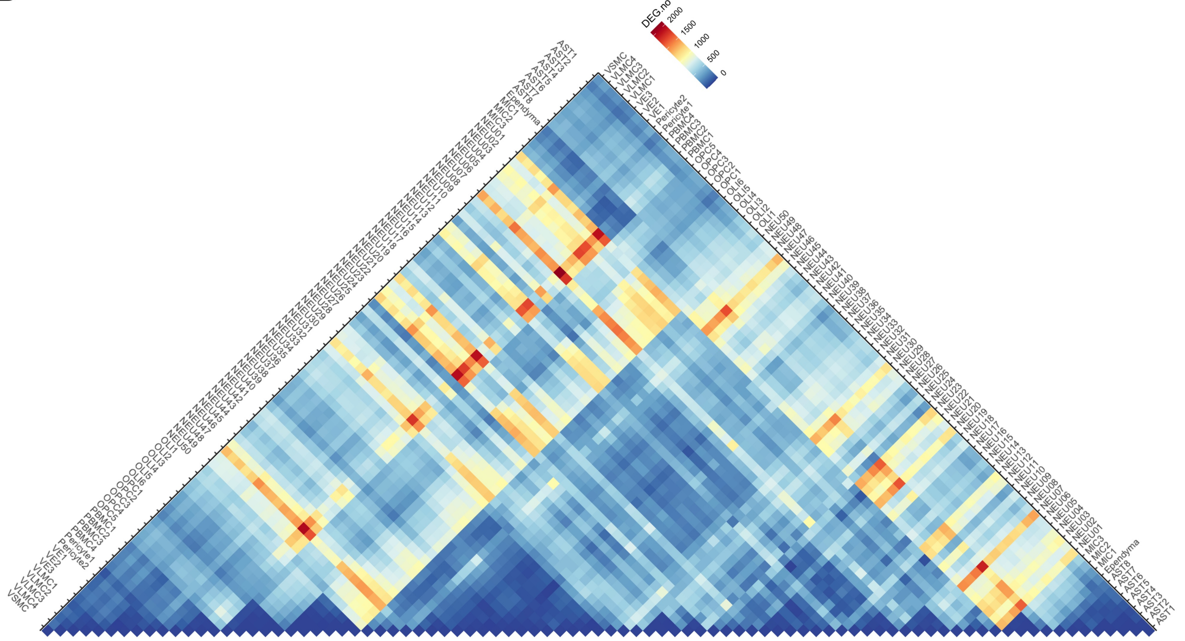

**Figure S6. Related to Figure 1. Assessment of the transcriptomic similarity across each cluster pair.**

(A) Heatmap showing the paired linear correlation (adjusted $r^2$) between 87 subclusters (C50). The expression levels of all genes within each subcluster were normalized and averaged before the comparison.

(B) Heatmap showing the number of genes differentially expressed between nuclei in each pair of the 87 subclusters. Both significantly increased and significantly decreased differentially expressed genes are included. The differentially expressed genes were further filtered by their log fold-change (ln(fold-change) > 0.25) and detection frequency (detected in at least 10% of the nuclei).

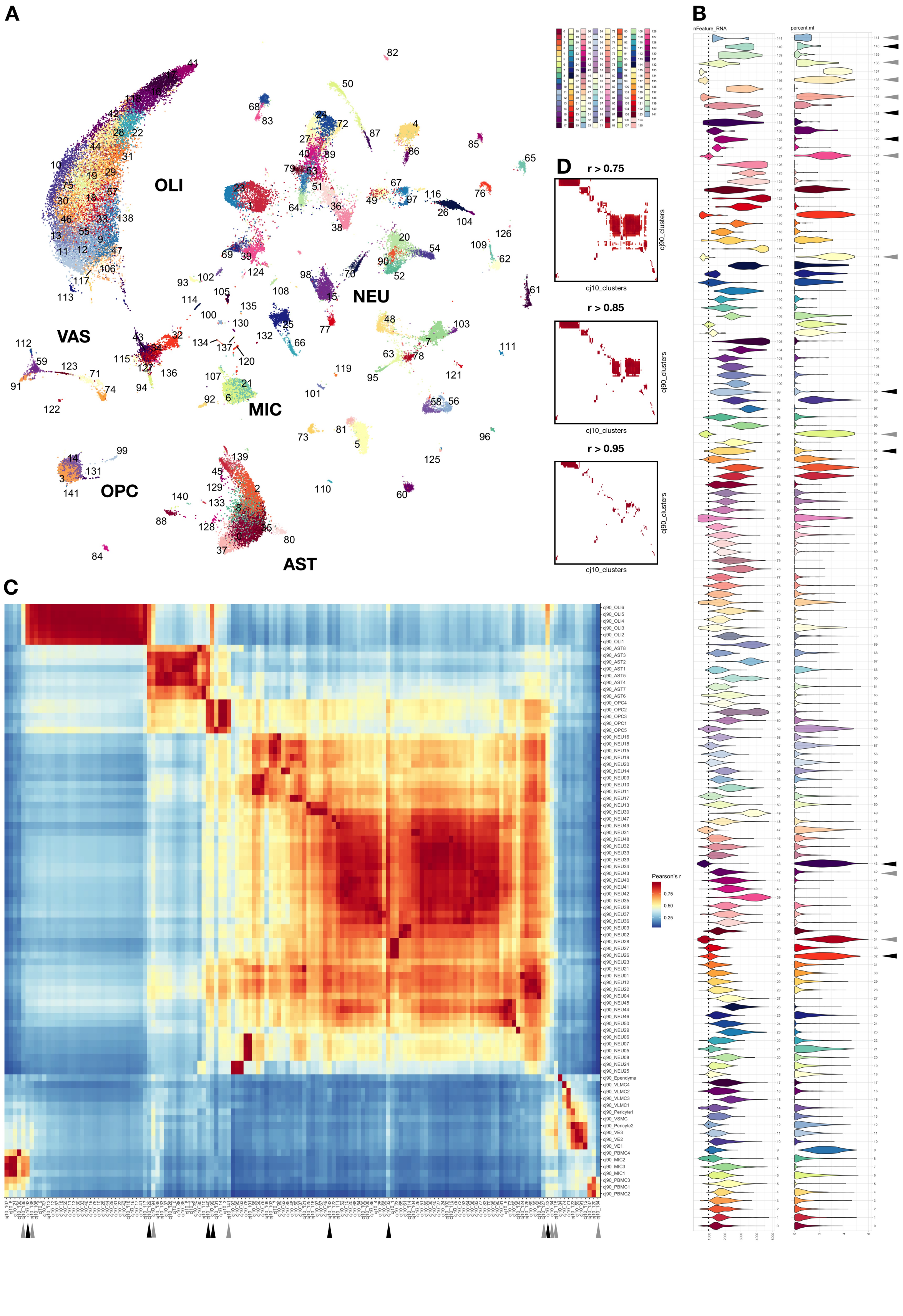
A
OLI
NEU
VAS
MIC
OPC
AST
B
nFeature_RNA
percent.mt
C
Pearson's r
D
r > 0.75
r > 0.85
r > 0.95
cj90_clusters
cj10_clusters

**Figure S7. Related to Figure 1. Unbiased classification of 10% a priori set-aside data yields clusters that recapitulate knowledge-based nuclei classes and subclusters.**

(A) UMAP scatter-plot visualization of 10% of nuclei (61,803 nuclei) that were randomly set aside from our dataset after initial quality control but prior to further analysis. Analysis parameters were intentionally set to yield a large number of clusters. The top 5000 variable genes were used to calculate 100 principal components and harmonized over IL01_uniqueID labels, then UMAP space and nearest neighbor analyses were calculated on all 100 Harmony embeddings, yielding 142 unbiased clusters.

(B) Violin plot showing the percentage of RNA reads that mapped to mitochondria genome (percent.mt) and the number of RNA species (nFeature_RNA) in each cluster. Median is annotated (–).

(C) Heatmap showing the Pearson's correlation (r) between the 87 manually annotated subclusters (C50 object, vertical axis) and 142 unbiased clusters (cj10_0 – cj10_141) from the 10% set-aside data (horizontal axis). The expression levels of all genes within each subcluster were normalized and averaged before comparison. Hot spots in the heatmap show correspondence between unbiased and knowledge-derived clusters for all cell classes. Artifact clusters, with "low quality" reads (e.g., low nFeature_RNA or high percent.mt, gray arrowheads) or nuclei doublets (high correlation with multiple cell classes, black arrowheads) are easily spotted on the heatmap.

(D) Filtered heatmaps showing the distribution of the paired correlation of Pearson's r over 0.75, 0.85, and 0.95 thresholds. At 0.75 threshold, all 87 annotated clusters correspond to one or several clusters from the set-aside data. At 0.95 cutoff, subclusters of neurons can be distinguished.

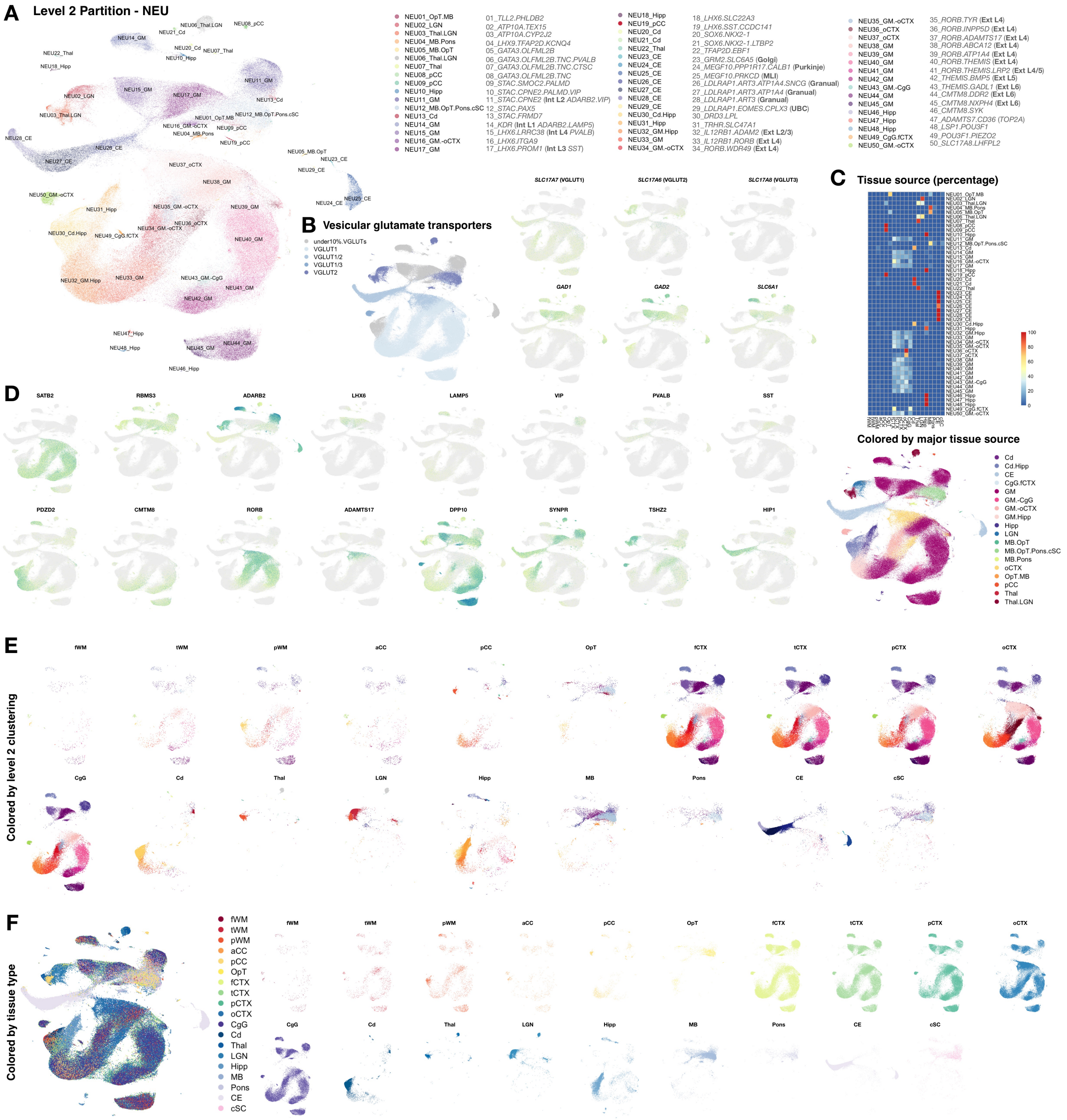


**Figure S8. Related to Figure 1. Although neuron subclusters in different cerebral cortices are mostly shared, subclusters unique to posterior corpus callosum, occipital cortex, lateral geniculate nucleus, thalamus, caudate, hippocampus, and cerebellum exist.**

(A) UMAP scatter-plot visualization of NEU nuclei colored by subcluster. Enriched markers selected from dot plot (Figure S9C) and reference indexed cell types are annotated. **L1 – L6**, layer 1 – layer 6; **Int**, inhibitory neurons; **Ext**, excitatory neurons; **MLI**, molecular layer interneurons; **UBC**, unipolar brush cell.

(B) Left, UMAP scatter-plot visualization of NEU nuclei colored by their primary VGLUT gene expression. Right, UMAP scatter-plot visualization of NEU nuclei colored by the expression of glutamatergic (*SLC17A6*, *SLC17A7*, *SLC17A8*) and GABAergic (*GAD1*, *GAD2*, *SLC6A1*) genes. See Figure S9E for detail.

(C) Top, heatmap showing the percentage of nuclei in each NEU subcluster drawn from different sampling sites; 100% per column (subcluster). Bottom, UMAP scatter-plot visualization of NEU nuclei colored by the similarity of tissue type based on the heatmap, regrouped by combining similar tissue types.

(D) UMAP scatter-plot visualization of NEU nuclei colored by the expression of genes specific to small subsets of neuron subclusters.

(E) UMAP scatter-plot visualization of NEU nuclei colored by subcluster and split by sampling site.

(F) UMAP scatter-plot visualization of neurons (NEU) colored and split by sampling site.

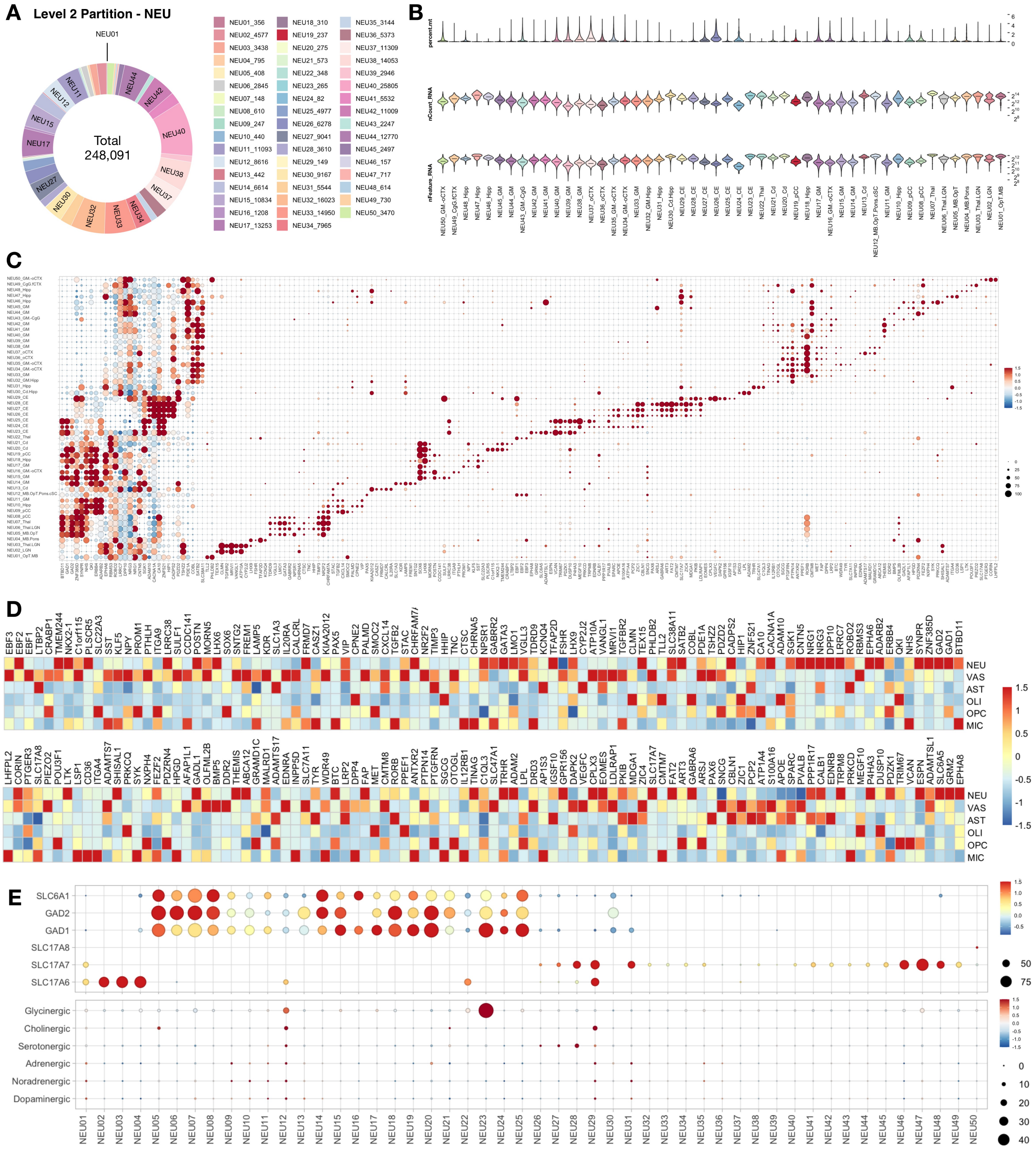


**Figure S9. Related to Figure 1. Nuclei features in neurons.**

(A) Donut plot showing the number of NEU nuclei that passed Level 2 quality control, annotated by subcluster.

(B) Violin plot showing the percentage of RNA reads that mapped to mitochondria genome (percent.mt), the number of RNA detected (nCount_RNA), and the number of RNA species (nFeature_RNA) in each cluster. Median is annotated (–).

(C) Dot plot showing mean-centered and z-score-scaled marker gene expression for NEU subclusters. Dot size indicates the percentage of nuclei in the subcluster in which each gene was detected. Scaling is relative to expression across all NEU nuclei in which a given gene was detected.

(D) Heatmap showing mean-centered and z-score-scaled NEU marker gene expression for all Level 1 classes (C50).

(E) Dot plots showing mean-centered and z-score-scaled expression of marker genes related to neurotransmission across NEU subclusters. Top, filtered dot plot; for clarity, the glutamatergic (*SLC17A6*, *SLC17A7*, *SLC17A8*) and GABAergic (*GAD1*, *GAD2*, *SLC6A1*) genes expression values were dropped if detected in <10% of nuclei in a given cluster (related to Figure S9B). Bottom, unfiltered dot plot: genes related to each neurotransmitter type were aggregated. Genes that are dopaminergic (*TH*, *DDC*, *SLC6A3*), noradrenergic (*TH*, *DCC*, *DBH*, *SLC6A2*), adrenergic (*TH*, *DDC*, *DBH*, *PNMT*), serotonergic (*TPH2*, *DDC*, *SLC6A4*), cholinergic (*CHAT*, *SLC5A7*), and glycinergic (*SLC6A9*, *SLC6A5*) were used to make this plot.

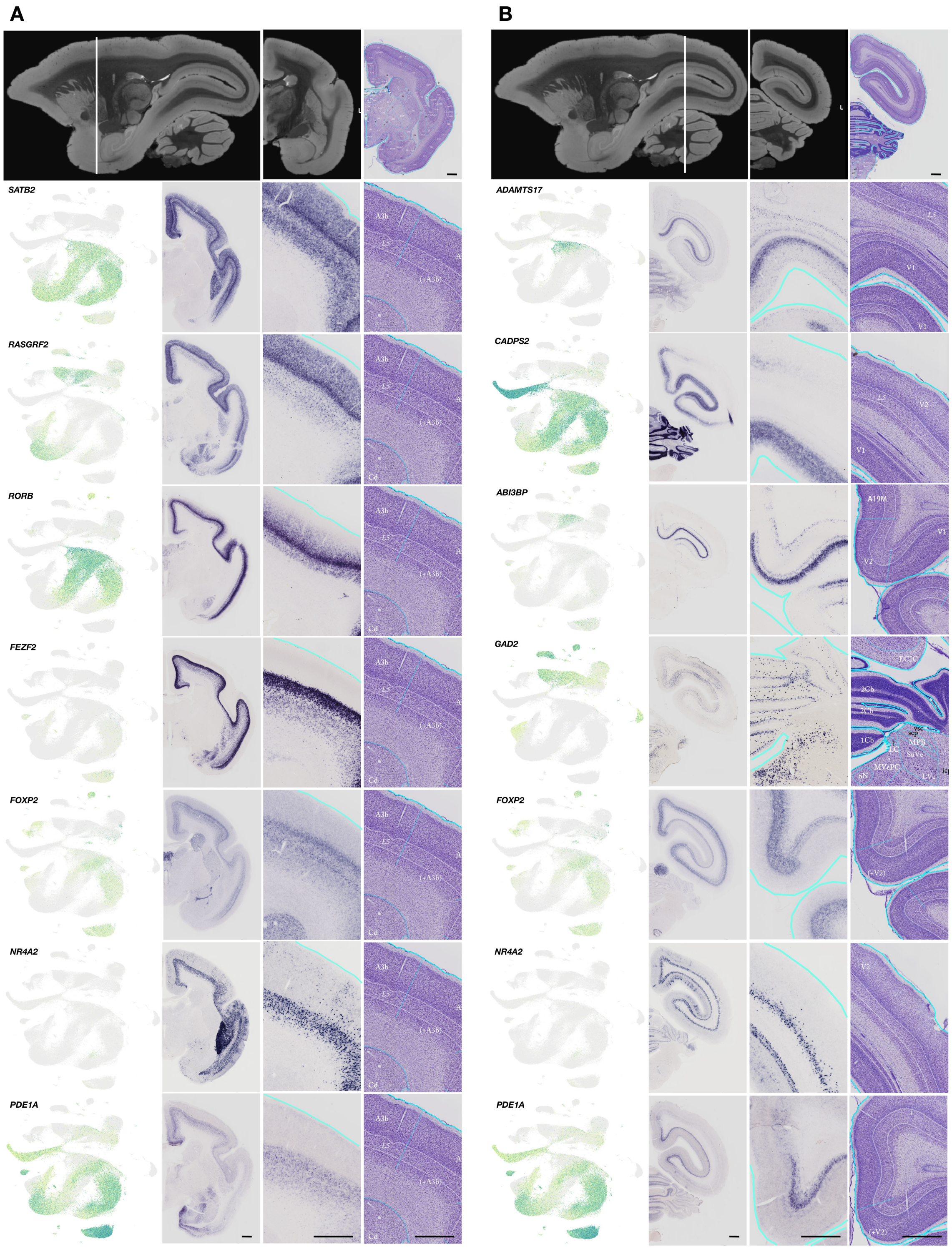


**Figure S10. Related to Figure 1. Cortical excitatory neurons are arranged into a continuous UMAP projection corresponding to lamination.**

(A – B) Approximate cross-sectional plane (white line) indicated on a sagittal view of an adult marmoset postmortem T2*-weighted MRI (row 1, left). Coronal view of the same adult marmoset T2*-weighted MRI (row 1 middle). Coronal histological section of a P0 marmoset stained with Nissl's method (right) and anatomically matched to the adult coronal T2*-weighted MRI slice (row 1, right). Rows 2–7: UMAP scatter-plot visualization of neurons colored by selected marker gene expression (column 1). Selected in situ hybridization (ISH) of genes enriched in all or a subset of cortical layers (column 2). Enlarged area of the ISH image was anatomically matched to a Nissl-stained section for regional reference (columns 3 and 4). Scale bar, 1mm. MRI images were acquired from the Marmoset Brain Mapping database. Histological Nissl stain and ISH images were acquired from the Marmoset Gene Atlas.

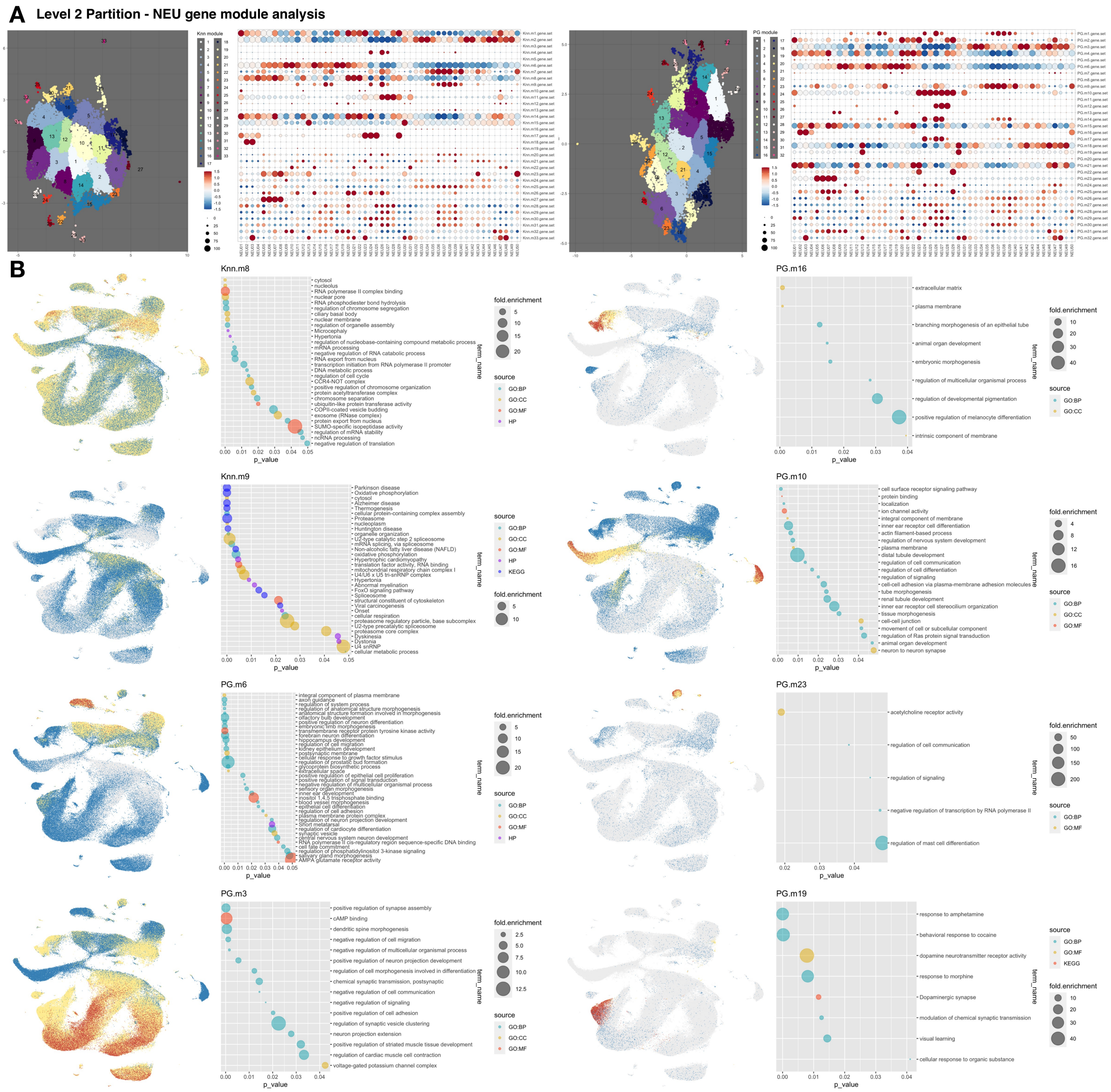


**Figure S11. Related to Figure 1. Gene module analysis in neurons.**

(A) UMAP scatter plot visualization of genes grouped by expression similarity across cells. Genes that passed Moran's *I* statistic spatial test (< 5% FDR) over *k*-nearest neighbor graph (Knn, k=25, left), or over trajectory-learned principal graph (PG, right), by Monocle3 graph_test function. Genes are grouped and colored by modules identified in each graph test by find_gene_modules function with resolution of 0.001. The list of genes in each module were aggregated through Seurat v3 AddModuleScore function. Dot plot showing the averaged and scaled expression of each module gene set across NEU subclusters.

(B) UMAP scatter-plot visualization of nuclei colored by averaged expression of selected modules enriched in a subset of neurons (see Table S2 for full list). Dot plot showing the enriched GO terms from the list of genes in each module.

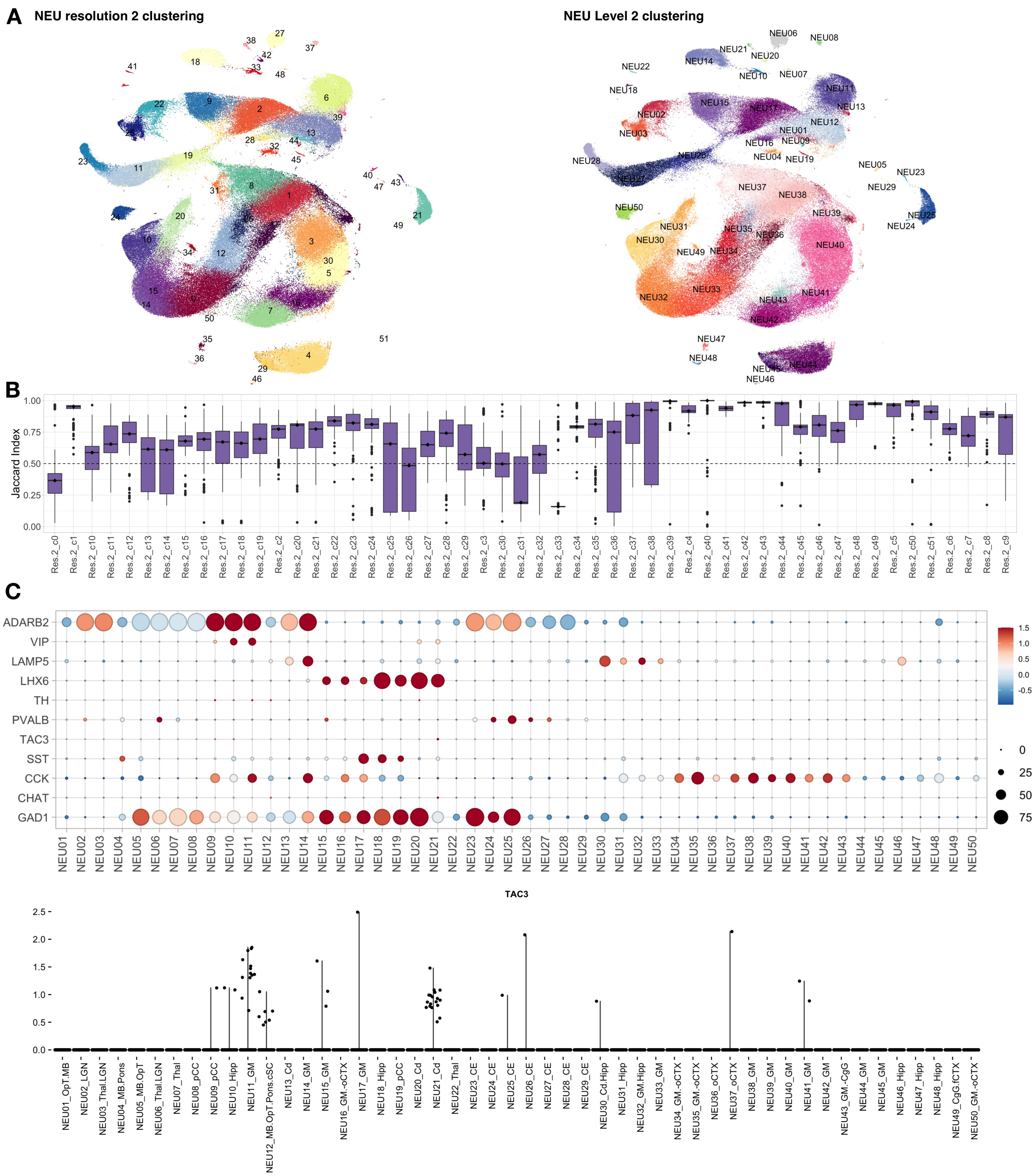


**Figure S12. Related to Figure 1. Neuron clustering stability test and the presence of TAC3+ inhibitory neurons.**

(A) Left, UMAP scatter plot visualization of nuclei colored by clustering at resolution 2 (Methods). Right, UMAP scatter plot visualization of nuclei colored by Level 2 clustering.

(B) Bar plot showing the Jaccard index as a measure of clustering stability (Methods). Total nuclei in the NEU partition were sampled at 90% repeatedly and reclustered over 100 iterations at resolution of 2. Median is annotated (◆).

(C) Top, dot plots showing mean-centered and z-score-scaled expression of marker genes across NEU subclusters. Bottom, dot plot showing the expression of primate specific gens (*TAC3*).

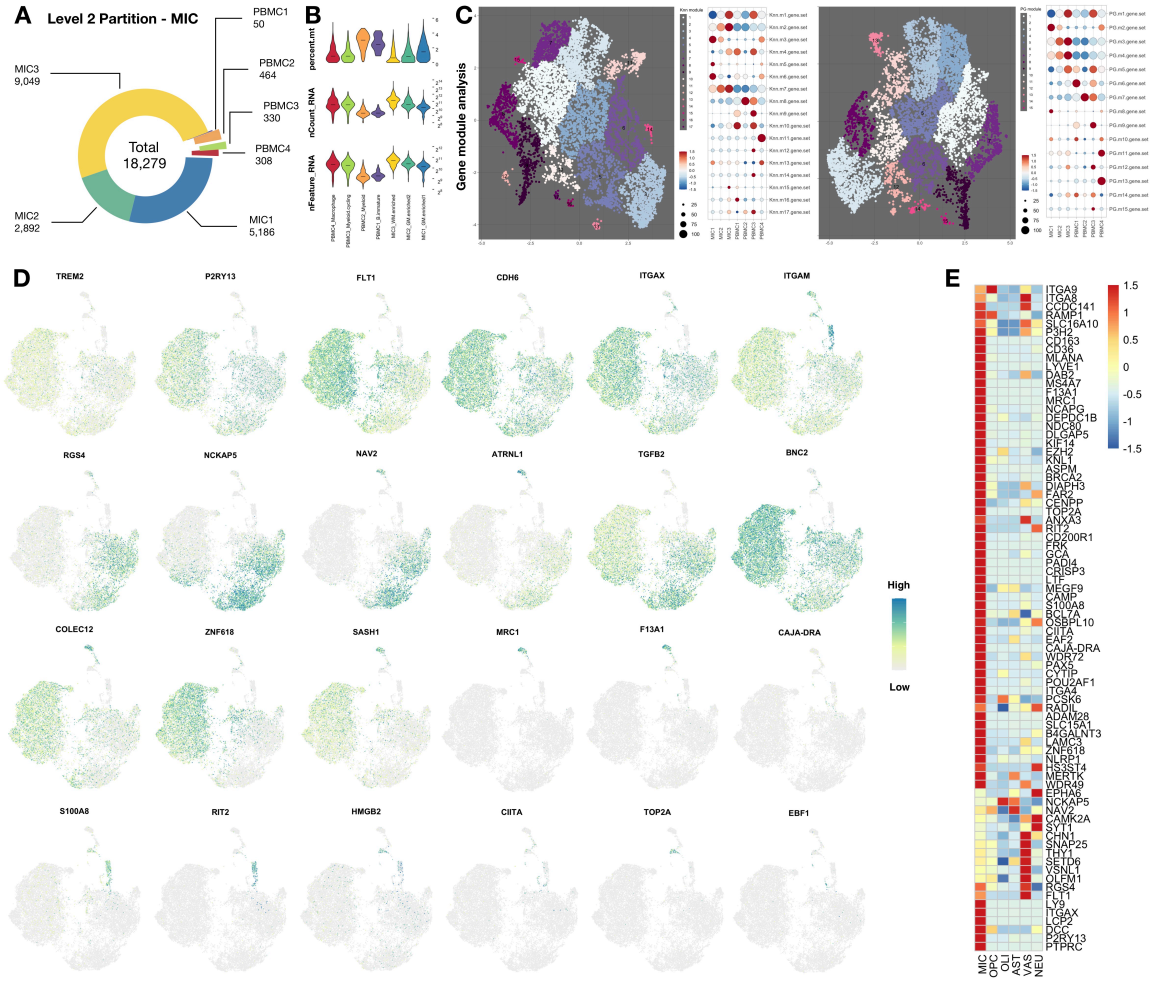


**Figure S13. Related to Figure 2. Nuclei features and gene module analysis in microglia/myeloid lineage cells (MIC).**

(A) Donut plot showing the number of nuclei that passed Level 2 quality control, annotated by subcluster.

(B) Violin plot showing the percentage of RNA reads that mapped to mitochondria genome (percent.mt), the number of RNA detected (nCount_RNA), and the number of RNA species (nFeature_RNA) in each cluster. Median is annotated (–).

(C) UMAP scatter-plot visualization of genes grouped by expression similarity across cells. Genes that passed Moran's *I* statistic spatial test (< 5% FDR) over *k*-nearest neighbor graph (Knn, k=25, left), or over trajectory learned principal graph (PG, right), were identified by Monocle3 graph_test function. Genes are grouped and colored by modules that were identified in each graph test by find_gene_modules function with resolution of 0.001. The list of genes of given module was aggregated through Seurat v3 AddModuleScore function. Dot plot showing the averaged and scaled expression of each module gene set across MIC subclusters.

(D) UMAP scatter plot visualization of MIC nuclei colored by selected marker gene expression.

(E) Heatmap showing averaged and z-score-scaled expression of MIC genes in other Level 1 clusters. The same object (C50 object) was used to make plots shown in Figure 1E.

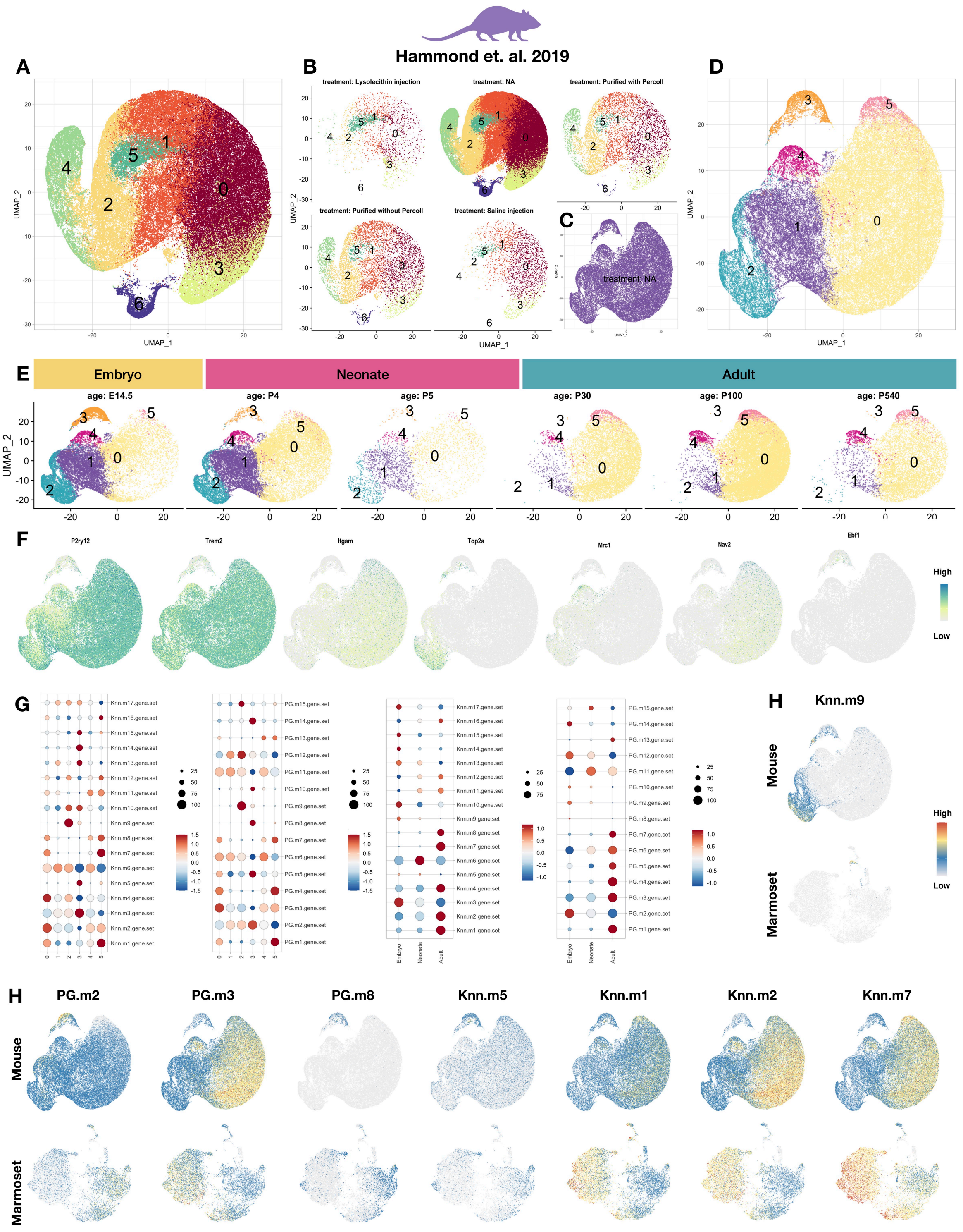


**Figure S14. The expression of gray matter microglia-enriched gene modules found in marmoset are elevated in microglia of young mouse, and the transcriptome of microglia of adult mouse resembles that of marmoset white matter microglia.**

(A) Previously published and deposited data were reanalyzed and reclustered with a pipeline similar to that used in the current study. UMAP scatter-plot visualization of mouse microglia colored by subclusters identified from our pipeline.

(B) UMAP scatter-plot visualization of mouse microglia colored by subclusters identified from our pipeline and split by treatment condition defined in the original publication.

(C) Mouse homeostatic microglia (collected from untreated animals) were selected, reanalyzed, and reclustered with our pipeline.

(D) UMAP scatter-plot visualization of mouse homeostatic microglia colored by subclusters identified from our pipeline.

(E) UMAP scatter-plot visualization of mouse homeostatic microglia colored by subclusters identified from our pipeline and split by the age of the cell as defined in the original publication.

(F) UMAP scatter-plot visualization of mouse microglia colored by selected marker-gene expression.

(G) Dot plot showing the averaged and scaled expression of each module's gene set defined in marmoset microglia across subclusters (left 2 panels) and age (right 2 panels) of mouse homeostatic microglia.

(H) UMAP scatter plot visualization of mouse microglia (top) and marmoset MIC nuclei colored by averaged expression of selected modules found in marmoset.

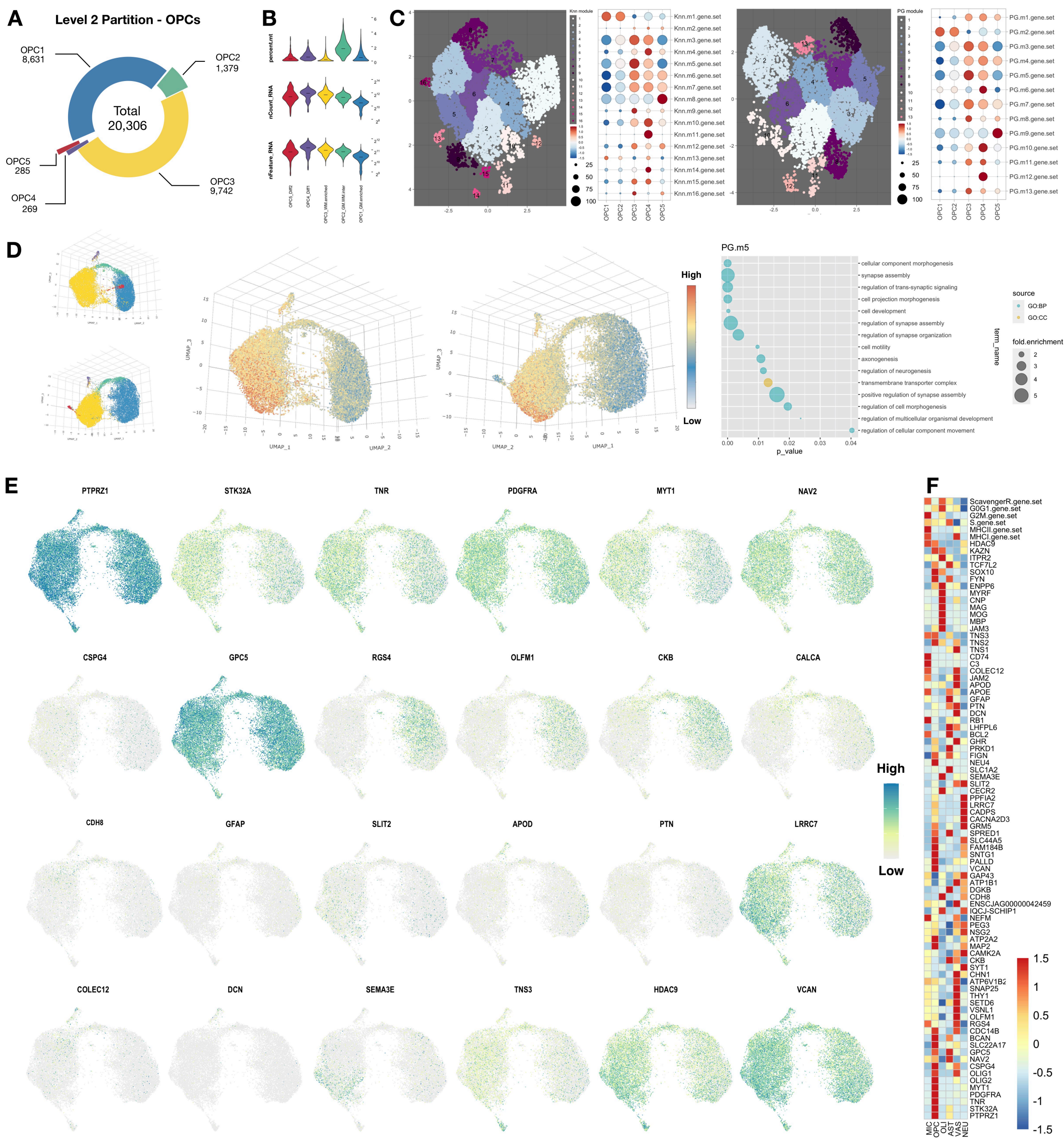


**Figure S15. Related to Figure 3. Assessment of nuclei features in, and gene module analysis on, oligodendrocyte progenitor cells (OPC).**

(A) Donut plot showing the number of nuclei that passed the level 2 quality control, annotated by subcluster.

(B) Violin plot showing the percentage of RNA reads that mapped to mitochondria genome (percent.mt), the number of RNA molecules detected (nCount_RNA), and the number of RNA species (nFeature_RNA) in each cluster. Median is annotated (–).

(C) UMAP scatter-plot visualization of genes grouped by expression similarity across cells. Genes that passed Moran's *I* statistic spatial test (< 5% FDR) over the *k*-nearest neighbor graph (Knn, k=25, left), or over the trajectory learned principal graph (PG, right), were identified by Monocle3 graph_test function. Genes are grouped and colored by modules that were identified in each graph test by find_gene_modules function with resolution of 0.001. The list of genes of each module was aggregated through Seurat v3 AddModuleScore function. Dot plot showing the averaged and scaled expression of each module gene set across OPC subclusters.

(D) Two angles of 3D UMAP scatter-plot visualizations of nuclei colored by OPC subcluster annotation and average expression of selected gene modules. Dot plot showing the enriched GO terms from the list of genes in each module.

(E) UMAP scatter-plot visualization of OPC nuclei colored by selected marker-gene expression.

(F) Heatmap showing the averaged and z-score scaled marker of OPC genes in other Level 1 clusters. The same object (C50 object) was used to make plots shown in Figure 1E.

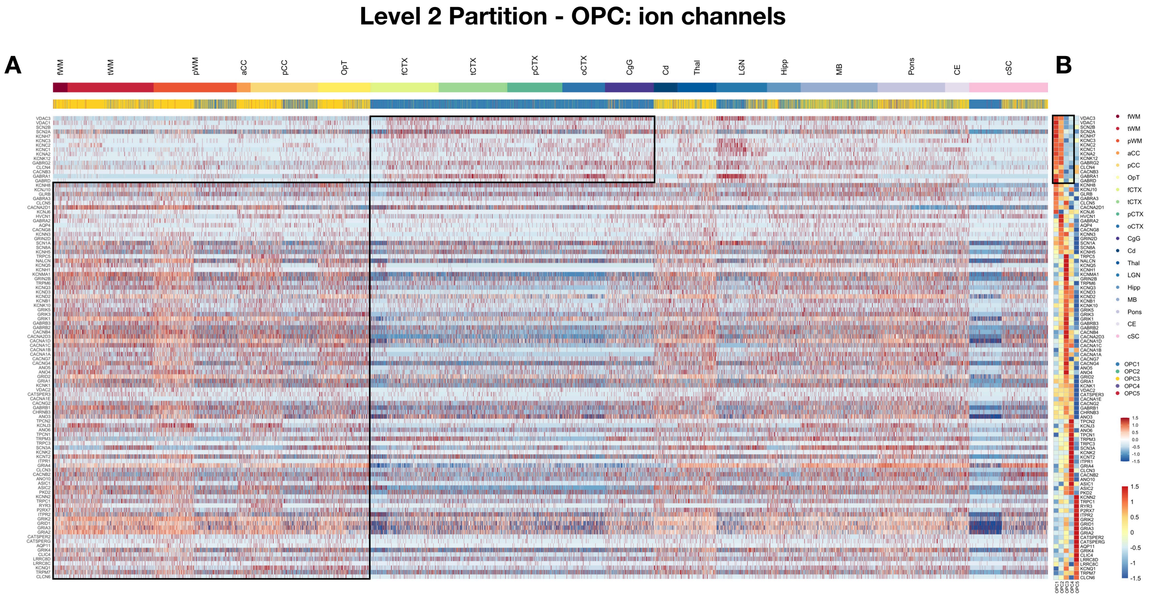


**Figure S16. Related to Figure 3. Oligodendrocyte progenitor cells (OPC) exhibit different ion channel profiles across tissue types.**

(A) Heatmap showing the z-score-scaled gene expression in OPC, with each vertical line representing the expression pattern of one nucleus. The first color bar represents the tissue origin of the OPC, and the second color bar the OPC subcluster annotation. The black boxes highlight two sets of ion channels with markedly different expression levels in white matter (higher for genes in the box on the lower left) and gray matter (higher in the box on the upper right).

(B) Heatmap showing the averaged and z-score-scaled expression in OPC subclusters of the same genes shown

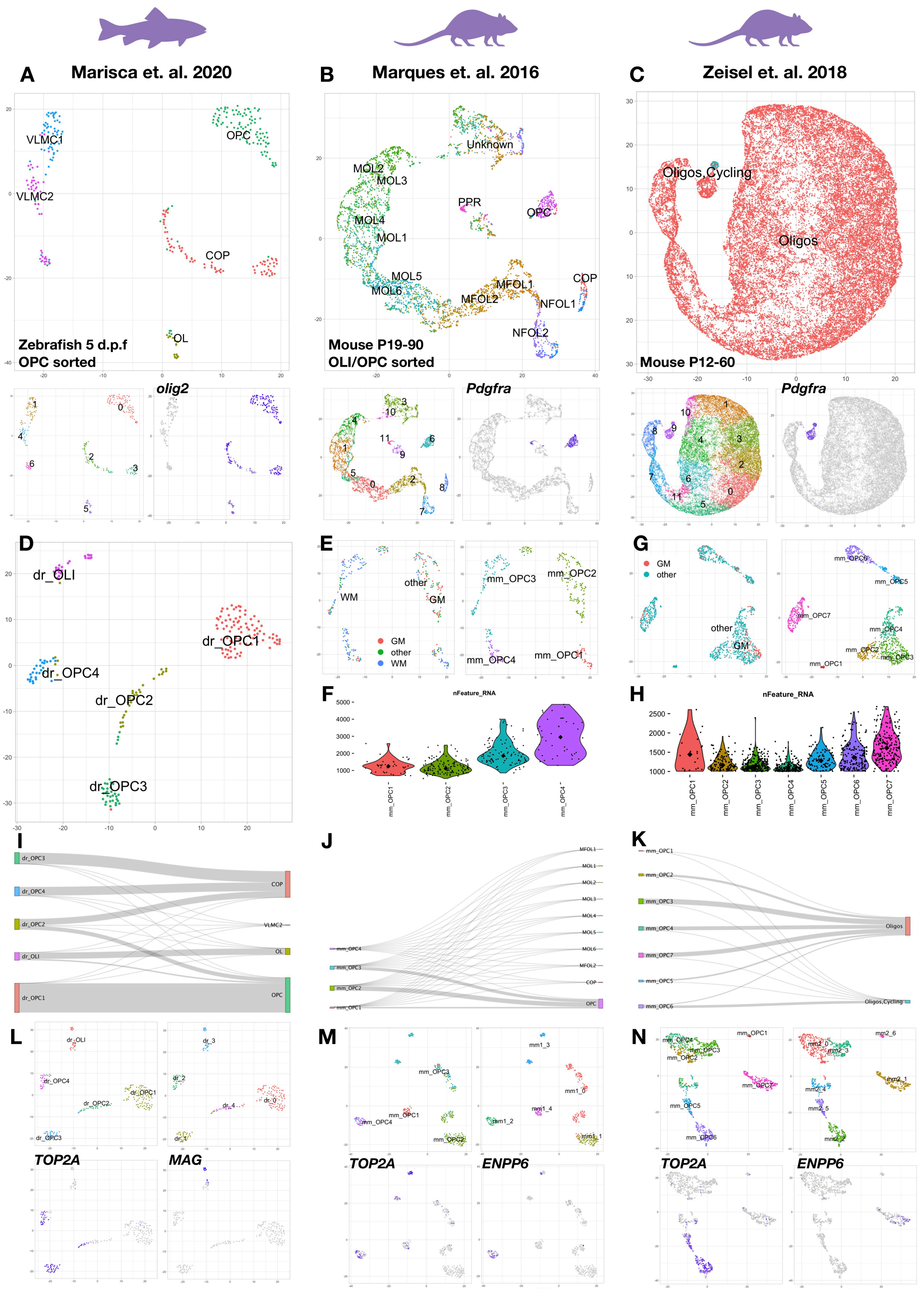


**Figure S17. Related to Figure 3. Reanalysis of zebrafish and mouse derived oligodendrocyte progenitor cells (OPC).**

(A – C) Data deposited from published studies in zebrafish sorted OPC (A), mouse oligodendrocyte lineage cells (B and C) were reanalyzed and reclustered with a pipeline similar to that used in the current study. The original annotation defined by the authors is overlaid onto the UMAP scatter plots in the top row. In the bottom row, for each dataset, the left panel shows our new cluster annotations. The right panel shows the OPC clusters further analyzed in subsequent panels of this figure, specifically clusters 0, 2, 3, 5 (zebrafish; *Olig2*$^{+}$), 5 (mouse, Marques et. al. 2016; *Pdgfra*$^{+}$), and 9 (mouse, Zeisel et. al. 2018; *Pdgfra*$^{+}$).

(D) New UMAP scatter plot showing reclustering of the selected zebrafish OPCs and oligodendrocytes (Clusters 0, 2, 3, 5 in lower-right panel of A) with new labels.

(E) New UMAP scatter plot showing reclustering of the selected mouse OPC clusters (Cluster 5 in lower-right panel of B) colored by new subclusters (right). Using information about the tissue origin provided by the original authors, subclusters were colored and annotated with new labels corresponding to the marmoset OPC subclusters in the current study (tissue in coarse category, left).

(F) Violin plot showing the number of genes detected in each mouse OPC subcluster. Median is annotated (◆).

(G) New UMAP scatter plot showing reclustering of the selected mouse OPC clusters (Cluster 9 in lower-right panel of C) colored by new subclusters (right). Using information about the tissue origin provided by the original authors, subclusters were colored and annotated with new labels corresponding to the marmoset OPC subclusters in the current study (tissue label in coarse category, left).

(H) Violin plot showing the number of genes detected in each mouse OPC subcluster. Median is annotated (◆).

(I – K) Sankey diagrams showing the proportional agreement between new labeling (left) and original labeling (right).

(L – N) New UMAP scatter plots showing the result of reclustering after humanizing gene names of each species with a one-to-one ortholog index based on BioMart information to facilitate cross-species analysis. The identity of cells before and after gene label transfer is mostly preserved. Upper left, UMAP colored by cluster identity as annotated before gene name translation (D, E, and G). Upper right, UMAP colored by cluster identity as annotated after gene name translation (used in Figure 3I). Lower panels, UAMP scatter plot colored by selected marker-gene expression to identify cycling (*TOP2A*$^{+}$) and differentiating (*ENPP6*$^{+}$) OPC.

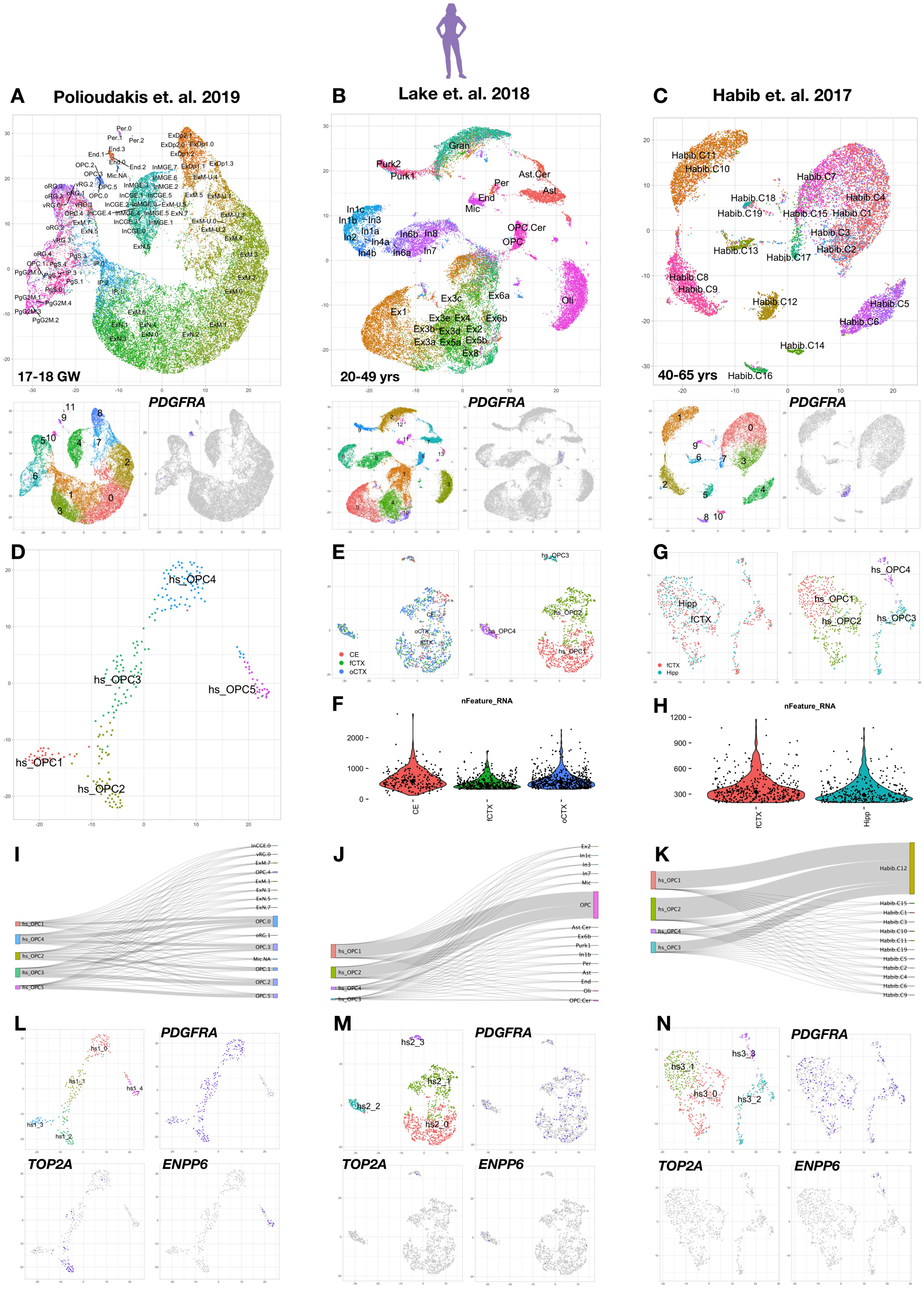


**Figure S18. Related to Figure 3. Reanalysis of human derived oligodendrocyte progenitor cells (OPC).**

(A–C) Data deposited from published studies in human brain were reanalyzed and reclustered with a pipeline similar to that used in the current study. The original annotation defined by the authors is overlaid onto the UMAP scatter plots in the top row. In the bottom row, for each dataset, the left panel shows our new cluster annotations, and the right panel shows the PDG-FRA$^{+}$ OPC clusters further analyzed in subsequent panels of this figure, specifically clusters 9 (A), 6 (B), 5 (C).

(D) New UMAP scatter plot showing reclustering of the selected human OPC from panel A (Cluster 9) with new labels.

(E) New UMAP scatter plot showing reclustering of the selected human OPC from panel B (Cluster 6) colored by new subclusters (right) and tissue type (left).

(F) Violin plot showing the number of genes detected in human OPC grouped by tissue type. Median is annotated (◆).

(G) New UMAP scatter plot showing reclustering of the selected human OPC from panel C (Cluster 5) colored by new subclusters (right) and tissue type (left).

(H) Violin plot showing the number of genes detected in human OPC grouped by tissue type. Median is annotated (◆).

(I – K) Sankey diagrams showing the proportional agreement between new labeling (left) and original labeling (right).

(L – N) Upper left, UMAP colored by cluster identity as used in Figure 3I. Other panels, UMAP scatter plots colored by selected marker-gene expression to identify cycling (*TOP2A*$^{+}$) and differentiating (*ENPP6*$^{+}$) OPC (*PDGFRA*$^{+}$).

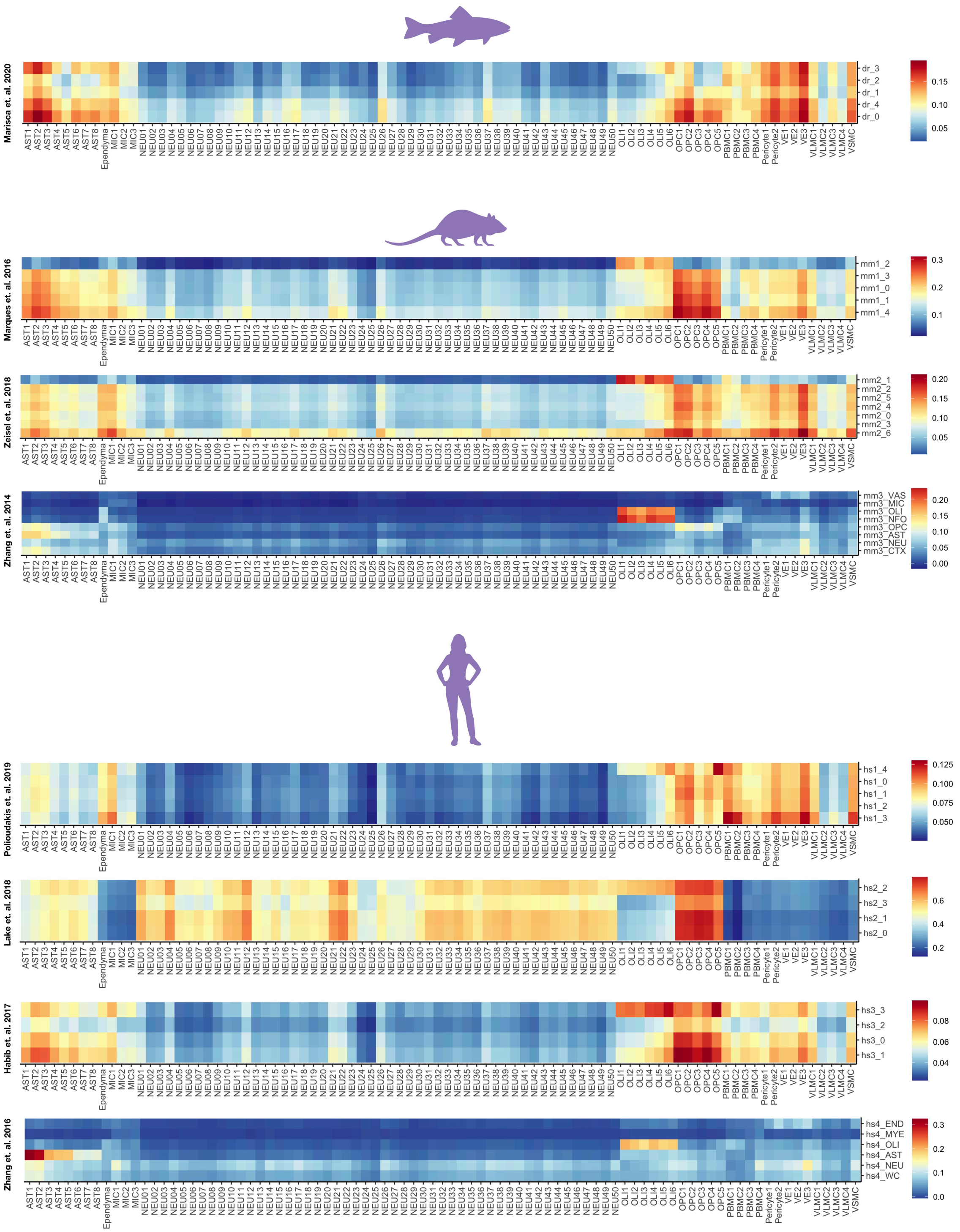


**Figure S19. Related to Figure 3. Cross-cluster comparison of marmoset nuclei to zebrafish, mouse, and human derived oligodendrocyte progenitor cells (OPC).**

Heatmap showing the Pearson's correlation (r) between the 87 marmoset subclusters (C50 object) and newly defined OPC subclusters from zebrafish (*Danio rerio*, dr), mouse (*Mus musculus*, mm), and human (*Homo sapiens*, hs).

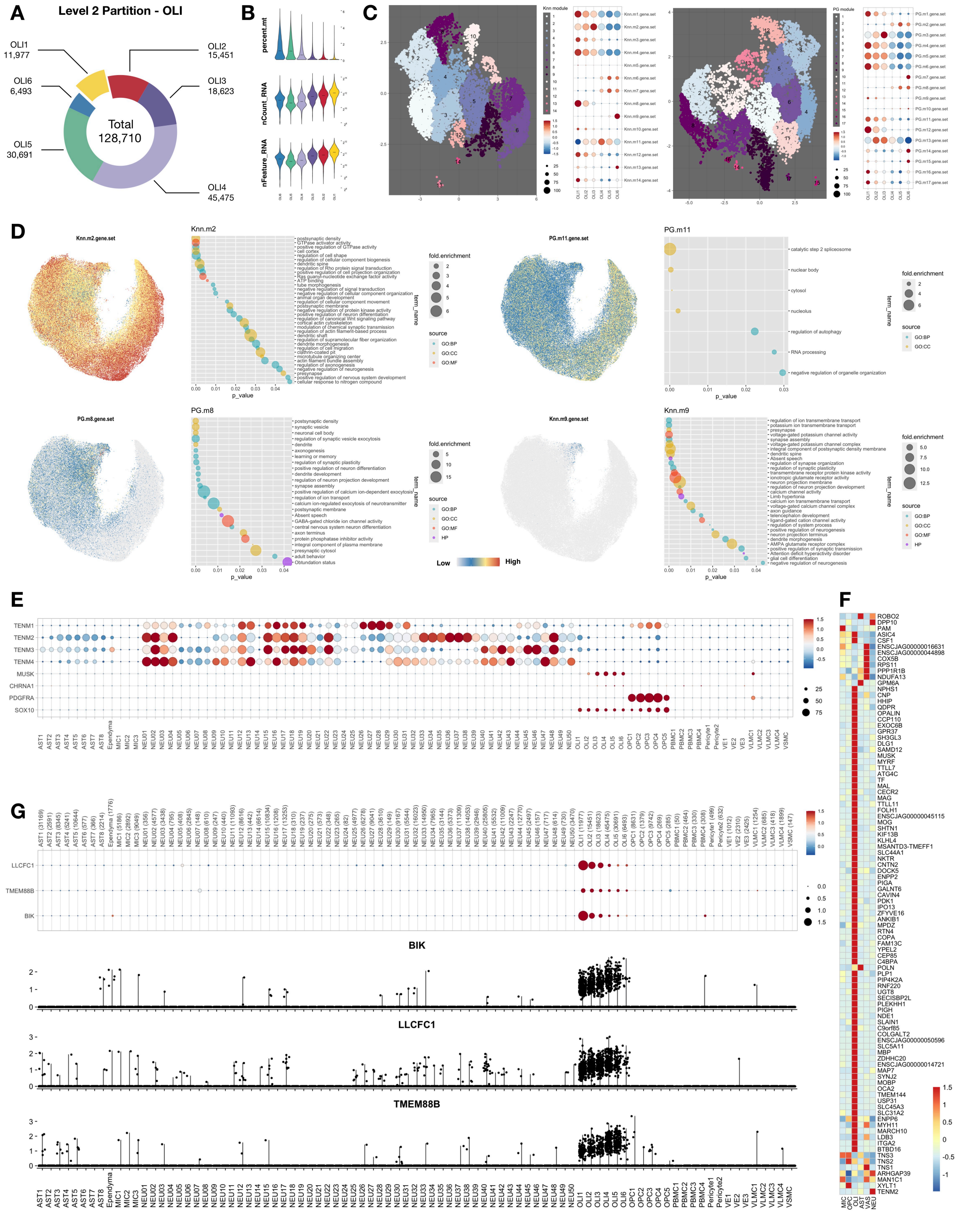


**Figure S20. Related to Figure 4. Nuclei features and gene module analysis in oligodendrocytes (OLI).**

(A) Donut plot showing the number of nuclei that passed Level 2 quality control and were annotated in each subcluster.

(B) Violin plot showing the percentage of RNA reads that mapped to the mitochondria genome (percent.mt), the number of RNA molecules detected (nCount_RNA), and the number of RNA species (nFeature_RNA) in each cluster. Median is annotated (–).

(C) UMAP scatter-plot visualization of genes grouped by expression similarity across cells. Genes that passed Moran's *I* statistic spatial test (< 5% FDR) over the *k*-nearest neighbor graph (Knn, k=25, left), or over the trajectory learned principal graph (PG, right), by Monocle3 graph_test function. Genes are grouped and colored by modules identified in each graph test by find_gene_modules function with resolution of 0.001. The list of genes of each module was aggregated through Seurat v3 AddModuleScore function. Dot plot showing the averaged and scaled expression of each module gene set across OLI subclusters.

(D) UMAP scatter-plot visualization of nuclei colored by averaged expression of each module (color bar is in the center of the bottom row). Dot plot showing the enriched GO terms from the list of genes in each selected module.

(E) Dot plot showing the expression across all 87 subclusters of teneurin family (*TENM1 – 4*), genes associated with muscle weakness (*MUSK*, *CHRNA1*), and oligodendrocyte lineage/OPC genes (*SOX10*, *PDGFRA*).

(F) Heatmap showing the mean-centered and z-score-scaled OLI marker-gene expression across Level 1 classes.

(G) Dot plot showing the expression of rare and understudied genes (*BIK*, *TMEM88B*, and *LLCFC1*) enriched in OLI.

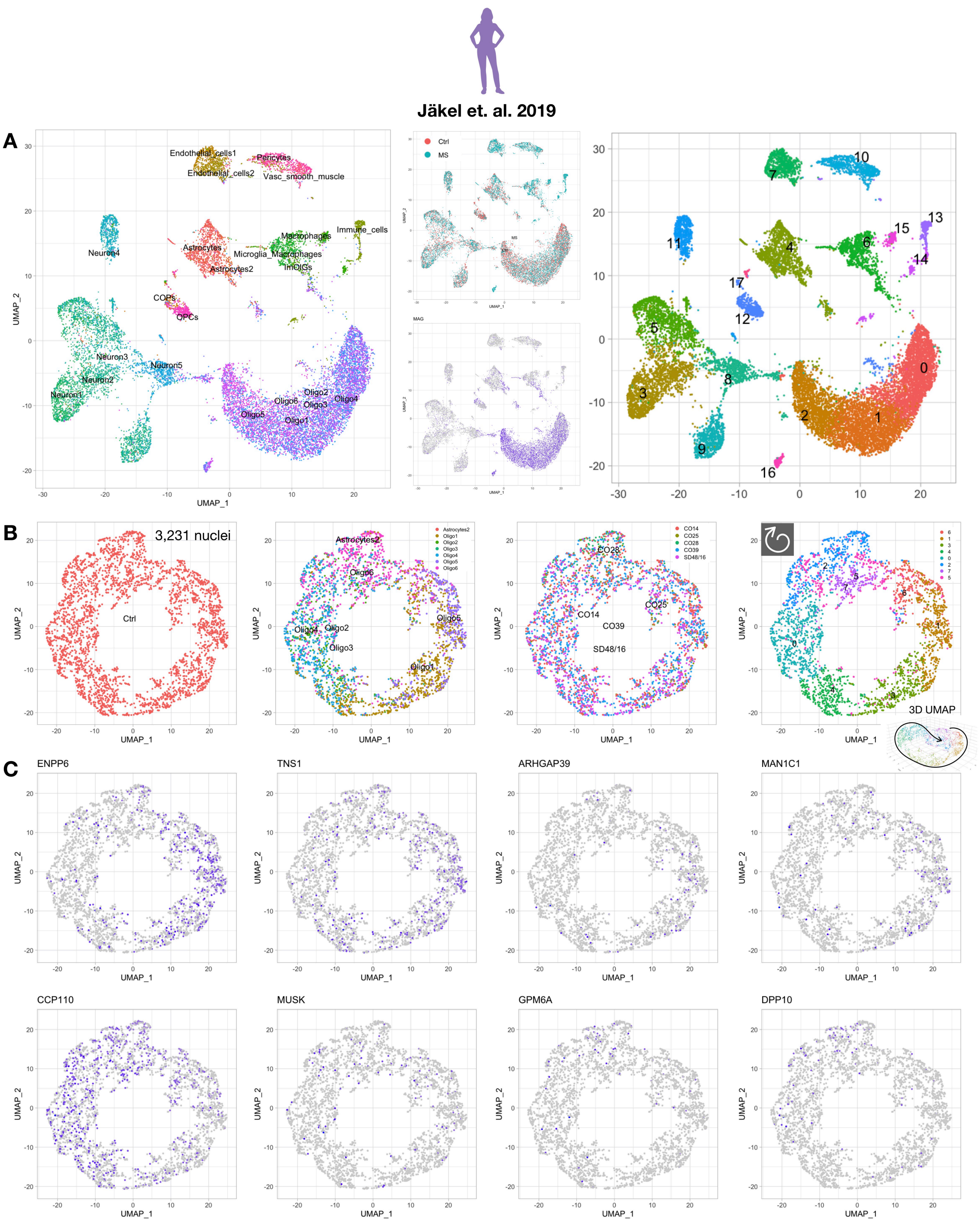


**Figure S21. Related to Figure 4. Adult human oligodendrocytes are arranged into a spiral pattern in UMAP space.**

(A) Previously published and deposited data were reanalyzed and reclustered with a pipeline similar to that used in this current study. UMAP scatter-plot visualization of cells colored by the original annotations defined by the authors (left: cell type; middle top: tissue source, control [Ctrl] or multiple sclerosis [MS]). UMAP plot of human oligodendrocytes colored by *MAG* expression to facilitate subset selection (middle bottom). UMAP plot colored by newly defined cluster number (right).

(B) Newly defined clusters 0, 1, 2 from control samples (excluding samples from MS patients) were selected for further subset analysis. UMAP scatter-plot visualization of cells colored by the original annotations defined by the authors (first three panels) and newly defined cluster numbers (right-most panel). 3D UMAP scatter-plot visualization of the nuclei colored by the newly defined cluster numbers (right bottom).

(C) UMAP scatter-plot visualization of human oligodendrocyte lineage cells colored by selected marker-gene expression.

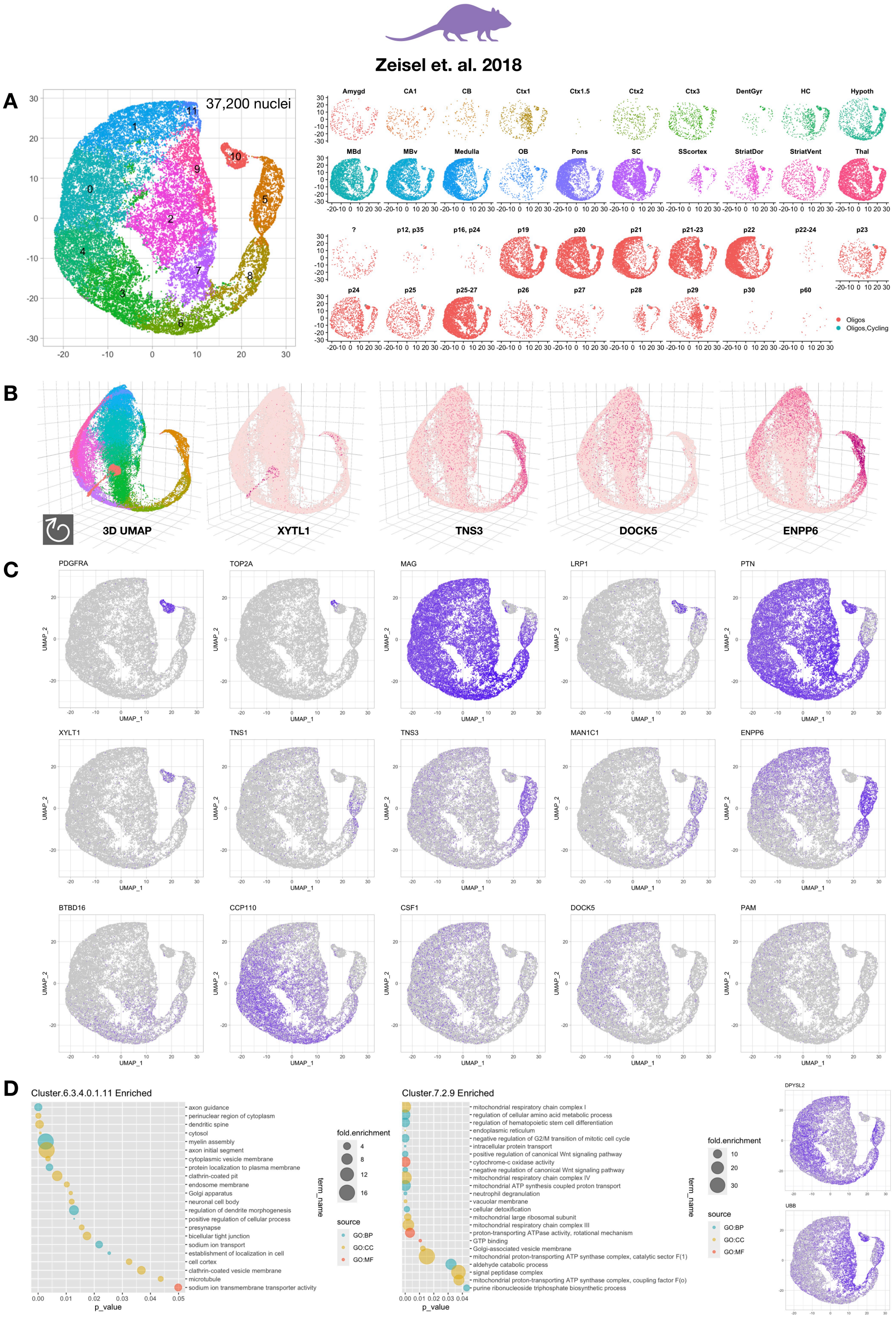


**Figure S22. Related to Figure 4. Mouse oligodendrocyte lineage cells are arranged into a spiral pattern in UMAP space along a putative differentiation trajectory.**

(A) Previously published and deposited data were reanalyzed and reclustered with a pipeline similar to that used in this current study. UMAP scatter-plot visualization of cells colored by newly defined cluster numbers (left). The annotations defined by the original authors, such as sampling source (top right) and age (bottom right), were overlaid onto the UMAP scatter plots.

(B) 3D UMAP scatter-plot visualization of mouse nuclei colored by the newly defined cluster numbers.

(C) UMAP scatter-plot visualization of mouse oligodendrocyte lineage cells colored by selected marker-gene expression.

(D) Dot plot showing the enriched GO terms from the list of differentially expressed genes between left (combining mouse OLI subclusters 6, 3, 4, 0, 1, 11) and right (combining subclusters 7, 2, 9) branches. Left, GO terms related to genes increased in the left branch, including pathways required for myelin formation and neural cell interaction. Middle, GO terms related to genes increased in the right branch, including pathways related to mitochondrial and metabolic functions. Right, UMAP scatter-plot visualization of oligodendrocyte lineage cells colored by markers enriched in each branch. *UBB* expression is elevated in the subset of oligodendrocyte lineage cells (right branch), indicating active protein turnover.

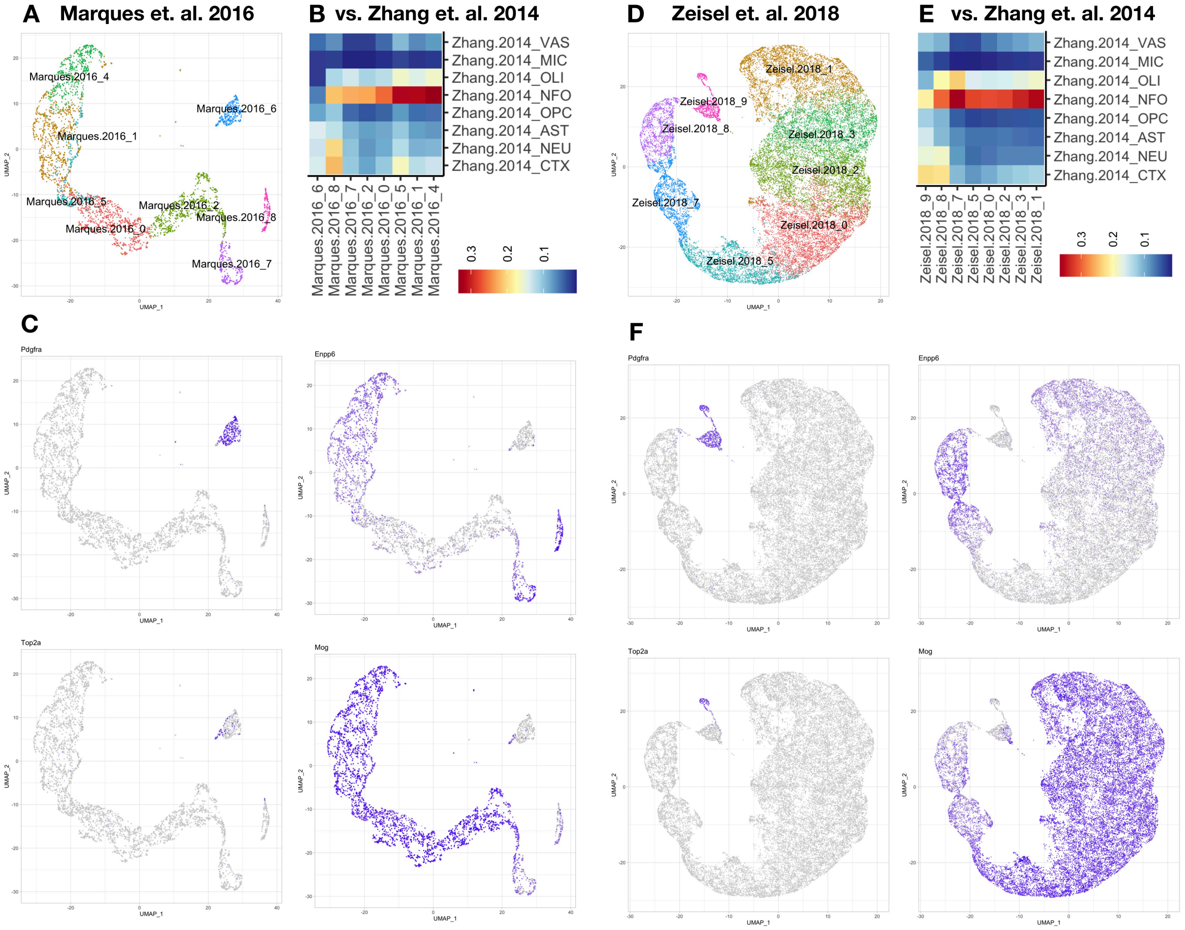


**Figure S23. Related to Figure 4. Cross-dataset comparison of mouse oligodendrocyte lineage cells.**

(A) Previously published and deposited data were reanalyzed and reclustered with a pipeline similar to that used in this current study. UMAP scatter-plot visualization of cells colored by newly defined cluster numbers. Clusters are reordered from young to old (Marques.2016_6, 8, 7, 2, 0, 5, 2, 4) along the putative trajectory of oligodendrocyte lineage cell differentiation.

(B) Heatmap showing the Pearson's correlation (r) of transcriptome between 8 subclusters of oligodendrocyte lineage cells from Marques et. al. 2016 and 8 types of immunopanned cells from Zhang et. al 2014 (VAS, endothelial cell; MIC, immune cell; OLI, myelinating oligodendrocyte; NFO, newly formed oligodendrocyte; OPC, oligodendrocyte progenitor cell; AST, astrocyte; NEU, neuron; CTX, unpurified total cell from cortex). NFO identified by immunopanning (Zhang.2014_NFO) correlate highest with the putative "older" oligodendrocyte (Marques.2016_5, 1, 4) identified by single cell analysis.

(C) UMAP scatter-plot visualization of mouse oligodendrocyte lineage cells from Marques et. al. 2016 colored by *Pdgfra*, *Enpp6*, *Top2a*, and *Mag* expression.

(D) Previously published and deposited data were reanalyzed and reclustered with a pipeline similar to that used in this current study. Putative stressed oligodendrocytes ($UBB^{high}$) were removed from previous analysis in Figure S22D and reclustered. UMAP scatter-plot visualization of cells colored by newly defined cluster numbers. Clusters are reordered from young to old (Zeisel.2018_9, 8, 7, 5, 0, 2, 3, 1) along the putative trajectory of oligodendrocyte lineage cell differentiation.

(E) Heatmap showing the Pearson's correlation (r) of transcriptome between 8 subclusters of oligodendrocyte lineage cells from Zeisel et. al. 2018 and 8 types of immuno-panned cells from Zhang et. al 2014. NFO identified by immunopanning (Zhang.2014_NFO) highly correlate with both putative "older" (Zeisel.2018_1) and "younger" (Zeisel.2018_7) oligodendrocyte identified by single cell analysis.

(F) UMAP scatter-plot visualization of mouse oligodendrocyte lineage cells from Zeisel et. al. 2018 colored by *Pdgfra*, *Enpp6*, *Top2a*, and *Mag* expression.

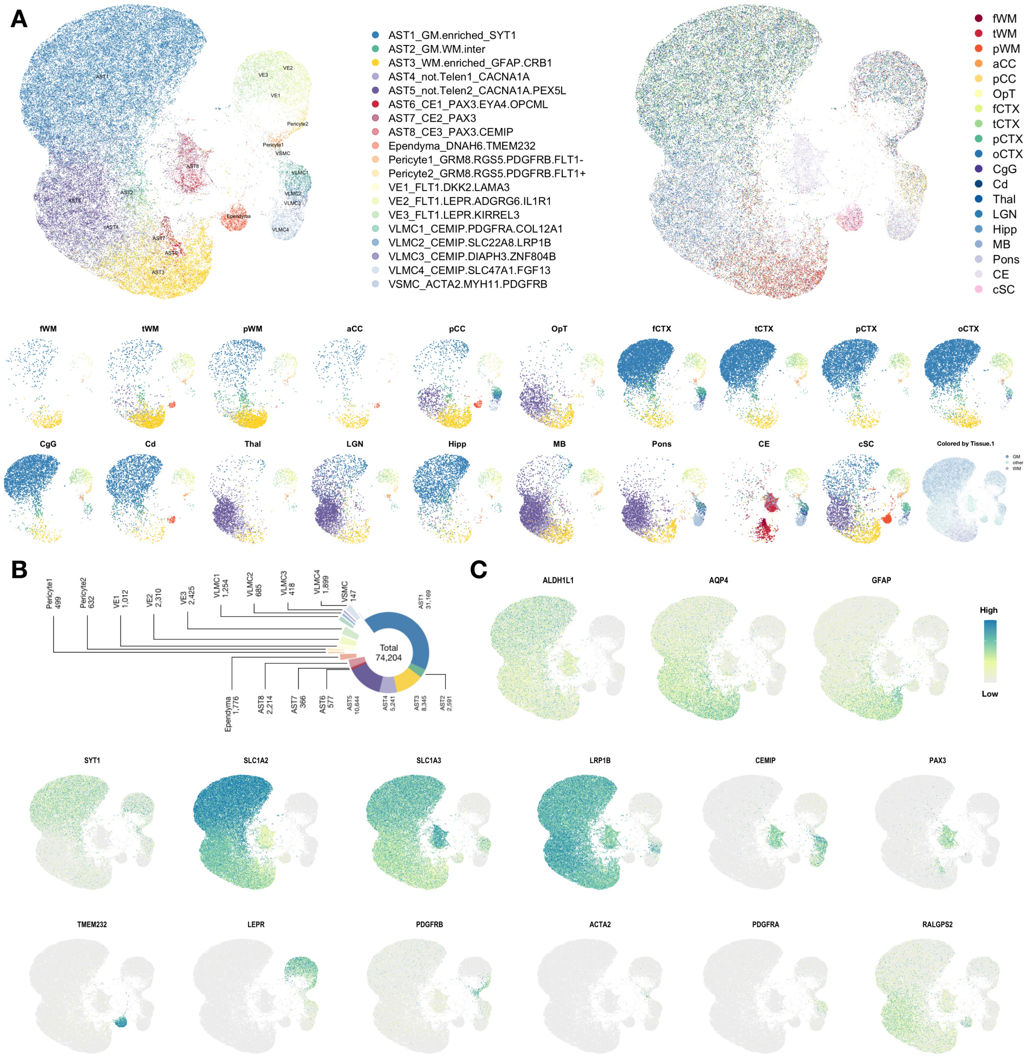


**Figure S24. Related to Figure 5. Nuclei features in cells at the barriers of the CNS.**

(A) UMAP scatter-plot visualization of combined astrocyte/vascular (AST/VAS) nuclei, colored by subcluster (upper left) and sampling site (upper right). UMAP scatter-plot visualization of AST/VAS subclusters split by sampling site and coarse tissue type (bottom).

(B) Donut plot showing the number of AST/VAS nuclei that passed Level 2 quality control and were annotated into subclusters.

(C) UMAP scatter-plot visualization of AST/VAS nuclei colored by selected marker gene expression.

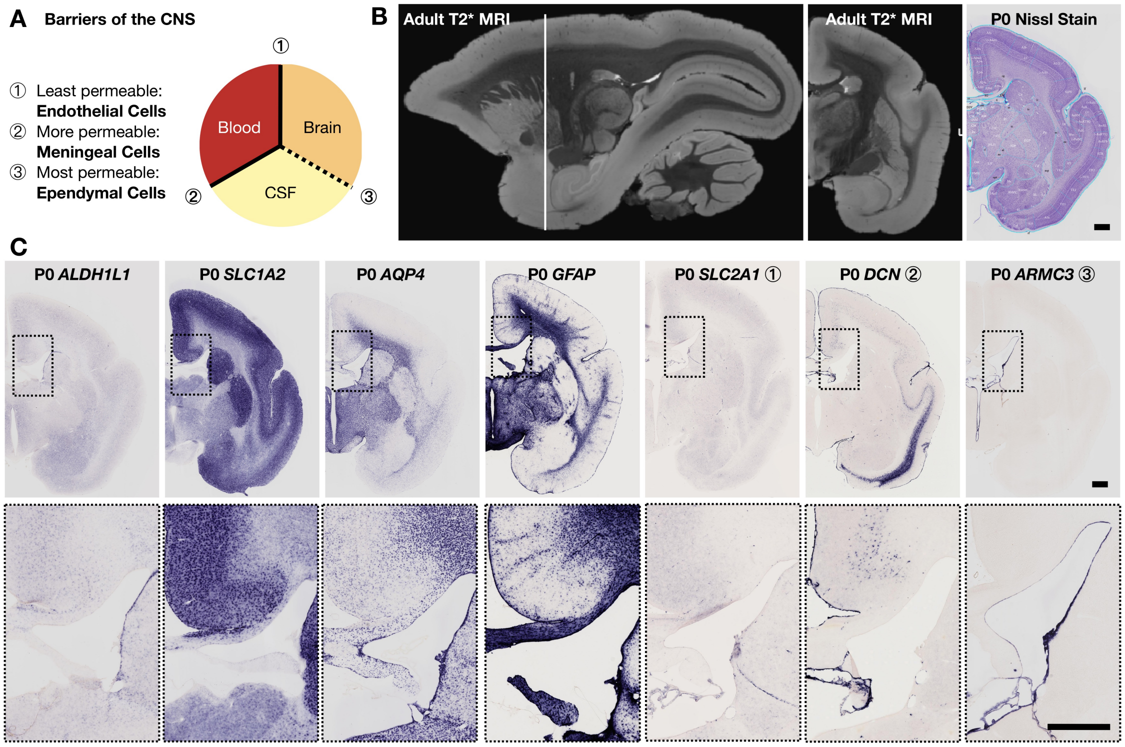


**Figure S25. Related to Figure 5. Cell types and their gene expression patterns at the barriers of the CNS.**

(A) Schematic illustration of the barriers of the CNS. The least permeable blood-brain barrier (BBB, ①) is formed mostly by vascular endothelial cells, the somewhat more permeable blood-CSF barrier (②) by meningeal cells, and the most permeable CSF-brain barrier (ventricle, ③) by ependymal cells.

(B) Approximate cross-sectional plane (white line) indicated on a sagittal view of an adult marmoset postmortem T2*-weighted MRI (left). Coronal view of the same adult marmoset T2*-weighted MRI (middle). Coronal histological section of a P0 marmoset stained with Nissl's method (right) and anatomically matched to the adult coronal T2*-weighted MRI slice.

(C) Selected in situ hybridization (ISH) of genes enriched in endothelial, meningeal, and ependymal cells. Enlarged areas are indicated by the box with dashed edge.

Scale bar, 1mm. MRI images were acquired from the Marmoset Brain Mapping database. Nissl stain and ISH images were acquired from the Marmoset Gene Atlas.

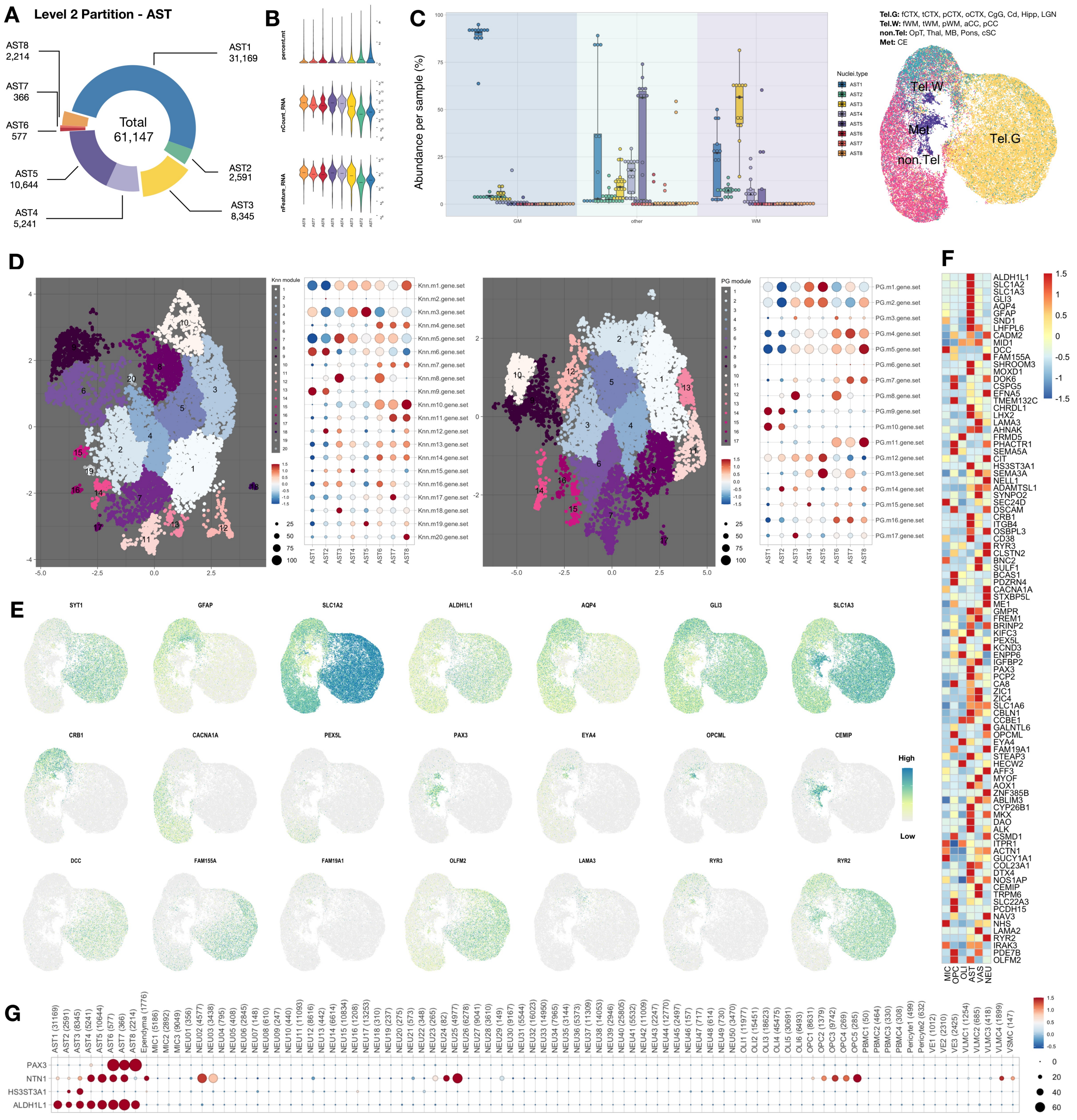


### Figure S26. Related to Figure 5. Assessment of nuclei features and gene module analysis of astrocytes (AST).

(A) Donut plot showing the number of nuclei that passed the Level 2 quality control and were annotated into AST subclusters.

(B) Violin plot showing the percentage of RNA reads that mapped to mitochondria genome (percent.mt), the number of RNA molecules detected (nCount_RNA), and the number of RNA species (nFeature_RNA) in each cluster. Median is annotated (–).

(C) Left, box plot showing the relative abundance of AST subclusters in each coarse sampling category (IL06_tissue.1 category; see Figure S1 legend for full list). AST1 is enriched in cortical gray matter (GM) and AST3 in cerebral white matter (WM). Median is annotated (◆). Right, UMAP scatter-plot visualization of AST partition colored by tissue types. **Tel.G**, telencephalon gray; **Tel.W**, telencephalon white; **non.Tel**, non-telencephalon; **Met**, metencephalon.

(D) UMAP scatter-plot visualization of genes grouped by expression similarity across cells. Genes that passed Moran's *I* statistic spatial test (< 5% FDR) over the *k*-nearest neighbor graph (Knn, k=25, left), or over the trajectory learned principal graph (PG, right), by Monocle3 graph_test function. Genes are grouped and colored by modules identified in each graph test by find_gene_modules function with resolution of 0.001. The list of genes in each module was aggregated through Seurat v3 AddModuleScore function. Dot plot showing the averaged and scaled expression of each module's gene set across AST subclusters.

(E) UMAP scatter-plot visualization of astrocytes colored by selected marker gene expression.

(F) Heatmap showing mean-centered and z-score-scaled AST marker gene expression across Level 1 classes.

(G) Dot plot showing selected mean-centered and z-score-scaled expression of selected subcluster-defining AST marker genes.

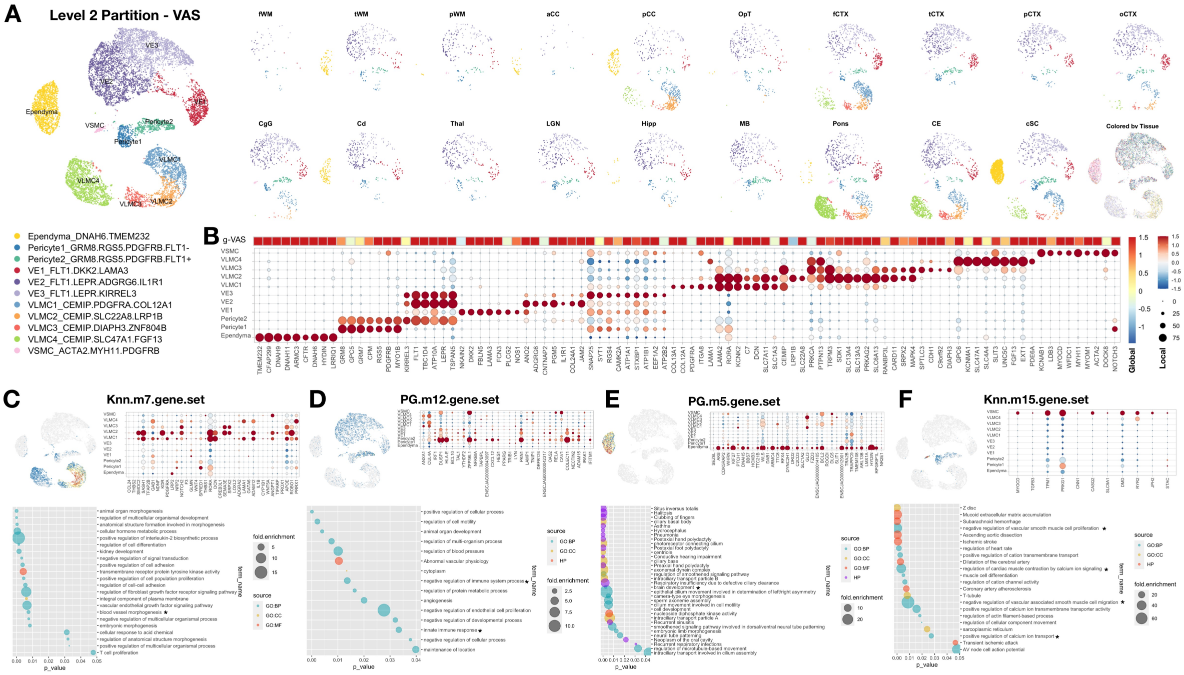


**Figure S27. Related to Figure 5. Nuclei features, gene expression, and gene modules in vascular and other cells at the CNS barrier cells (VAS).**

(A) Local UMAP scatter-plot visualization of other cells at the CNS barrier (mostly from the VAS class in Level 1 analysis) colored by subcluster identity and split by sampling site.

(B) Heatmap and dot plot showing mean-centered and z-score-scaled marker gene expression in global space (heatmap, VAS average expression among Level 1 classes, C50) and local space (dot plot, VAS subclusters). The portion of the heatmap that represents global VAS (g-VAS) is reproduced here from the full heatmap shown in Figure S28G. Dot size indicates the percentage of nuclei that expressed each gene in each subcluster. Scaling is relative to expression across all VAS nuclei in which a given gene was detected.

(C – F) Upper left, UMAP scatter-plot visualization of VAS nuclei colored by averaged expression of different modules. Bottom, dot plot showing the enriched Gene Ontology (GO) terms from the list of genes in each module; selected terms for marker expression plot are annotated (*). Upper right, dot plot showing the detected genes from selected terms. Due to their relevance for brain structure and function, terms such as “blood vessel morphogenesis” (C), “innate immune response” (D), “brain development” (E) “positive regulation of calcium transport” (F) were selected for gene expression dot plot.

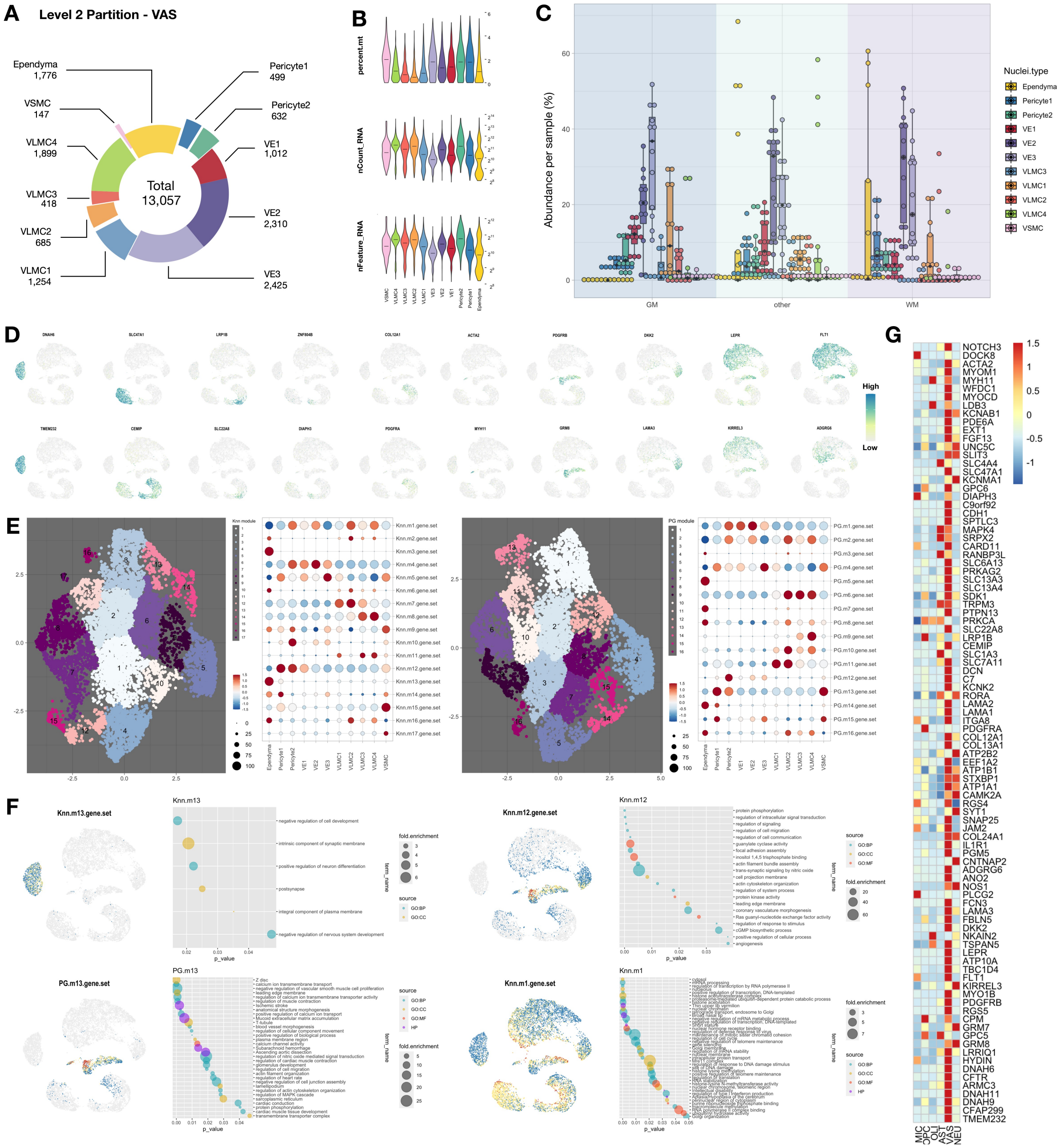


### Figure S28. Related to Figure 5. Further assessment of nuclei features and gene module analysis of vascular and other cells at the CNS barriers (VAS).

(A) Donut plot showing the number of nuclei that passed Level 2 quality control and were annotated into subclusters.

(B) Violin plot showing the percentage of RNA reads that mapped to mitochondria genome (percent.mt), the number of RNA molecules detected (nCount_RNA), and the number of RNA species (nFeature_RNA) in each cluster. Median is annotated (–).

(C) Box plot showing the relative abundance of VAS subclusters in each coarse sample (IL06_tissue.1, see Figure S1 legend for full list). Median is annotated (◆).

(D) UMAP scatter-plot visualization of VAS nuclei colored by selected marker-gene expression.

(E) UMAP scatter plot visualization of genes grouped by expression similarity across cells. Genes that passed the Moran's *I* statistic spatial test (< 5% FDR) over *k*-nearest neighbor graph (Knn, k=25, left), or the over trajectory learned principal graph (PG, right), by Monocle3 graph_test function. Genes are grouped and colored by modules identified in each graph test by find_gene_modules function with resolution of 0.001. The list of genes in each module was aggregated through Seurat v3 AddModuleScore function. Dot plot showing the averaged and scaled expression of each module's gene set across VAS subclusters.

(F) UMAP scatter-plot visualization of VAS nuclei colored by averaged expression of selected modules enriched in a subset of VAS subclusters.

(G) Heatmap showing mean-centered and z-score-scaled VAS marker gene expression across Level 1 clusters.

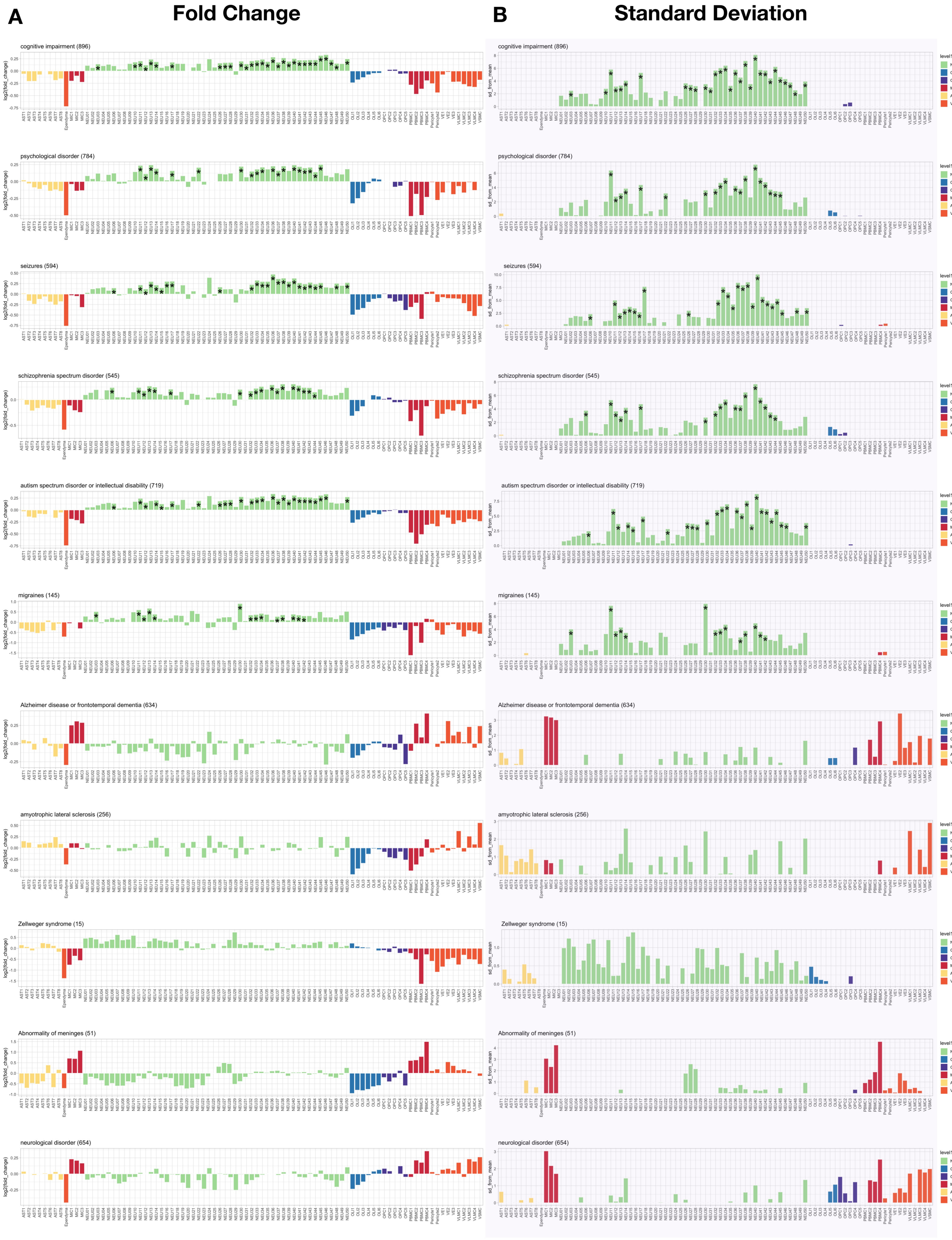
A
Fold Change
B
Standard Deviation
cognitive impairment (896)
psychological disorder (784)
seizures (594)
schizophrenia spectrum disorder (545)
autism spectrum disorder or intellectual disability (719)
migraines (145)
Alzheimer disease or frontotemporal dementia (634)
amyotrophic lateral sclerosis (256)
Zellweger syndrome (15)
Abnormality of meninges (51)
neurological disorder (654)
log2(fold_change)
sd_from_mean
level1.anno
NEU
OLI
OPC
MIC
AST
VAS

Continue

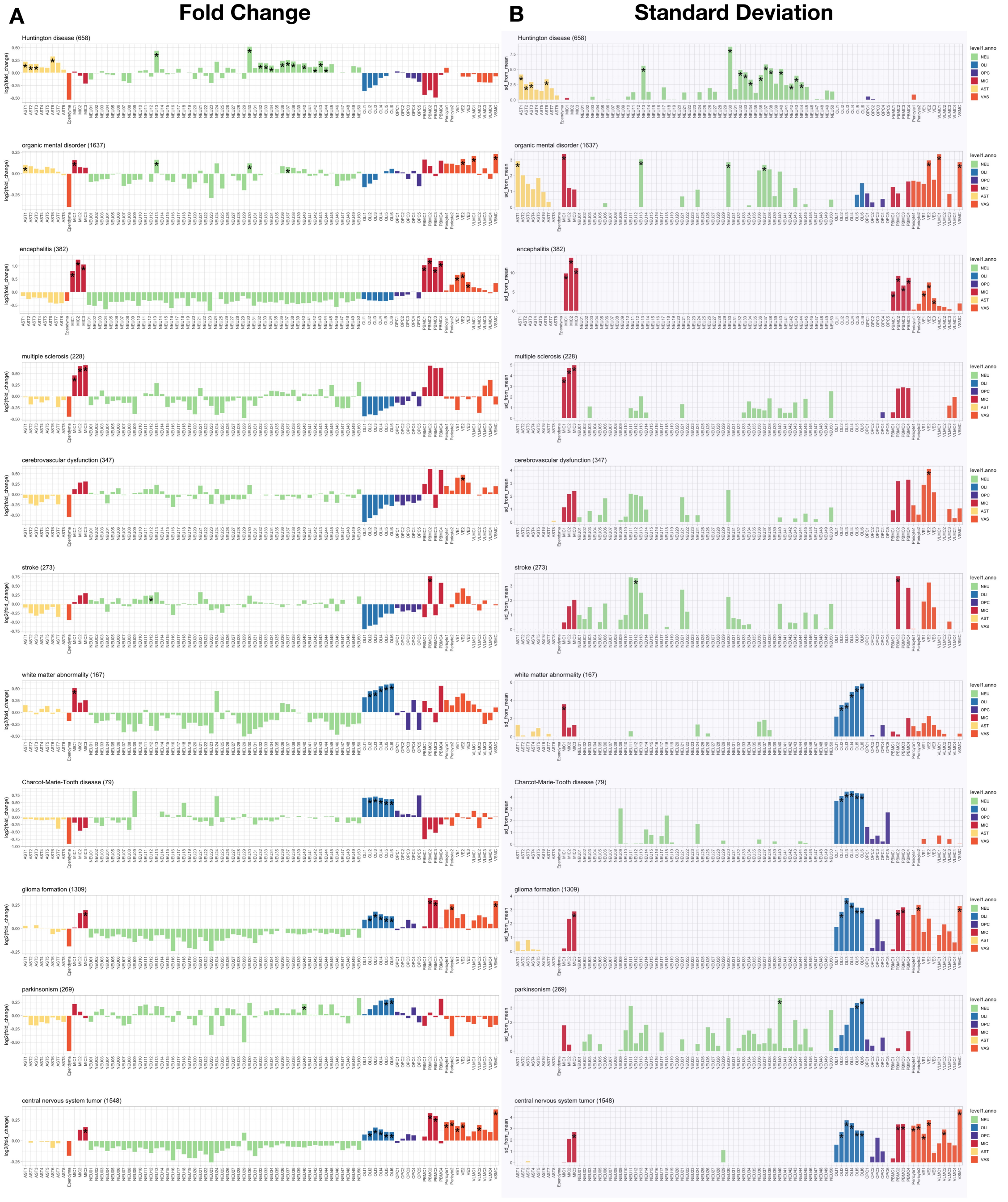


**Figure S29. Related to Figure 6. Cellular contributions to selected human neurological disorders**

(A) Bar graph showing cell type-specific enrichment of genes annotated in the Ingenuity Pathway Analysis database as being associated with cognitive impairment, psychological disorder, seizures, schizophrenia spectrum disorder, autism spectrum disorder or intellectual disability, migraine, Alzheimer disease or frontotemporal dementia, Zellweger syndrome, abnormality of meninges, neurological disorder, Huntington disease, organic mental disorder, encephalitis, multiple sclerosis, cerebrovascular dysfunction, stroke, white matter abnormality, Charcot-Marie-Tooth disease, glioma formation, parkinsonism, and central nervous system tumor. Statistically significant cell types are labeled (Benjamini-Hochberg corrected *$p < 0.05$).

(B) Bar graph showing the standard deviation of the enrichment probability for neurological disorder-associated genes calculated by Expression-Weighted Cell-type Enrichment (EWCE) analysis, after bootstrapping. Bar color represents the Level 1 class of origin.

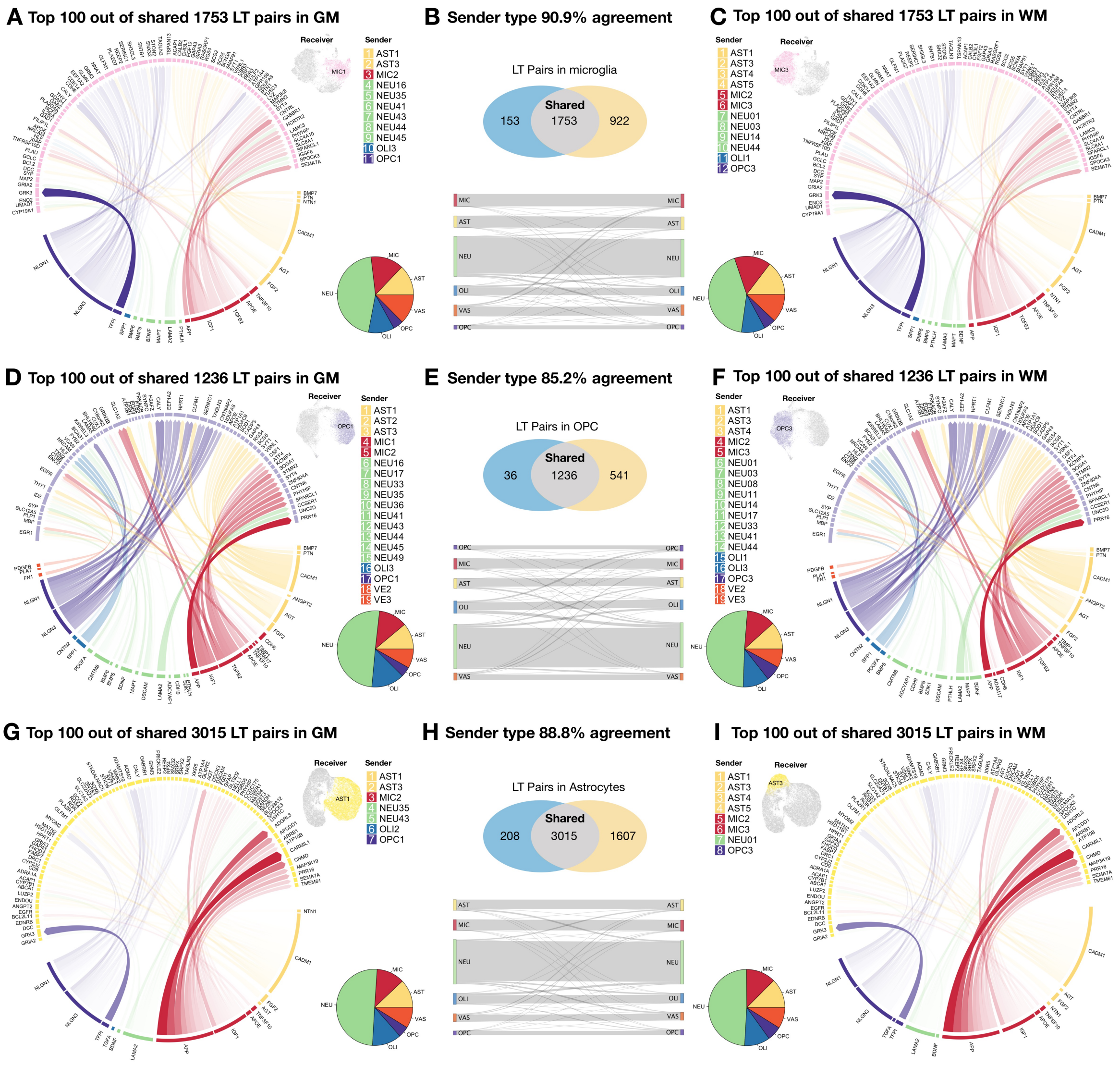


**Figure S30. Related to Figure 7. Mapping intercellular communication in cerebral white matter (WM) and cortical gray matter (GM).**

(A – I) Shared ligand-target (LT) pairs in GM and WM for MIC (A – C), OPC (D – F), and AST (G – I). Circos plots showing the intercellular ligand-target pairs. Top 100 unique ligand-target pairs in GM. Pie chart showing the sender contribution of shared ligand-target pairs grouped by Level 1 class in GM (A, D, and G). Circos plots showing the intercellular ligand-target pairs. Top 100 unique ligand-target pairs in WM. Pie chart showing the sender contribution of shared ligand-target pairs grouped by Level 1 class in WM (C, F, and I).

(B, E, and H) Top, the number of final established ligand-target (LT) pairs in GM or WM are compared in Venn diagrams. Bottom, Sankey diagram showing the agreement of sender type between GM and WM.

**Data S1. Sample attributes.**

**Data S2. List of genes in each module.**

**Data S3. NicheNet LRT analysis results.**

**Data S4. List of genes in each plot.**
